# The Origins and Drivers of Neotropical Diversity

**DOI:** 10.1101/2021.02.24.432517

**Authors:** Andrea S. Meseguer, Alice Michel, Pierre-Henri Fabre, Oscar A. Pérez-Escobar, Guillaume Chomicki, Ricarda Riina, Alexandre Antonelli, Pierre-Olivier Antoine, Frédéric Delsuc, Fabien L. Condamine

**Author notes:** Corresponding author: Andrea Sánchez Meseguer: Real Jardín Botánico de Madrid (RJB-CSIC), Plaza de Murillo 2, 28014 Madrid, Spain; 0034-914203017 (Ext. 144). ***Statement of authorship:*** A.S.M., F.D. and F.L.C designed research; A.S.M., A.M., P.-H.F., R.R., and F.L.C. performed research; A.S.M., A.M., and F.L.C analysed data; A.S.M. wrote the paper with the input of F.L.C. and contributions of all authors.

## Abstract

The origins and evolution of the outstanding Neotropical biodiversity are still debated. A comprehensive understanding is hindered by the lack of deep-time comparative data across wide phylogenetic and ecological contexts. Here, we evaluate four evolutionary scenarios assuming different diversification trajectories and drivers of Neotropical diversification, and assess their variation across Neotropical regions and taxa. Our analysis of 150 phylogenies (12,512 species) of seed plants and tetrapods reveals that Neotropical diversity has mostly expanded through time (70% of the clades), while scenarios of saturated and declining diversity also account for 21% and 9% of Neotropical diversity, respectively. We identify five biogeographic areas that represent distinctive units of long-term Neotropical evolution (Pan-Amazonia, Dry Diagonal, Bahama-Antilles, Galapagos, and an ‘elsewhere’ region) and find that diversity dynamics do not differ across these areas, suggesting no geographic structure in long-term Neotropical diversification. In contrast, diversification dynamics differ substantially across taxa: plant diversity mostly expanded through time (88%), while a substantial fraction (43%) of tetrapod diversity accumulated at a slower pace or declined toward the present. These opposite evolutionary patterns may reflect different capacities for plants and tetrapods to cope with climate change, with potential implications for future adaptation and ecosystem resilience.

## Introduction

Comprising most of South America, Central America, tropical Mexico and the Caribbean Islands, the Neotropics are arguably the most biodiverse region on Earth, being home to at least a third of global biodiversity (1). This region not only includes the largest tropical rainforest, Amazonia, but also eight of the world’s 34 biodiversity hotspots (2). The tropical Andes, in particular, is considered to be the most species-rich region in the world for amphibians, birds, and plants (3), while Mesoamerica and the Caribbean Islands are the richest regions for squamates, and Amazonia has been identified as the primary biogeographic source of Neotropical biodiversity (4). The underlying drivers of the extraordinary biodiversity of the Neotropics are hotly debated in evolutionary ecology, and the mechanisms behind the origin and maintenance of diversity remain elusive (5–9).

Most attempts to explain Neotropical diversity have typically relied on two evolutionary models. In the first, tropical regions are described as the “*cradle of diversity*”, the centre of origin from which species appeared, radiated, and colonized other areas (10– 12). In the other, tropical regions are considered a “*museum of diversity*”, where species suffered relatively fewer environmental disturbances over evolutionary time, allowing ancient lineages to be preserved for millennia (6, 13, 14). Although not mutually exclusive (15), the cradle *vs*. museum hypotheses primarily assume evolutionary scenarios in which diversity expands through time without limits (16). However, expanding diversity models may be limited in their ability to explain the entirety of the diversification phenomenon in the Neotropics; for example, expanding diversity models cannot explain the occurrence of ancient and species-poor lineages in the Neotropics (17–19) or the decline of diversity observed in the Neotropical fossil record (20–22).

A more comprehensive view of Neotropical diversification should consider four alternative evolutionary trajectories of species richness to explain the accumulation of Neotropical diversity (**Fig. 1**):

**Figure 1.**
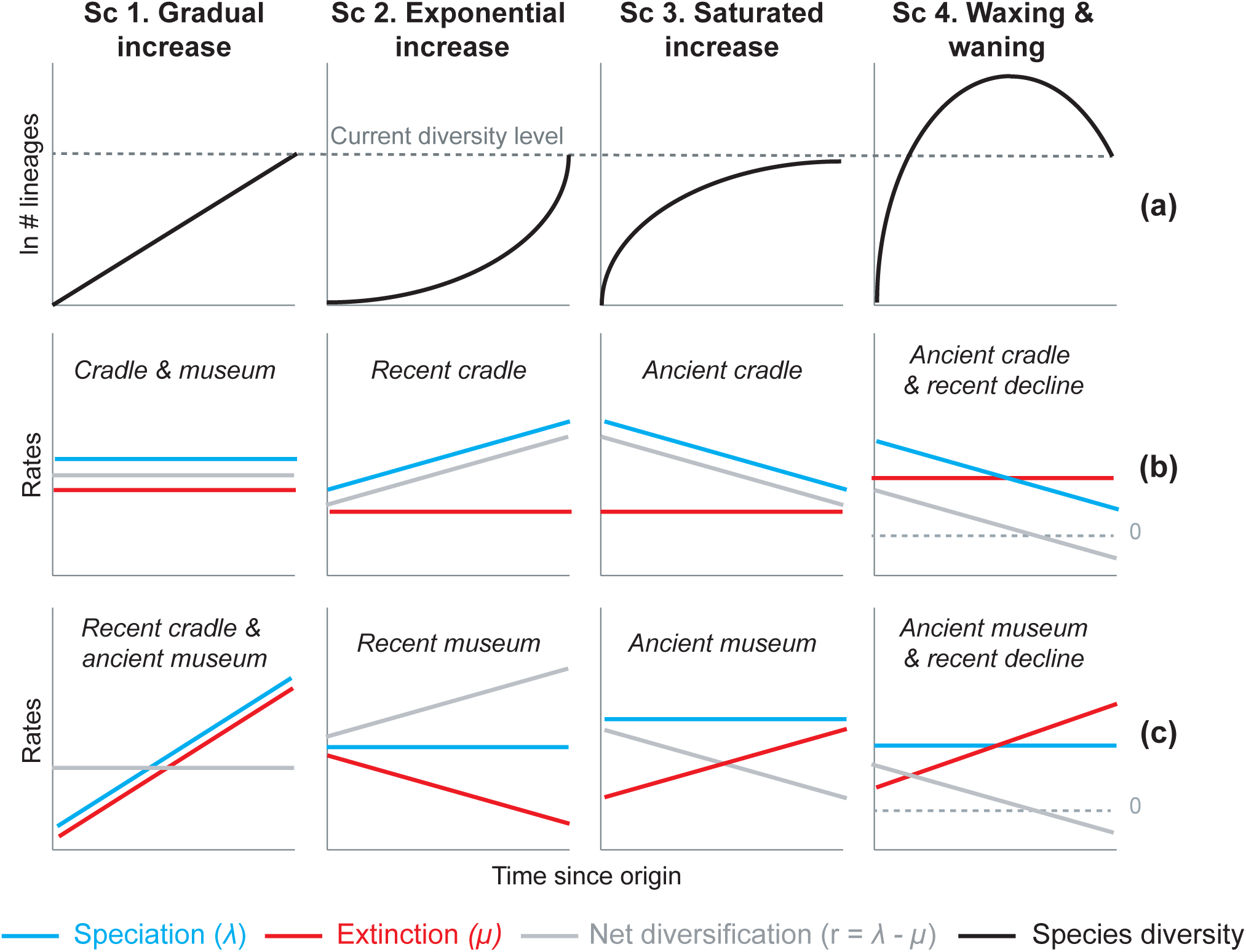
Alternative hypotheses to explain current Neotropical diversity. **(a)** Main species richness dynamics through time, and **(b**,**c)** the alternative evolutionary processes that could generate the corresponding patterns. **(Sc. 1a)** A gradual increase of species richness could result from constant speciation and extinction rates (1b; cradle & museum model), or through a comparable increase in speciation and extinction rates (1c; recent cradle & ancient museum model). **(Sc. 2a)** An exponential increase in species numbers could be attained through constant extinction and increasing speciation (2b; recent cradle model), or constant speciation and decreasing extinction rates (2c; recent museum). **(Sc. 3a)** Saturated increase scenarios, with species accumulation rates slowing down towards the present, could result from constant extinction and decreasing speciation (3b; ancient cradle model), or through constant speciation and increasing extinction rates (3c; ancient museum model). **(Sc. 4a)** Waxing and waning dynamics could result from constant extinction and decreasing speciation (4b; ancient cradle & recent decline model), or constant speciation and increasing extinction (4c; ancient museum & recent decline model). Waxing and waning scenarios differ from saturated increases in that extinction exceeds speciation towards the present, such that diversification goes below 0. Scenarios (b-c) represent the simplest and most general models to explain species richness patterns in (a), but other combinations of speciation and extinction rates could potentially generate these patterns; for example, an exponential increase of species (2a) could also result from increasing speciation and punctual increases in extinction, or through increasing speciation and decreasing extinction.

### Gradual expansions (Scenario 1)

This scenario proposes that species richness accumulated gradually through time in the Neotropics until the present, due, for example, to constant speciation and extinction rates. The gradual increase model received substantial support in the early and recent literature (14, 23–27), and is generally associated with the long-term environmental stability and large extension of the tropical biome across the South American continent (6, 13).

### Exponential expansions (Scenario 2)

An exponential increase in diversity model asserts that species richness accumulated through pulses. Such a pattern can result, for example, from constant extinction and increasing speciation rates. Support for this model generally comes from studies suggesting that geological and climatic perturbations, mostly associated with the elevation of the Andes, promoted pulses of diversification (8, 9, 28). This diversity scenario is probably the most supported across Neotropical studies, although never quantified, with models of increasing speciation (29–38) more often put forward than models of decreasing extinction (18).

### Saturated or asymptotic expansions (Scenario 3)

A saturated diversity model postulates that species richness accumulated more slowly towards the present than in the past, reaching a diversity plateau. This can result from constant extinction and decreasing speciation for example, such that speciation and extinction rates become equal towards the present. Diversification decreases could be due to ecological limits (39), damped increases (40, 41), or abiotic fluctuations (42). Some studies support this model for the Neotropics, and they generally associate it with an early burst of diversification under favourable climatic conditions, followed by decelerations due to global cooling, and dispersal constraints (25, 43–46).

### Declines in diversity (Scenario 4)

Waxing and waning dynamics characterize clades that decline in diversity after periods of expansion. In a declining dynamic, diversification rates also decrease towards the present, but differ from saturated diversity in that extinction exceeds speciation, and diversity is lost. Waxing and waning dynamics may seem unlikely in a tropical context, but evidence for tropical diversity declines has been found at the global scale (47–49) and at the Neotropical scale in the fossil record (20–22, 50–53). Fossil studies additionally suggest a link between decreases in Neotropical diversity and global temperature. For example, plant diversity inferred from fossil morphotypes reached its maximum levels during hyperthermal periods in the Eocene, and decreased sharply with subsequent cooling (20, 21, 54).

Despite an increasing number of evolutionary studies on Neotropical groups, today the prevalence of these alternative modes of species accumulation and diversification (Sc. 1– 4, **Fig. 1**) at a continental scale has been difficult to tease apart empirically (**Question 1**). But such an assessment is required to understand the origin of Neotropical diversity and why the Neotropics are more diverse than other regions in the world. Illuminating the historical causes of Neotropical diversity further requires a closer look at the regional determinants of diversification. Are species richness trends (Sc. 1–4) related to particular environmental drivers (**Question 2**), geographic settings (**Question 3**), or taxonomic groups (**Question 4**) in the Neotropics?

Previous studies indicate that diversification rates might be structured geographically in the Neotropics (55, 56), with geography and climate being strong predictors of evolutionary rate variation (57, 58). For example, speciation may be elevated in regions subjected to environmental perturbations, such as orogenic activity (33, 35, 59–61). However, this view is not without controversy, as some studies found similar diversification patterns among Neotropical regions (27, 62, 63).

Still, little is known about the variation in diversity dynamics (Sc. 1–4, **Fig. 1**) across Neotropical regions at large temporal scales. Most studies investigating spatial patterns of Neotropical diversification focus on long-term diversification dynamics of particular clades, for example, diversification trends of orchids across Neotropical regions (36), or cross-taxonomic patterns in shallow evolutionary time, i.e., present-day speciation rates (27, 58, 63). However, present-day speciation rates might not represent long-term diversification dynamics, specially when rates vary through time. Diversification could be higher in one region than in other without providing information on the underlying diversification trend. Under time-variable rate scenarios, analyzing diversity trends is crucial, but requires changing the focus from species to clades as units of the analyses. Unfortunately, there is still a lack of large-scale comparative data across wide phylogenetic and ecological contexts (59, 64). Given the long history and vast heterogeneity of the Neotropics, general insights can only be provided if long-term patterns and drivers of diversification are shared among Neotropical lineages.

This lack of knowledge may be also due to the challenge of differentiating between evolutionary scenarios based on birth-death models and phylogenies of extant species alone (65, 66). Recent studies have raised concerns on difficulties in identifying parameter values when working with birth-death models under rate variation scenarios (67, 68), showing that speciation (birth, *λ*) and extinction (death, *μ*) rates sometimes cannot be inferred from molecular phylogenies (69). This calls for *(i)* analysing ‘congruent’ models with potentially markedly different diversification dynamics but equal likelihood for any empirical tree (69), or *(ii)* implementing a solid hypothesis-driven approach, in which a small number of alternative hypotheses about the underlying mechanism are compared against data (70).

Based on an unparalleled comparative phylogenetic dataset containing 150 well-sampled species-level molecular phylogenies and 12,512 extant species, we evaluate the prevalence of macroevolutionary scenarios 1–4 (**Fig. 1**) as general explanations for Neotropical diversification at a continental scale (**Q1**), their drivers **(Q2)**, and their variation across biogeographic units **(Q3)** and taxonomies **(Q4)**. To address Q3, we previously identify evolutionary arenas of Neotropical diversification suitable for comparison. Depending on the taxonomic source (1, 71), our dataset represents ∼47–60% of all described Neotropical tetrapods and ∼5–7% of the known Neotropical plant diversity.

## Results

### Neotropical phylogenetic dataset

We constructed a dataset of 150 time-calibrated clades of Neotropical tetrapods and plants derived from densely-sampled molecular phylogenies (**Fig. 2** and *Appendix 1*). The dataset includes a total of 12,512 species, consisting of 6,222 species of plants, including gymnosperms and angiosperms (66 clades, representing 5–7% of the described Neotropical seed plants; Supporting Information Table S1); 922 mammal species (12 clades, 51–77% of the Neotropical mammals; Table S2); 2,216 bird species (32 clades, 47–59% of the Neotropical birds; Table S3); 1,148 squamate species (24 clades, 30–33% of the Neotropical squamates; Table S4); and 2,004 amphibian species (16 clades, 58–69% of the Neotropical amphibian diversity; Table S5). Each clade in our dataset includes 7 to 789 species (mean=83.4), with 53% of the phylogenies including more than 50% of the described taxonomic diversity (sampling fraction mean=57%). Clade ages range from 0.5 to 88.5 Myr (mean=29.9; **Fig. 2**, Fig. S1). Our dataset triples the data presented in previous meta-analyses of the Neotropics in terms of number of species, for example, 214 clades and 4,450 species in (4), and quadruples it in terms of sampling, with 20.8 species per tree in (4).

**Figure 2.**
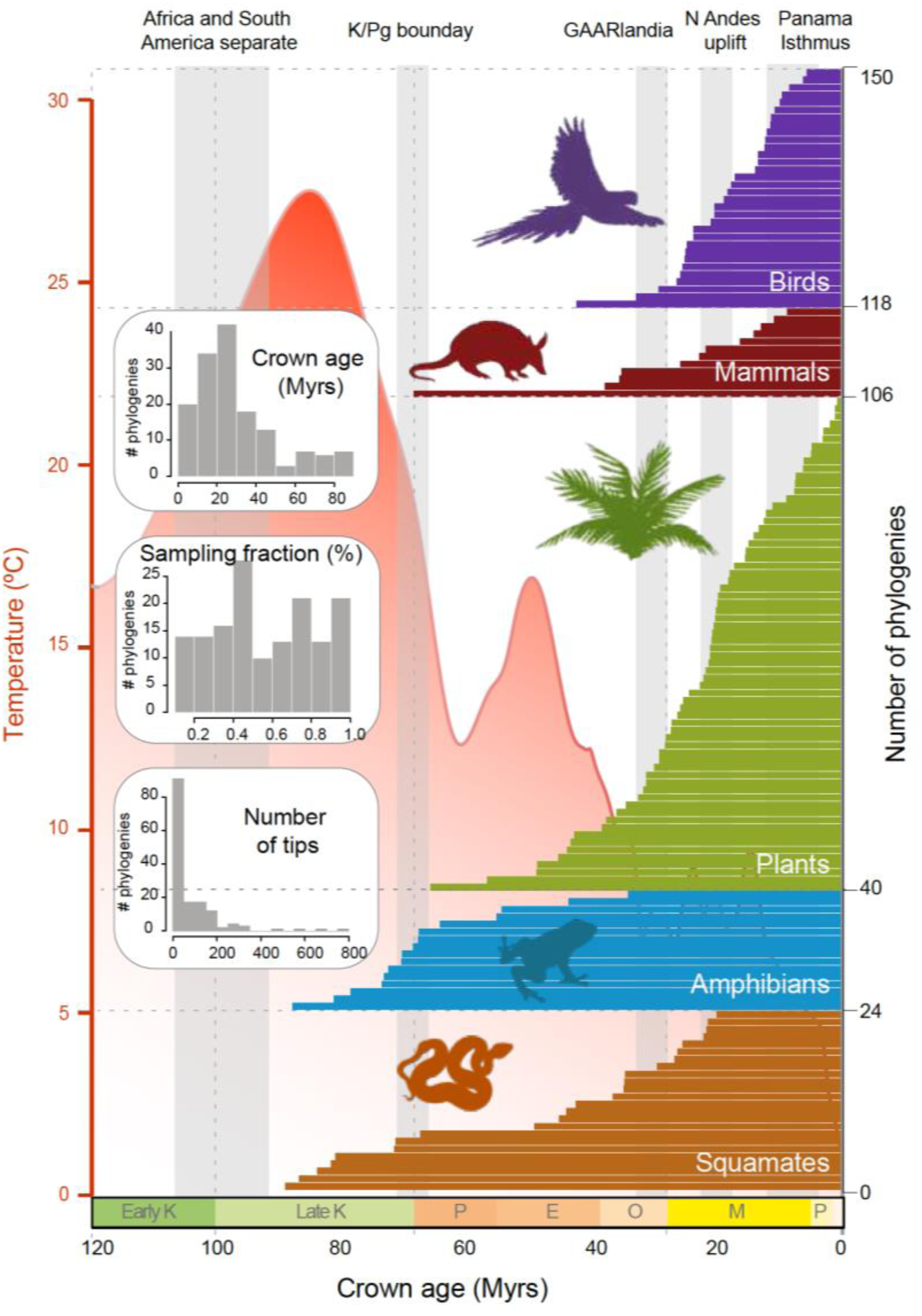
Time of origin for Neotropical tetrapods and plants. Horizontal bars represent crown ages of 150 phylogenies analysed in this study. Shaded boxes represent the approximate duration of some geological events suggested to have fostered dispersal and diversification of Neotropical organisms. Inset histograms represent summary statistics for crown age (mean = 29.9 Myrs), sampling fraction (mean = 57%) and tree size (mean= 83.4 species/tree). Mean global temperature curve from (Zachos et al. 2008). Abbreviations: K, Cretaceous; P, Paleocene; E, Eocene; 0, Oligocene; M, Miocene; P, Pliocene (Pleistocene follows but is not shown); GAARlandia, Greater Antilles and Aves Ridge. Animal and plant silhouettes from PhyloPic (http://-phylopic.org/).

### Estimating the tempo and mode of Neotropical diversification

#### a) Diversification trends based on traditional diversification rates

To understand the tempo (**Q1)** and drivers of Neotropical diversification (**Q2**), we compared the fit of birth-death models applied to 150 phylogenies, including models where diversification rates are constant, vary through time, vary as a function of past global temperatures, or vary according to past Andean elevation (see *Methods*). When only models with constant diversification and time-varying rates were considered, constant models best fit 67% of the phylogenies (101 clades), (Table S6). In the remaining 49 trees, we detected variation in diversification rates. Speciation decreased towards the present in 28 trees (57%), increased in 12, and remained constant (being extinction time-variable) in 9, although the proportions varied between lineages (**Fig. 3**). The proportion of clades that evolved at constant diversification decreased to 50.6% (76 clades) when the comparison included more complex environmental models (**Fig. 4;** *Appendix 2;* Table S7, S8). The proportion of time-variable models also increased to 74 trees.

**Figure 3.**
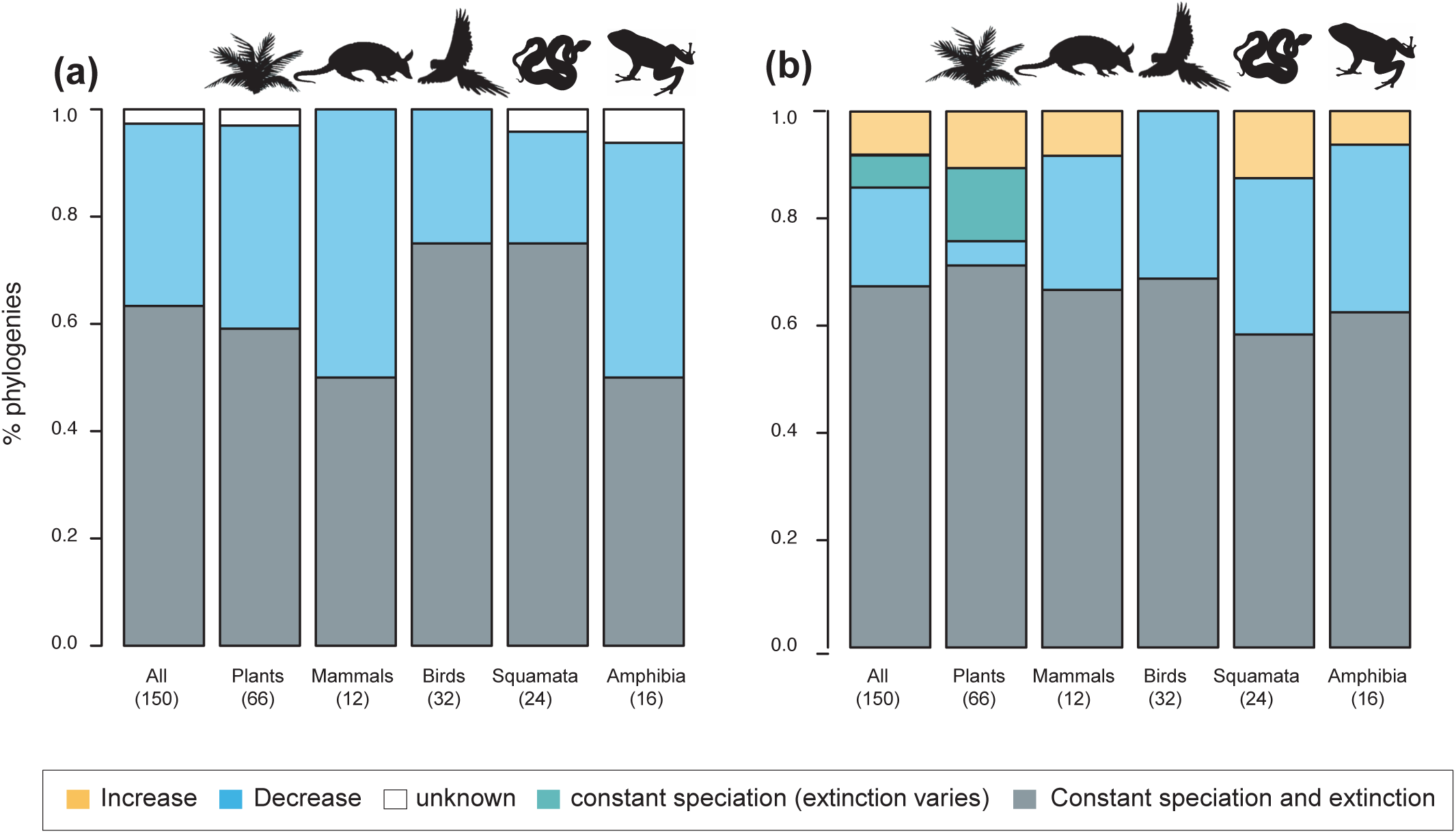
Speciation trends in 150 phylogenies of Neotropical plants and tetrapods. The histograms show the proportion of phylogenies for which constant vs. time-variable diversification models were the best fit, as derived from pulled diversification rates **(a)**, and canonical diversification rates **(b)**, when comparing time-dependent models against constant models. In subfigure b, the proportion of time-variable models is subdivided by the proportion of phylogenies in which speciation rates increase through time, decrease through time, or speciation remains constant (being extinction time-variable). In subfigure a, speciation trends are derived from present-day pulled extinction rates *μp(0):* negative present-day pulled extinction rates values (*μp(0)* < 0) indicate decreasing speciation trends through time (Louca & Pennell, 2020). Positive *μp(0)* > 0 values are possible under both increasing or decreasing speciation rates, in which case speciation trends are designed as “unknown”.

**Figure 4.**
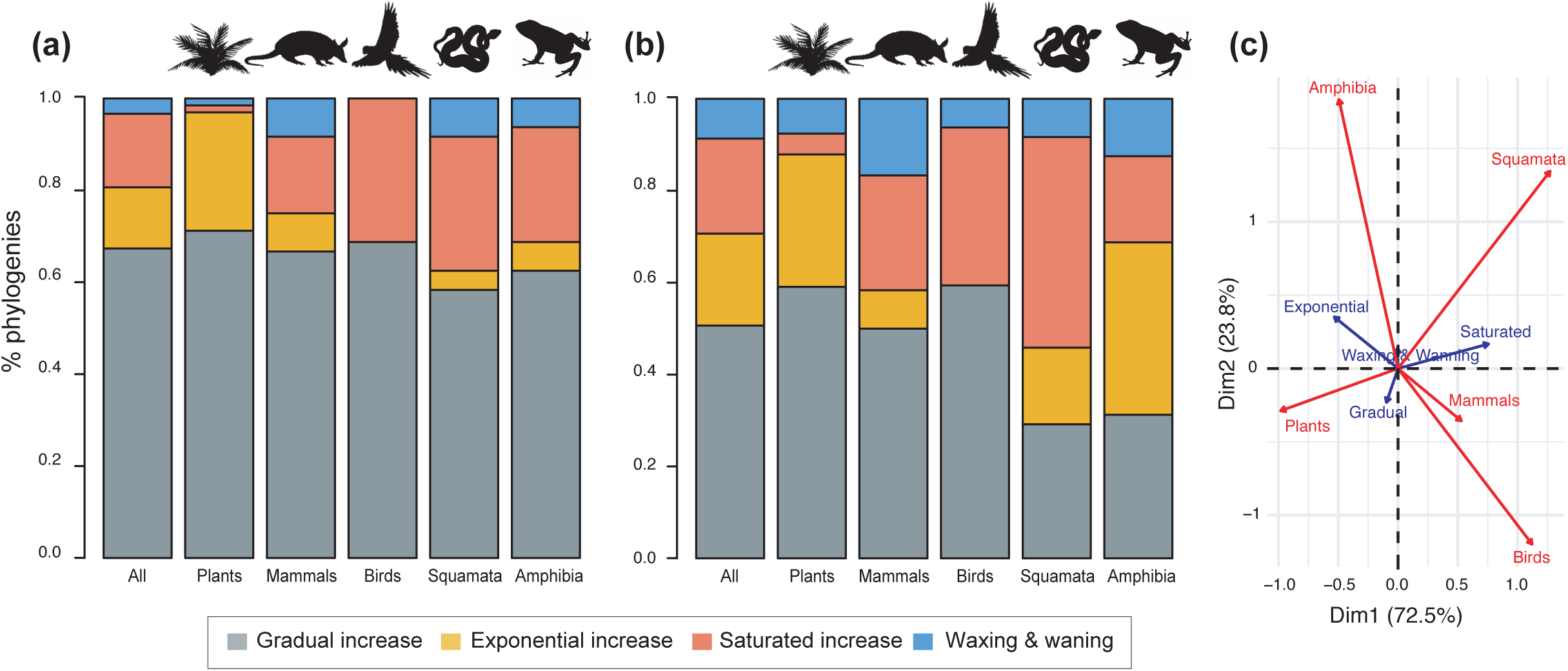
Diversity dynamics in 150 phylogenies of Neotropical plants and tetrapods. The histograms show the proportion of phylogenies for which gradual increase (Sc. 1), exponential increase (Sc. 2), saturated increase (Sc. 3) and waxing & waning (Sc. 4) scenarios were the best fit, as derived from net diversification trends when comparing **(a)** time-dependent models against constant models and **(b)** environmental (temperature- and uplift-dependent models) against time-dependent and constant models. **(c)** Correspondence analysis showing the association between species richness dynamics (represented by blue arrows) and major taxonomic groups (red arrows). If the angle between two arrows is acute, then there is a strong association between the corresponding variables.

The empirical support for the main species richness dynamics from the 150 phylogenies was as follows: gradual expansions (Sc. 1, constant diversification) were detected in 101 to 76 phylogenies if environmental models were considered; exponential expansions (Sc. 2, increases in diversification) were detected in 20–30 clades; and saturated expansions and declining dynamics (Sc. 3 and 4, diversification decreases) were supported in 24–31 and 5–9 clades, respectively (**Table 1** and **Fig. 4**). Diversification trends remained similar when small (<20 species) or poorly sampled (<20% of the species sampled) phylogenies were excluded from the analyses (99 and 137 trees remaining, respectively), although the proportion of constant diversification models decreased in all cases (55–35%; Fig. S2).

**Table 1.**
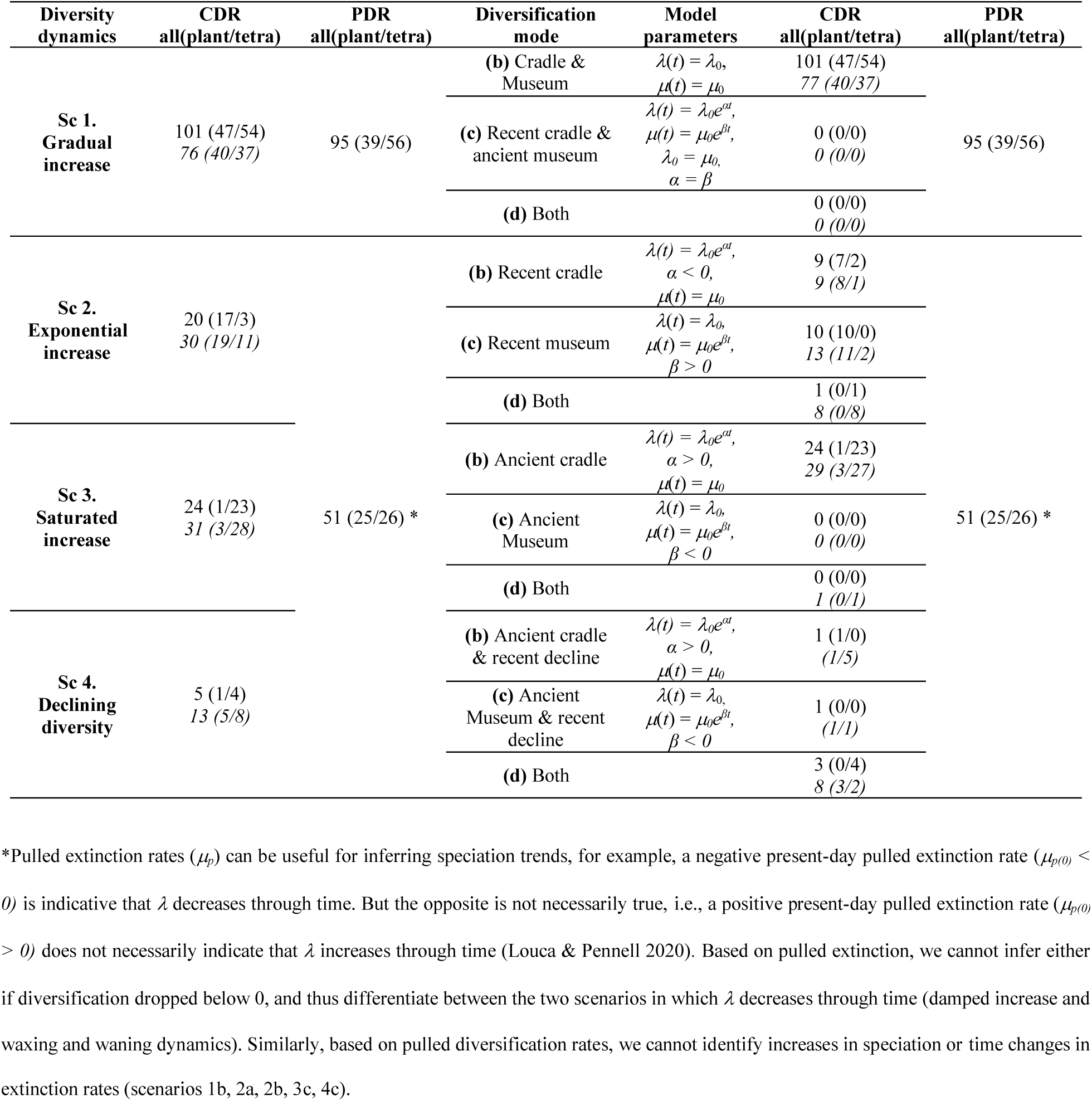
Alternative species richness dynamics (sc. 1–4) and the corresponding diversification models (a–c) able to explain Neotropical diversity. Species richness dynamics represent scenarios of expanding (sc. 1–2), saturating (sc. 3) and contracting (sc. 4) diversity, in which speciation *(λ)* and/or extinction *(μ)* remain constant or vary through time. The number of phylogenies supporting each model is provided for all lineages pooled together, and for plants and tetrapods separately. Empirical support for each evolutionary model is based on canonical diversification rates (CDR), and pulled diversification rates (PDR), by comparing the constant model against different sets of time-variable models. For CDR, we provide as well the results (in italic) based on model comparisons including constant, time-variable, and paleoenvironmental dependent (temperature and uplift) models.

Rate variation was inferred from models that are able to capture the dependency of speciation and/or extinction rates over time (time-dependent models) or over an environmental variable (either temperature- or uplift-dependent models). Among them, temperature-dependent models explained diversification in 40 phylogenies (26.7%). Time-dependent models best fit 17 clades (11%). Uplift-dependent models explained another 11% (**Fig. 5**, Table S7). The relative support for time-, temperature- and uplift-dependent models remained similar regardless of whether we compared the fit of the best or second-best models (defined based on ΔAIC values; Fig. S3), although overall support for constant-rate scenarios decreased in the latter. Results also remained stable regardless of the paleo-temperature curve (72–74) considered for the analyses (see SI; Fig. S4).

**Figure 5.**
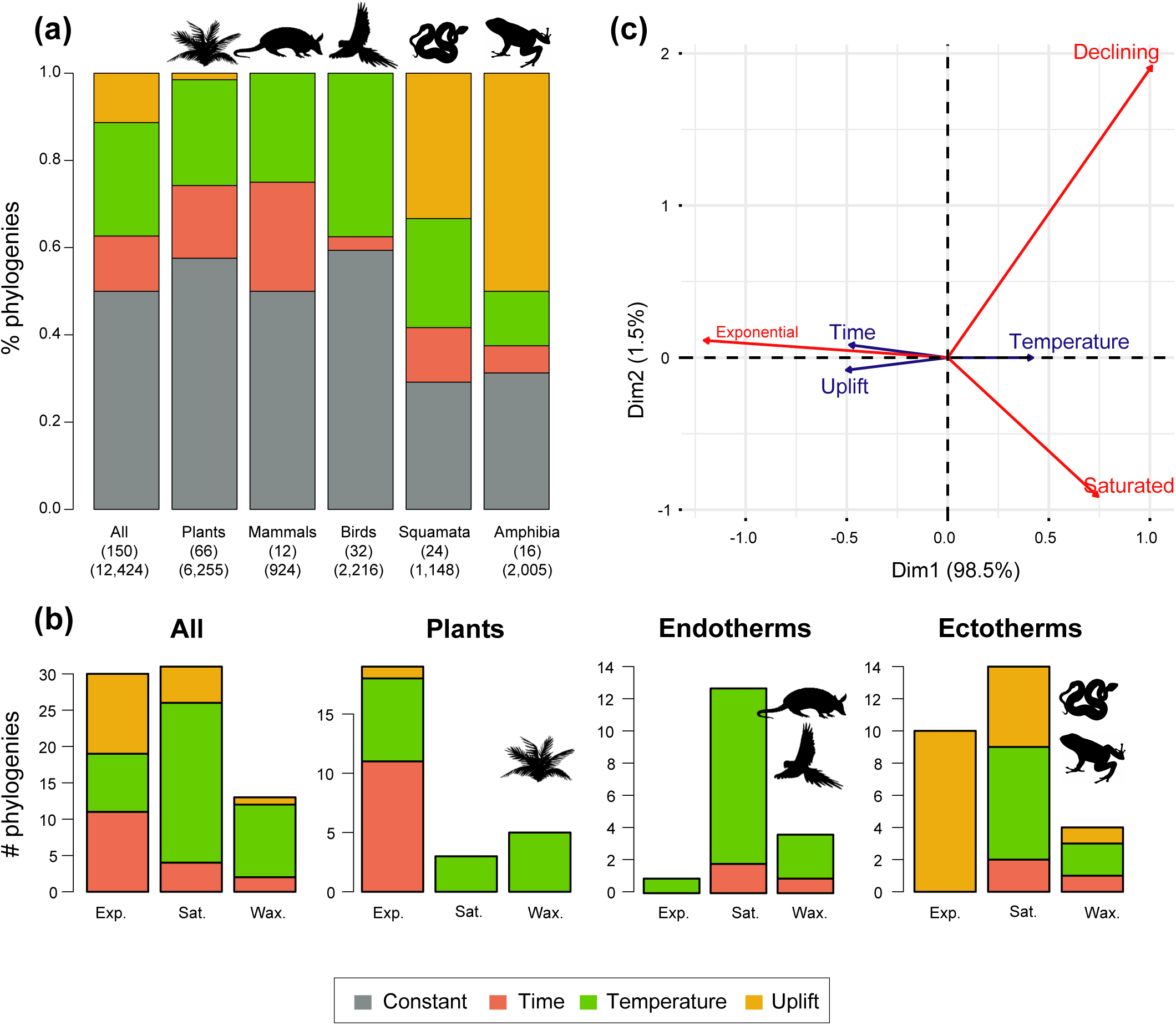
Drivers of Neotropical diversification in 150 phylogenies of Neotropical plants and tetrapods. The histograms report the proportion of ***(a)*** phylogenies whose diversification rates are best explained by a model with constant, time-dependent, temperature-dependent, or uplift-dependent diversification. The number of phylogenies (and species) per group is shown in parentheses. ***(b)*** The histograms report the number of phylogenies whose diversification rates are best explained by a model with constant, time-, temperature-, or uplift-dependent diversification according to different species richness scenarios (Exp = Exponential increase [Sc.2], Sat = Saturated increase [Sc.3] and Dec = Declining diversity [Sc.4]), for plants, endotherm tetrapods, ectotherms and all clades pooled together. ***(c)*** Correspondence analysis for the pooled dataset showing the association between species richness dynamics (represented by red arrows) and the environmental drivers (blue arrows). If the angle between two arrows is acute, then there is a strong association between the corresponding variables.

#### b) Diversification trends based on pulled diversification rates

To gain further insights in Neotropical diversification (**Q1**), we explored congruent diversification models defined in terms of pulled diversification rates (PDR, *r*_*p*_), (69, 75). These analyses recovered consistent diversification trends with those found above: 63% of the phylogenies (95 clades) better fit constant pulled models (**Fig. 3**; Table S9). Meanwhile in 37% of the phylogenies (55 clades) we found variation in PDR through time. Diversification trends remained similar when small (<20 species) or poorly sampled (<20% of the species sampled) phylogenies were excluded from the analyses (Fig. S5). We also detected negative pulled present-day extinction rates *μ*_*p*_*(0)* in most of the phylogenies (51 clades, 92%) in which PDR varied through time, suggesting that speciation was decreasing. Based on PDR, we could only detect constant diversification (Sc. 1) or decreases in speciation, and thus the combined support for Sc. 2, 3 and 4 (**Table 1**).

### Neotropical bioregionalization

To examine the spatial variation of diversification dynamics within the Neotropics (**Q3**), we first had to identify geographic units of long-term Neotropical evolution suitable for comparison. We found that most clades in our study were distributed in most Neotropical WWF ecoregions (Table S10), suggesting that species presence-absence data might be of limited use for delimiting geographic units at the macroevolutionary scale of this study. In contrast, based on clades’ abundance patterns, we identified 5 clusters of regional assemblages that represent long-term clade endemism (**Fig. 6a, b;** Fig. S6; Table S11): cluster 1 (including the Amazonia, Central Andes, Chocó, Guiana Shield, Mesoamerica, and Northern Andes), cluster 2 (Atlantic Forest, Caatinga, Cerrado, Chaco, and temperate South America), cluster 3 (Caribbean), cluster 4 (“elsewhere” region), and cluster 5 (Galapagos). An alternative clustering (Fig. S7) separating Mesoamerica from cluster 1, and the Chaco and temperate South America from cluster 2, received lower support (Fig. S6).

**Figure 6.**
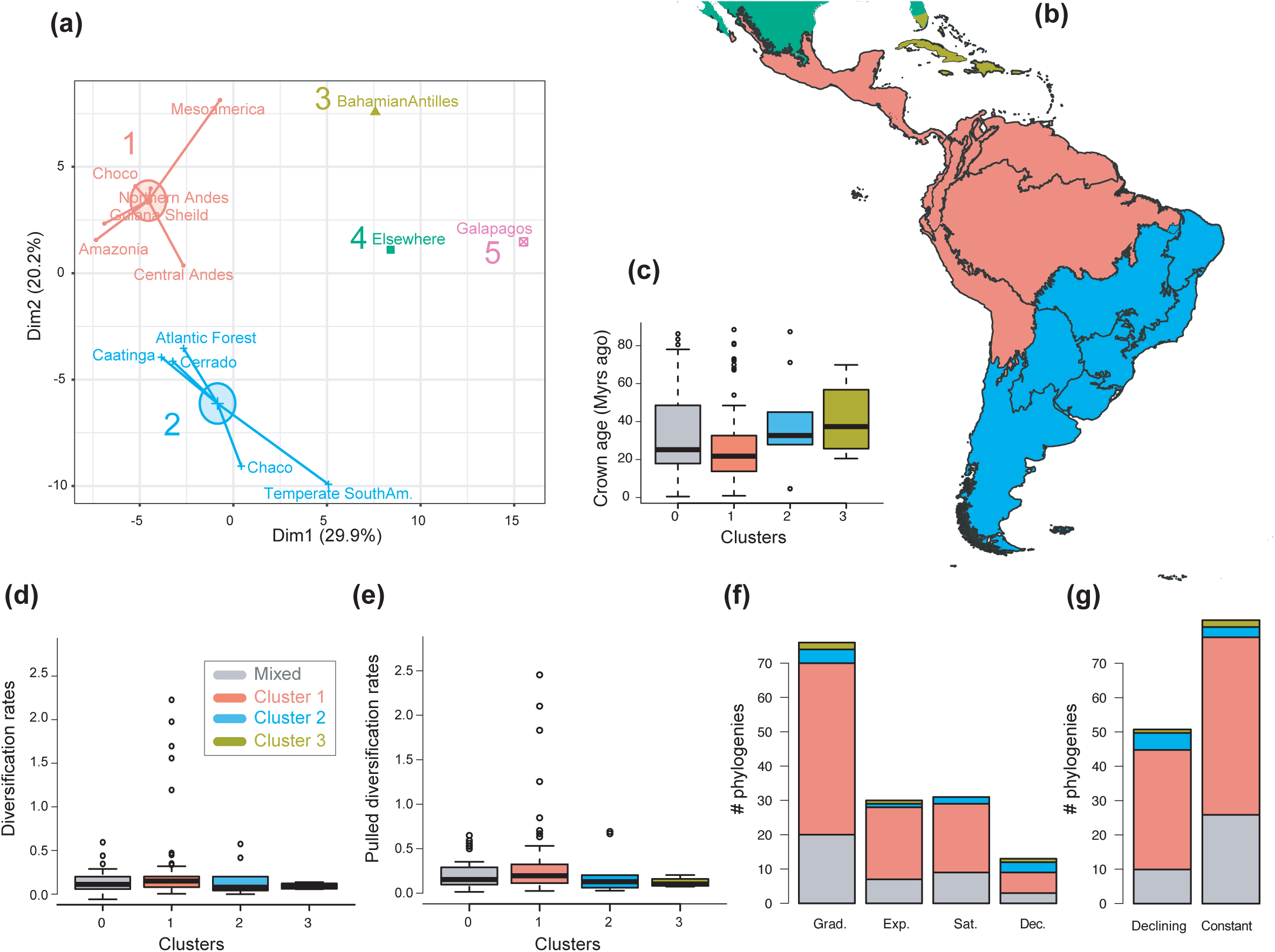
The geographical structure of Neotropical diversification. **(a)** PCA representation of the 5 biogeographic clusters identified based on K-means clustering of 13 areas (WWF ecoregions) and 150 clades. **(b)** Resulting clusters (1–5) in the geographic space. Colors correspond with the biogeographic clusters in (a). Thick lines delineate the original 13 ecoregions used in the analyses. **(c)** Box plot showing differences in crown age of the phylogenies distributed in each of the biogeographic clusters. **(d)** Variation in diversification and **(e)** pulled diversification rates (derived from the constant-rate model) across geographic clusters. **(f)** Number of phylogenies for which species richness scenarios Sc. 1–4 (Grad = Gradual increase [Sc.1], Exp = Exponential increase [Sc.2], Sat = Saturated increase [Sc.3] and Dec = Declining diversity [Sc.4]) were the best fit, across geographic clusters as derived from cannonical diversification rates. **(g)** Number of phylogenies for which constant *vs*. declining speciation rates were the best fit, across geographic clusters as derived from pulled diversification rates.

### Variation of diversification dynamics across taxa, environmental drivers, and biogeographic units

We evaluated the prevalence of macroevolutionary scenarios 1–4 (**Fig. 1**) across environmental drivers **(Q2)**, biogeographic units **(Q3)** and taxonomies **(Q4)** (see *Methods*). **Table 2** summarize all the results. We found that species richness dynamics were related to particular environmental drivers (*p*=0.003; **Q2**). Pairwise comparisons indicated that temperature-dependent models tended to best fit clades experiencing saturating (*p*=0.049) and declining (*p*=0.05) diversity dynamics. Meanwhile, uplift- and time-dependent models tended to best fit clades with exponentially increasing diversity (*p*=0.03) (**Fig. 5c**).

**Table 2.** Summary *p* value results derived from the analysis of canonical diversification (*r*) and pulled diversification (*r*_*p*_) rates. Significant differences in the proportion of clades experiencing different diversity trajectories (based on canonical diversification rates: gradual expansions, exponential expansions, saturation or declining diversity; based on pulled diversification rates: expanding vs. declining speciation) across biogeographic units, elevations, taxonomic groups, and environmental drivers as derived from Fisher’s exact tests. Significant differences in net diversification, pulled diversification, and speciation rates across biogeographic units, elevations and taxonomic groups derive from Kruskal-Wallis chisquared analyses. Significant results are highlighted in bold.

|  | Diversity trajectories |  | Diversification rates |  | Speciation rates |
| --- | --- | --- | --- | --- | --- |
| | $r$ | $r_p$ | $r$ | $r_p$ | $r$ |
| **Biogeographic units (5 clusters)** | 0.459 | 0.252 | 0.168 | 0.083 | 0.248 |
| **Biogeographic units (7 clusters)** | 0.503 | 0.947 | 0.198 | 0.424 | 0.277 |
| **Elevation** | 0.504 | 0.839 | 0.672 | 0.277 | **0.034** |
| **Elevation (lowland-montane combined)** | 0.062 | 0.062 | 0.332 | 0.869 | **0.031** |
| **Taxonomic groups** | **0.000** | 0.126 | **0.000** | **0.000** | **0.000** |
| **Environmental drivers** | **0.003** | – | – | – | – |

In contrast, there is no evidence to suggest that species richness dynamics are related to particular geographic conditions when considering the whole dataset (**Fig. 6c–f; Q3**). Results of the Fisher’s exact test show no significant differences in the proportion of clades experiencing gradual expansions, exponential expansions, saturation or declining diversity across biogeographic units (*p*=0.45) or elevation ranges (*p*=0.062). We obtained similar results when the montane category was analyzed separately (*p*=0.5, Fig. S8). Analyses based on PDR produce the same results, with no differences in the proportion of clades experiencing constant (i.e., expanding diversity dynamics) or declining speciation trends across biogeographic units (*p*=0.25), or elevation ranges (*p*=0.062), even when the montane category was analyzed separately (*p*=0.839). Estimates of net diversification rates (rather than diversity trajectories) derived from the constant diversification model did not differ across biogeographic units (χ^2^=5.05, *p*=0.17) or altitudinal ranges (χ^2^=2.20; *p*=0.332) either. Speciation rates did not differ between biogeographic units (χ^2^=4.1, *p*=0.25), but did vary across altitudinal ranges (χ^2^=6.9, *p*=0.03). Speciation rates were significantly higher across highland taxa (**Fig. 6**). In addition, PDR did not differ across biogeographic units (χ^2^=6.7; *p*=0.083) or elevations (χ^2^=0.28; *p*=0.87).

Finally, diversity trajectories (Sc. 1–4) differed across taxonomic groups (*p*<0.0001, Fisher’s exact test; **Q4**). Pairwise comparisons indicated that plants differed significantly from birds in the proportion of gradual (*p*<0.02), exponential (*p*<0.02) and saturated (*p*<0.0001) increase models after correcting for multiple comparisons. Birds also differed from amphibians in the proportion of saturated and exponential increases (*p*<0.02). Plants differed from squamates in the proportion of exponential (*p*<0.0006) and saturated (*p*<0.008) increases (**Fig. 4c**). Net diversification rates were also significantly lower for Neotropical ectotherm tetrapods than for endotherms and plants (Kruskal-Wallis chi-squared: χ^2^=36.7, *p*<0.0001) (**Fig. 7**). We also found statistically significant differences in speciation rates across groups (χ^2^=60.8, *p*<0.0001): plants showed higher speciation rates than endotherms, the latter, in turn, with higher speciation rates than ectotherms.

**Figure 7.**
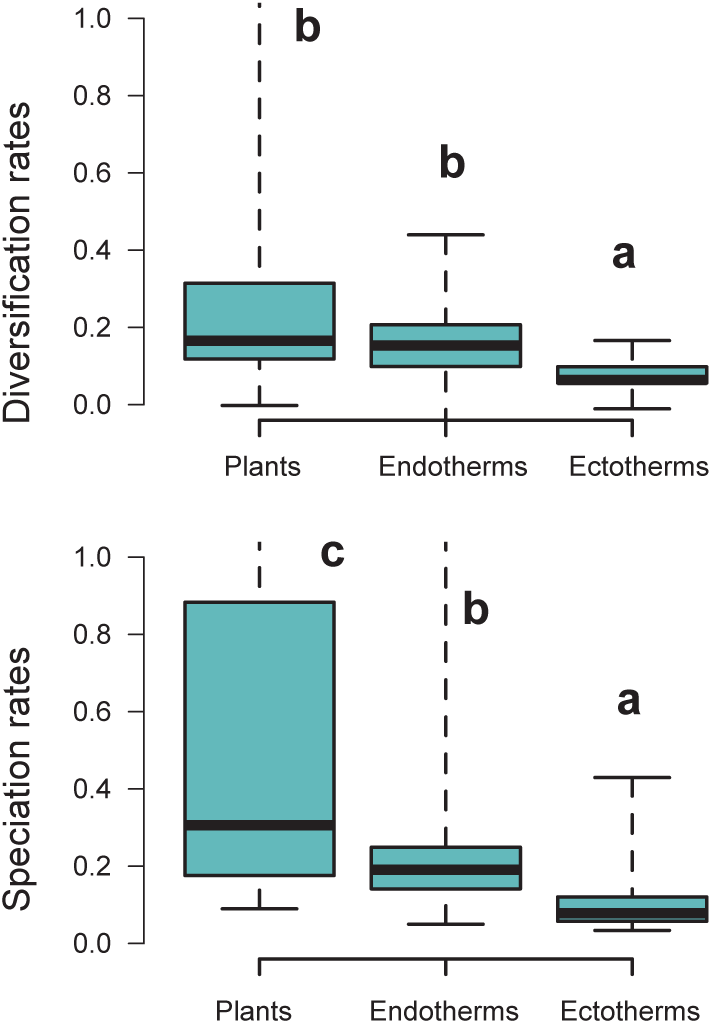
Diversification rates compared across plants and tetrapods (endotherms and ectotherms). Diversification and speciation rates are derived from the constant-rate model (Table S6). Letters are used to denote statistically differences between groups, with groups showing significant differences in mean values denoted with different letters. The y-axis was cut off at 1.0 to increase the visibility of the differences between groups. Upper values for plants are therefore not shown, but the quartiles and median are not affected. Units are in events per million years.

The number of species per phylogeny differed between model categories (phylogenetic ANOVA: *F*=10.9, *p=*0.002). Clades fitting gradual expansion models tended to have less species than clades fitting exponential (*p*=0.006) and declining (*p*=0.03) dynamics (Fig. S9). Taxon sampling, however, did not differ significantly (*F*=4.5, *p=*0.53). Crown age differed between model categories, being on average younger for gradual scenarios than for exponential (*p*=0.03) and declining (*p*=0.03) dynamics.

## Discussion

### Diversification dynamics

Neotropical biodiversity has long been considered as being in expansion through time due to high rates of speciation and/or low rates of extinction, either through a long-term (13, 27) or a recent (71) cradle and museum of diversity. Yet, the prevalence of the expanding dynamic was never quantified. The higher support for the expanding diversity trend found here align with these ideas. The majority of Neotropical clades (between 80% to 70%, if environmental models are considered) displayed expanding diversity dynamics through time (**Fig. 4**; **Table 1**). Most of these clades experienced a gradual accumulation of lineages (Sc. 1; between 67– 50%), and a lower proportion (14–16%) expanded exponentially (Sc. 2), thus diversity accumulation accelerated recently. Results based on PDR support these conclusions, with the largest proportion of clades expanding diversity (63%) due to gradual increases (Sc. 1; **Fig. 3**).

Our results also provide evidence that cradle/museum models are not sufficient to explain Neotropical diversity. Based on traditional diversification rates, 16–21% of the Neotropical clades, mostly tetrapod clades, underwent a decay in diversification, hence a slower accumulation of diversity toward the present (Sc. 3). While a pervasive pattern of slowdowns in speciation has been described at various geographic and taxonomic scales, e.g., (41, 43, 76, 77), Neotropical tetrapod diversity levels have only rarely been perceived as saturated (25, 27, 43, 46). Furthermore, waxing-and-waning dynamics (Sc. 4) also characterize the evolution of 3–9% of the Neotropical diversity, consistent with paleontological studies (20–22). We found that the species richness of five plant and eight tetrapod clades decined toward the present (e.g., *Sideroxylon* [Sapotaceae], *Guatteria* [Annonaceae], caviomorph rodents, Thraupidae birds, or Lophyohylinae [Hylidae] frogs). This proportion might seem minor but is noteworthy when compared with the low support for this model found in the Neotropical literature, which could be explained by the difficulties in inferring negative diversification rates based on molecular phylogenies (66). Inferring diversity declines is challenging, and often requires accounting for among-clade rate heterogeneity (78). As shown here, incorporating environmental evidence could also help identifying this pattern, increasing support for this scenario relative to the comparisons without these models (**Fig. 4**).

Support for decreasing diversification through time was larger when PDR were considered; 34% of the clades showed slowdowns in speciation (**Fig. 3**). Based on PDR, however, we cannot infer if declines of speciation were accompanied by constant, declining or increasing extinction (69), and thus determine the relative support for Sc. 2–4. If declines in speciation were accompanied by larger extinction declines, it would be still possible to recover expanding dynamics (Sc. 1, 2), but in most other cases, they would lead to declines in diversification (Sc. 3, 4). The limited interpretability of PDR prevents the extraction of further conclusions based on these rates (70).

Still, our study illustrates the robustness of the diversification trend in the Neotropics to different modelling approaches. Despite parameter values varying substantially for some trees between the traditional and PDR methods (Table S7, S9, **Fig. 6 d, e**), a pattern also described in recent studies (70), our analyses support the generality of the expanding diversity trend for most clades in the Neotropics (**Fig. 4**).

#### Sampling artefacts and reliability of these results

The pattern found in this study appears to be robust to sampling artefacts. Clade age and tree size can partially explain the better fit of the phylogenies to the constant diversification model, thus the gradually-expanding trend (Sc. 1), although these factors do not explain the relative support between time-varying increasing (Sc. 2) versus decreasing (Sc. 3, 4) scenarios (Fig. S9). Constant diversification prevails among recently-originated and species-poor clades, which may reflect that these clades had less time to experience changes in diversification. Alternatively, it has been suggested that the power of birth–death models to detect rate variation can decrease with the number of species in a phylogeny (79), suggesting that tree size could hinder the finding of rate-variable patterns. Still, the largest support for the expanding diversity trend persisted (72–60% of clades) after excluding small trees from the analyses (<20 species; Fig. S2). Then, the relative support for the exponentially expanding scenario (Sc. 2) increased at the expense of the gradually expanding scenario (Sc. 1), strengthening the conclusion of the generality of the expanding trend in the Neotropics.

It is also known that incomplete taxon sampling has the effect of flattening out lineages-through-time plots towards the present and artificially increasing the detection of diversification slowdowns (80). If this artefact affected our results, we would expect to see that weakly-sampled phylogenies tend to fit saturated diversity models (Sc. 3). Instead, we found that sampling fraction did not differ between lineages fitting saturated and expanding diversity models (Fig. S9), and the proportion of clades fitting saturated models increased (17–22%) after excluding poorly-sampled phylogenies (<20% of the species sampled; Fig. S2).

### Taxon-specific patterns and drivers of diversification

Our study revealed contrasting evolutionary patterns for plants and tetrapods in the Neotropics (**Fig. 4**). Diversity expansions (Sc. 1, 2) were more frequent in plants (∼88%, 59 clades) than in tetrapods (∼57%, 48 clades). In contrast, asymptotic increases (Sc. 3) were more frequently detected in tetrapods (33%, 28 clades) than in plants (4.5%, 3 clades; *Tynanthus* [Bignoniaceae], Chamaedoreae [Arecaceae], and Protieae [Burseraceae]). The study of PDR did not help to confirm/reject these conclusions. This contrasting evolutionary pattern may result from differential responses of plants and tetrapods to environmental changes (**Fig. 5**).

Global temperature change during the Cenozoic is found to be the main driver behind diversification slowdowns (Sc. 3) and declines (Sc. 4), especially for endotherms (**Fig. 5**). The positive correlation between diversification and past temperature in our temperature-dependent models indicates these groups diversified more during warm periods, such as the Eocene or the middle Miocene, and diversification decreased during cool periods. This result is in agreement with a recent study showing a negative effect of climate cooling (and a positive effect of Andean orogeny; see below) on Neotropical tetrapod diversification based on spatially explicit simulation models of diversification (81). Only the New World monkeys (Platyrrhini), diversified more as temperature dropped. This could reflect the role of Quaternary events on primate speciation (9), and/or be an artefact of taxonomic over-splitting in this clade (82). In contrast, for the few plant clades influenced by temperature changes, diversification increased during Neogene cooling (i.e., a negative correlation between diversification and temperature).

Diversification slowdowns and saturating dynamics have often been interpreted as the signal of ecological limits on the number of species within a clade, implying that diversity is bounded (39, 83). Among the tetrapod phylogenies supporting diversification slowdowns, time-dependent models explain only 3% of them (4 phylogenies; **Fig. 5**, Table S7), suggesting that ecological limits do not play an important role. Time-dependent models with decreasing speciation have been suggested to be a good approximation of diversity-dependent diversification, whereby speciation rates decline as species accumulate (84, 85). In fact, recent studies show time- and diversity-dependent models are not distinguishable based on extant phylogenies (86). Our results lend support to an alternative explanation for diversification slowdowns: the idea that tetrapods fail to keep pace with a changing environment (42, 87). According to the Metabolic Theory of Biodiversity, high temperatures can increase enzymatic activity, generation times and mutation rates (88), which may in turn affect diversification (89). Climate cooling could also decrease global productivity, resource availability, population sizes (90) or even species interactions (91).

Diversification increases are attributed to different factors in our study, including the uplift of the Andes and temperature cooling (**Fig. 5)**. The Andean orogeny mostly impacted tetrapod diversification, especially ectotherms. Diversification of some lineages increased as the Andes rose, including Andean-centred lineages such as Liolaemidae lizards, but also others predominantly distributed outside the Andes, such as Leptodactylidae frogs. Sustained diversification in the context of Andean orogeny, both into and out of the Andean region, could be explained by increasing thermal and environmental gradients, from the equatorial areas to Patagonia or from west-east (92, 93). Other possible correlates include changes in elevational distributions of lineages (94, 95), or recurrent migrations (25, 33).

In contrast, plant diversity expansions were primarily associated with temperature cooling and with time, where the latter represents a null hypothesis; the better fit of a time-dependent model, in comparison to environmental models, is generally indicative of factors not being investigated here (85). Many of the plant lineages fitting time-dependent models represent textbook examples of ongoing radiations; e.g., centropogonids (35), *Lupinus* (34), or *Inga* (96) whose diversification has been associated with biotic drivers, such as the evolution of key adaptations or pollination syndromes. These factors are taxon-specific and were not evaluated in this study, where we focused on global phenomena. Our results therefore add support to the role of environmental and biotic factors as non-mutually exclusive drivers of macroevolutionary changes on Neotropical plants.

Our plant dataset represents just a fraction (> 6,000 species, 66 clades; ∼7% of the total species) of the vast diversity described in the region. As such, future investigations would be necessary to confirm the generality of the expanding trend for plants. Still, our results suggest that Cenozoic environmental changes stimulated plant diversification, while they drove diversification slowdowns for some tetrapods, with plants adapting to deteriorating climatic conditions. Higher (mean) speciation rates in plants than in tetrapods (**Fig. 7**) could have provided plant lineages more opportunities for adaptation to new environments. Greater dispersal abilities in plants (4, 97) may also explain this pattern.

In our study, the relative support for time-, temperature- and uplift-dependent models remained stable to AIC variations (Fig. S3). Model support also remained stable regardless of the paleo-temperature curve considered for the analyses (Fig. S4). Still, the power of AIC values to differentiate model support has been recently questioned (42, 69). As such, these results should be interpreted with caution. The use of an hypothesis-driven framework has been suggested as a potential solution to alleviate the problem of non-identifiability of diversification parameters, by setting up explicit prior assumptions and delimiting the potential parameter space (69, 70, 98). Here, we do not evaluate every possible factor that could potentially explain Neotropical biodiversity, but only confront scenarios capturing well-established hypotheses on Neotropical diversification. We focus on the role of the Andes and climatic oscillations because they have previously been pinpointed as essential for explaining Neotropical biodiversity (8, 57, 81). Thus, our interest is to explore which of these factors best explains the data compiled, although other factors could have played a role.

### The geographical structure of Neotropical diversification

Understanding the spatial variation of biodiversity dynamics is key to understanding the determinants of the exceptional diversity of the Neotropics. The first step towards this is the identification of evolutionary arenas of Neotropical diversification.

Conventional bioregionalizations schemes, such as biomes (99), ecoregions (100) or other pre-defined biogeographic units (4, 101, 102), could represent evolutionary arenas of diversification suitable for comparison. Often, these schemes have been shown to be useful for categorizing actual species ranges, but our study shows they are less appropriate for examining lineage endemism at the macroevolutionary scale (i.e., above the species level); most clades in our study are distributed in most Neotropical ecoregions (Table S10). The lack of a clear geographical structure for taxa of higher rank could be explained by the fact that conventional bioregionalizations generally represent categorizations based on data on the contemporary distribution of species without explicitly considering ancestral distributions or the relationships among species (103, 104).

We propose here an alternative bioregionalization scheme of the Neotropical region that accounts for long-term regional assemblages at macroevolutionary scales (**Fig. 6**). We identify five biogeographic units that represent macroregions of shared evolutionary history for Neotropical clades – macroregions where independent Neotropical radiations occurred over millions of years of biotic evolution. These regions are defined in terms of species-richness patterns within clades (Table S10), showing that species-rich clades in the Amazonia, also tend to be species rich in the Andes, Chocó, Guiana Shield, and Mesoamerica (biogeographic cluster 1), without excluding that some species within these clades occur in other regions. Meanwhile, clades that are species-rich in the Atlantic Forest tend to be rich in the Caatinga, Cerrado, Chaco, and temperate South America (cluster 2). As such, these clusters form distinctive units of Neotropical evolution and represent long-term clade endemism.

The biogeographic cluster 1 roughly corresponds with a broad ‘pan-Amazonian’ region that relied on the ancient Amazon Craton (8). Cluster 2 broadly groups different formations of the area known as the ‘Dry Diagonal’ (105, 106), which are geologically younger, dating from the Miocene (107–109), although lineage crown ages do not differ between these regions (**Fig. 6**). Clusters 1 and 2 include regions identified as transition zones in previous studies – Mesoamerica and temperate South America, respectively (110). Our analyses merged these regions with the core area with which it showed the greatest affinity, although other less supported classification schemes separate transition regions into individual clusters (Fig. S6, S7).

When examining the connection between geography and long-term diversity dynamics, we did not find evidence to reject the null hypothesis of equal diversification, with similar diversity dynamics (Sc. 1–4) found across the biogeographic units of Neotropical evolution identified here (**Fig. 6, Table 2**). We obtained the same result when Mesoamerica and temperate South America transition zones were analysed separately (Fig. S7). In addition, we did not find differences in diversification dynamics between elevational ranges. These results were consistent whether we analysed net diversification rates or their derived diversity trends (Sc. 1–4). In the former, Neotropical lineages distributed in different elevations did differ in their speciation rates, as found in previous studies: speciation increased with altitude (34, 46, 58, 59, 111). Elevated speciation rates might result from ecological opportunities on newly formed high-altitude environments, or those newly exposed after periods of cooling (112–114). However, elevated speciation rates were also accompanied by elevated extinction in these habitats, hence net diversification remains comparable. The hypothesis of comparable diversification was also supported when comparing PDR (**Fig. 6**).

The lack of any clear geographic structure of diversification reveals that the evolutionary forces driving diversity in the Neotropics acted at a continental scale when evaluated over tens of millions of years. Despite the suggestion that present-day diversification rates are structured geographically in the Neotropics (55, 57), this variation might not represent long-term evolutionary dynamics. Evolutionary time and extinction could have eventually acted as levelling agents of diversification across the Neotropics over time. Geographic diversification may vary within taxonomic groups, though small sample sizes prevent us from drawing any firm conclusions on this. Future studies evaluating this question on a greater sample of Neotropical clades could help to assess the generality of this pattern.

### Conclusion

The study of macroevolution using phylogenies has never been more exciting and promising than today (85, 115). The successful development of powerful analytical tools, in conjunction with the rapid increase in the availability of biological data, will allow us to disentangle complex diversification histories. We still face important limitations in data availability and methodological shortcomings, e.g., (68, 69, 86), but by acknowledging them we can better target our joint efforts as a scientific community (116). We hope that our approach will encourage movement in that direction, and that it will provide interesting perspectives for future investigations in the Neotropics and other regions.

This study represents a quantitative assessment of the prevailing macroevolutionary dynamics in the Neotropics, and their spatial variation, at a continental scale. Whether the pattern found here, i.e., a predominant expansion of diversity over time, can contribute to explain why the Neotropics have more species than other regions in the world remains to be evaluated based on comparative data from other regions (117). Such a comparison could reveal contrasted diversity trajectories in different continents, and help to elucidate the association between current diversity levels and long-term diversity dynamics. By identifying distinct evolutionary arenas of long-term Neotropical diversification, we hope our study will also stimulate research to examine hitherto investigated macroecological questions in shallow evolutionary times, from a deep-time and clade-level macroevolutionary perspective.

Our results also have implications for discussing the future of biodiversity in the context of current environmental changes and human-induced extinction (118). If half of the lineages in our study expanded diversity through time following a constant diversification mode, it may take tens of millions of years for biodiversity to reach its pre-extinction level.

## Methods

### Data compilation

Neotropical clades, representing independent radiations in the Neotropics, were pulled from large-scale time-calibrated phylogenies of frogs and toads (95), salamanders (119, 120), lizards and snakes (121), birds (56) (including only species for which genetic data was available), mammals (122, 123), and plants (124). To identify independent Neotropical radiations, species in these large-scale phylogenies were coded as distributed in the Neotropics – delimited by the World Wide Fund for Nature WWF (100) – or elsewhere using the R package *speciesgeocodeR* 1.0-4 (125), and their geographical ranges extracted from the Global Biodiversity Information Facility “GBIF” (https://www.gbif.org/), the PanTHERIA database (https://omictools.com/pantheria-tool), BirdLife (http://www.birdlife.org) and eBird (http://ebird.org/content/ebird), all accessed in 2018, in a procedure similar to (71). Next, we pruned the trees to extract the most inclusive clades that contained at least 80% Neotropical species, as previously defined. This procedure ensures that the diversification signal pertains to the Neotropics. In addition, phylogenies of particular lineages not represented in the global trees (or with improved taxon sampling) were obtained from published studies or reconstructed *de novo* in this study (for caviomorph rodents, including 199 species; see SI). In the case of plants and mammals, most phylogenies were obtained from individual studies, given the low taxon sampling of the plant and mammal large-scale trees. However, whenever possible, we extracted phylogenies from a single dated tree rather than performing a meta-analysis of individual trees from different sources (8, 126), such that divergence times would be comparable. The resulting independent Neotropical radiations could represent clades of different taxonomic ranks. We did not perform any specific selection on tree size, crown age, or sampling fraction, but tested the effect of these factors on our results.

### Estimating the tempo and mode of Neotropical diversification

#### a) Diversification trends based on traditional diversification rates

We compared a series of birth-death diversification models estimating speciation *(λ)* and extinction *(μ)* rates for each of the 150 phylogenies with the R package *RPANDA* 1.9 (127). To make these results comparable with those derived from pulled diversification rates below, we followed a sequential approach by including models of increasing complexity. We first fitted a constant-rate birth–death model and compared it with a set of three models in which speciation and/or extinction vary according to time (78): *λ(t)* and *μ(t)*. For time-dependent models, we measured rate variation for speciation and extinction rates with the parameters *α* and *β*, respectively: *α* and *β* > 0 reflect decreasing speciation and extinction toward the present, respectively, while *α* and *β* < 0 indicate the opposite, increasing speciation and extinction toward the present.

We further compared constant and time-dependent models, described above, with a set of environment-dependent diversification models that quantify the effect of environmental variables on diversification (Q2) (128). Environmental models extend time-dependent models to account for potential dependencies between diversification and measured environmental variables, for example, speciation and extinction rates can vary through time and both can be influenced by environmental variables. We focus here on mean global temperatures and Andean uplift. Climate change is probably one of the most important abiotic factors affecting biodiversity, of which global fluctuation in temperatures is the main component (129). In addition, the orogenesis of the Andes caused dramatic modifications in Neotropical landscapes and has become paradigmatic for explaining Neotropical biodiversity (8). Temperature variations during the Cenozoic were obtained from global compilations of deep-sea oxygen isotope ratios (δ^18^O) (72, 129), but we also analysed other curves to assess the impact of paleotemperature uncertainty on our results (see SI). For Andean paleo-elevations we retrieved a generalized model of the paleo-elevation history of the tropical Andes, compiled from several studies ((35) and references therein). The elevation of the Andes could have indirectly impacted the diversification of non-Andean groups. We thus applied uplift models to all clades in our study.

We fitted three environmental models in which speciation and/or extinction vary continuously with temperature changes *(λ[T]* and *μ[T])*, and three others with the elevation of the Andes *(λ[A]* and *μ[A])*. In this case, *λ*_*0*_ *(μ*_*0*_) is the expected speciation (extinction) rate under a temperature of 0°C (or a paleo-elevation of 0 m for the uplift models). We also analysed whether the speciation *(α)* and extinction *(μ)* dependency were positive or negative. For temperature models, *α(μ)* > 0 reflects increasing speciation (extinction) with increasing temperatures, and conversely. For the uplift models, *α(μ)* > 0 reflects increasing speciation (extinction) with increasing Andean elevations, and conversely. We accounted for missing species for each clade in the form of sampling fraction *(ρ)* (78) and assessed the strength of support of the models by computing Akaike information criterion (AICc), ΔAICc, and Akaike weights (AICω) to select the best-fit model. We derived diversity dynamics (Sc.1–4) based on the inferred diversification trends according to **Fig. 1**.

#### b) Diversification trends based on pulled diversification rates

To gain further insights in Neotropical diversification (Q1), we explored congruent diversification models defined in terms of pulled diversification rates (PDR, *r*_*p*_), and pulled extinction rates (PER, *μ*_*p*_) (69, 75). Two models are congruent if they have the same *r*_*p*_ and the same product *ρλ*_*0*_, in which *ρ* is the sampling fraction and *λ*_*0*_ *= λ(0). r*_*p*_ is equal to the net diversification rate (*r = λ* − *μ*) whenever *λ* is constant in time (*dλ/dτ = 0*), but differs from *r* when *λ* varies with time. The PER *μ*_*p*_ is equal to the extinction rate *μ* if *λ* is time-independent, but differs from *μ* in most other cases. Pulled and canonical diversification parameters are thus not equivalent in most cases. Biological interpretation of pulled parameters is not obvious. However, some specific properties of PDR and PER allowed us to compare diversification dynamics estimated based on pulled and canonical diversification parameters. Specifically, changes in speciation and/or extinction rates usually lead to similarly strong changes in PDR, while constant PDR are strong indicators that both *λ* and *μ* were constant or varied only slowly over time (69, 75). PDR can also yield other valuable insights: if *μ*_*p*_*(0)* is negative, this is evidence that speciation is currently decreasing over time (69, 75).

We estimated PDR values using the homogenous birth-death model on the R package *castor* 1.5.7 with the function *fit_hbd_pdr_on_grid* (130). We compared constant models (1 time interval) with models in which PDR values are allowed to vary independently on a grid of 3 time intervals. We set up the age grid non-uniformly, for example, age points were placed closer together near the present (where information content is higher), and we selected the model that best explained the lineage-through-time (LTT) of the Neotropical time trees based on AIC. To avoid non-global local optima, we performed 20 independent fitting trials starting from a random choice of model parameters. The *fit_hbd_pdr_on_grid* function additionally provided estimates of *ρλo* values. Knowing *ρ, λ*_*0*_ could be derived as follows: *λ*_*0*_ = *λ*_*0*_*ρ/ρ*. Similarly, pulled extinction rates for each time interval could be derived as follows: *μp*:= *λ*_*0*_ – *rp*. We limited the estimates to time periods with >10 species, using the *oldest_age* function in *castor*, to avoid points in the tree close to the root, where estimation uncertainty is generally higher.

### Neotropical bioregionalization

We used a quantitative approach to identify geographic units of long-term Neotropical evolution. We divided the Neotropical region into 13 operational areas based on the WWF biome classification (100) and similar to other studies, e.g., (4, 95) – Amazonia, Atlantic Forest, Bahama-Antilles, Caatinga, Central Andes, Cerrado, Chaco, Chocó, Guiana Shield, Mesoamerica, the Northern Andes, temperate South America, and an “elsewhere” region (see SI for detailed definitions) – and assessed the distribution in these areas of the 12,512 species included in our 150 phylogenies. Georeferenced records were downloaded for each species through GBIF using the R package *rgbif* 0.9.9 (131). We removed points with precision below 100 km, entries with mismatched georeference and country, duplicates, points representing country capitals or centroids, using the R package *CoordinateCleaner* 1.0-7 (132). Then, we created 13 georeferenced polygons delimiting each operational area using the WWF terrestrial ecoregions annotated shapefile in QGIS, and species were assigned to each polygon according to coordinate observations using the R package *speciesgeocodeR* 1.0-4. GBIF records can result in an overestimation of widespread ranges (133), so species distributions were manually inspected for completeness and accuracy with reference to databases (AmphibiaWeb 2018, Uetz et al. 2018, GBIF.org 2018, IUCN 2018). Based on the number of species belonging to each phylogenetic clade in the 13 ecoregions, we created a species abundance table (number of species per region per clade) that formed the basis for subsequent analyses.

The number of species distributed in each region within each clade were transformed using Hellinger transformations to account for differences in species richness between clades, and the *Morisita-horn* distance metric was selected to quantify pairwise dissimilarities of regional assemblages using the R package *vegan* 2.5-7 (134). We used K-means cluster analyses to form groups of similar regional assemblages. We determined the optimum number of groups by the elbow method. We use the function *fviz_cluster* in the R package *factoextra* 1.0.7 (135) to visualize K-means clustering results using principal component analysis.

### Variation of diversification dynamics across taxa, environmental drivers, and biogeographic units

We classified each clade in our study according to their main taxonomic group (plant [*n=*66], mammal [*n=*12], bird [*n=*32], squamate [*n=*24], amphibian [*n=*16]), environmental correlate (as estimated above: time [*n=*17], temperature [*n=*40] or uplift [*n=*17]), species richness dynamic based on canonical diversification rates (as estimated above: Sc. 1 [*n=*76], Sc. 2 [*n=*30], Sc. 3 [*n=*31], Sc. 4 [*n=*13]), and species richness dynamic based on pulled diversification rates (constant speciation [*n=*83], and decreasing speciation [*n=*51]).

We also classified each clade into the biogeographic units identified above (see results): cluster 1 (including the Amazonia, Central Andes, Chocó, Guiana Shield, Mesoamerica, and Northern Andes, [*n=*97]), cluster 2 (Atlantic Forest, Caatinga, Cerrado, Chaco, and temperate South America, [*n=*10]), cluster 3 (Bahama-Antilles, [*n=*4]), cluster 4 (“elsewhere” region, [*n=*0]), or cluster 5 (Galapagos, [*n=*0]). Clades were assigned to a given cluster only if > 60% of the species appeared in the cluster, otherwise clades were classified as ‘mixed’ (*n=*39). We additionally classified clades according to the main elevational range of their constituent species (lowland [< 1000 m; *n=*42], montane [1000–3500 m; *n=*8], highland [> 3500 m; *n=*6], mixed [*n=*94]; see SI for detailed definitions). Note that for the elevation, in our dataset most clades fell into the mixed category, with montane species most often occurring within clades of lowland species, and rarely forming a clade of their own. To account for this pattern (and minimize the number of clades classified as “mixed”), we performed additional analyses pooling “lowland” and “montane” categories and considered a clade “mixed” only if contained species in lowlands, montane and highlands (lowland-montane [*n=*124], highland [*n=*6], mixed [*n=*20]).

We assessed the phylogenetic signal of each multi-categorical trait (i.e., biogeographic units, elevation, diversity dynamics, and environmental drivers) using the *δ* statistics (136) over a phylogeny including one tip for each of the 150 clades represented in this study. This tree was constructed using TimeTree (137). High *δ*-value indicates strong phylogenetic signal. *δ* can be arbitrarily large, and thus significance was evaluated by comparing inferred *δ*-values to the distribution of values when the trait was randomised along the phylogeny. We evaluated the phylogenetic signal of continuous traits (i.e., diversification *[r]*, speciation *[λ]*, and pulled diversification *[r*_*p*_] rates) using Blomberg’s K (138) in the R package *phytools* 0.7-80 (139). Since time-varying diversification curves are hardly summarized in a single value, comparisons of net diversification values are based on estimates derived from the constant-rate model.

As any continuous (K_*r*_=0.06, *p*=0.6; K_*λ*_=0.07, *p*=0.4; K_*rp*_=0.07, *p*=0.6) or multi-categorical trait displays phylogenetic signal (Fig. S10), suggesting that the distribution of trait values is not explained by the phylogeny itself, statistical tests were conducted without applying phylogenetic corrections to account for the non-independence of data points. Fisher’s exact test was used in the analysis of contingency tables, performing pairwise-comparison with corrections for multiple testing (140), and Kruskal-Wallis Tests for comparing means between groups.

We also tested the effect of clade age, size and sampling fraction on the preferred species-richness dynamic (Sc. 1–4) using a phylogenetic ANOVA in *phytools* with posthoc comparisons, checking if the residual error controlling for the main effects in the model and the tree were normally distributed. We applied phylogenetic corrections in this case because phylogenetic signal was detected for sampling fraction (K_sampling_=0.12, *p*=0.001) and crown age (K_age_=0.22, *p*=0.001), not for tree size (K_size_=0.49, *p*=0.9).

## Acknowledgments

We thank all researchers who shared their published data through databases or to us directly (Drs. Arevalo, Martins, Fortes Santos, Simon, Lohmann, Mendoza, Swenson, Erkens, van der Meijden, and Freitas). Drs. J. Muñoz, J. Lobo, I. Sanmartín, S. Louca, P. Manzano and M. Godefroid for invaluable comments on the manuscript and/or analyses. This work was funded by an “*Investissements d’Avenir*” grant managed by the Agence Nationale de la Recherche (CEBA, ref. ANR-10-LABX-25-01) and the ANR GAARAnti project (ANR-17-CE31-0009). A.S.M. was also supported by the *Atracción de Talento* project (2019-T1/AMB-12648) and a Juan de la Cierva grant (IJCI-2017-32301).

A.A. is supported by the Swedish Research Council, the Swedish Foundation for Strategic Research, and the Royal Botanic Gardens, Kew. O.A.P.E. is funded by the Swiss Orchid Foundation and the Sainsbury Orchid Fellowship at the Royal Botanic Gardens, Kew. G.C. is funded by a Natural Environment Research Council Independent Research Fellowship (NE/S014470/1). R.R. is funded by the Spanish Ministry of Science (PID2019-108109GB-I00, AEI/FEDER).

## References

1. P. H. Raven, et al., The distribution of biodiversity richness in the tropics. Sci. Adv. 6, eabc6228 (2020).

2. R. A. Mittermeier, W. R. Turner, F. W. Larsen, T. M. Brooks, C. Gascon, “Global biodiversity conservation: the critical role of hotspots” in Biodiversity Hotspots, 1st Ed., F. E. Zachos, J. C. Habel, Eds. (Springer-Verlag, 2011), pp. 3–22.

3. N. Myers, R. A. Mittermeler, C. G. Mittermeler, G. A. B. Da Fonseca, J. Kent, Biodiversity hotspots for conservation priorities. Nature 403, 853–858 (2000).

4. A. Antonelli, et al., Amazonia is the primary source of Neotropical biodiversity. Proc. Natl. Acad. Sci. U. S. A. 115, 6034–6039 (2018).

5. A. H. Gentry, Neotropical floristic diversity: phytogeographical connections between Central and South America, Pleistocene climatic fluctuations, or an accident of the Andean orogeny? Ann. - Missouri Bot. Gard. 69, 557–593 (1982).

6. G. G. Simpson, Splendid isolation: the curious history of South American mammals (Yale University Press, 1980).

7. A. Antonelli, I. Sanmartín, Why are there so many plant species in the Neotropics? Taxon 60, 403–414 (2011).

8. C. Hoorn, et al., Amazonia through time: Andean uplift, climate change, landscape evolution, and biodiversity. Science 330, 927–931 (2010).

9. V. Rull, Neotropical biodiversity: Timing and potential drivers. Trends Ecol. Evol. 26, 508–513 (2011).

10. L. Diels, Pflanzengeographie (GJ Göschen, 1908).

11. J. W. Bews, Studies in the ecological evolution of the angiosperms. New Phytol. 26, 1–21 (1927).

12. A. Cronquist, The evolution and classification of flowering plants. (Thomas Nelson & Sons Ltd., 1968).

13. G. L. Stebbins, Flowering plants: evolution above the species level (Harvard University Press, 1974).

14. A. R. Wallace, Tropical nature, and other essays (Macmillan, 1878).

15. D. D. McKenna, B. D. Farrell, Tropical forests are both evolutionary cradles and museums of leaf beetle diversity. Proc. Natl. Acad. Sci. 103, 10947–10951 (2006).

16. J. Hey, Using phylogenetic trees to study speciation and extinction. Evolution 46, 627–640 (1992).

17. F. L. Condamine, N. S. Nagalingum, C. R. Marshall, H. Morlon, Origin and diversification of living cycads: A cautionary tale on the impact of the branching process prior in Bayesian molecular dating. BMC Evol. Biol. 15, 65 (2015).

18. A. Antonelli, I. Sanmartín, Mass extinction, gradual cooling, or rapid radiation? Reconstructing the spatiotemporal evolution of the ancient angiosperm genus Hedyosmum (Chloranthaceae) using empirical and simulated approaches. Syst. Biol. 60, 596–615 (2011).

19. G. C. Gibb, et al., Shotgun mitogenomics provides a reference phylogenetic framework and timescale for living xenarthrans. Mol. Biol. Evol. 33, 621–642 (2016).

20. C. Hoorn, J. Guerrero, G. A. Sarmiento, M. A. Lorente, Andean tectonics as a cause for changing drainage patterns in Miocene northern South America. Geology 23, 237–240 (1995).

21. C. Jaramillo, M. J. Rueda, G. Mora, Cenozoic plant diversity in the Neotropics. Science 311, 1893–1896 (2006).

22. P.-O. Antoine, R. Salas-Gismondi, F. Pujos, M. Ganerød, L. Marivaux, Western Amazonia as a Hotspot of Mammalian Biodiversity Throughout the Cenozoic. J. Mamm. Evol. 24, 5–17 (2017).

23. T. L. Couvreur, et al., Early evolutionary history of the flowering plant family Annonaceae: steady diversification and boreotropical geodispersal. J. Biogeogr. 38, 664–680 (2011).

24. E. P. Derryberry, et al., Lineage diversification and morphological evolution in a large-scale continental radiation: The Neotropical ovenbirds and woodcreepers (aves: furnariidae). Evolution 65, 2973–2986 (2011).

25. J. C. Santos, L. A. Coloma, K. Summers, Amazonian amphibian diversity is primarily derived from late Miocene Andean lineages. PLoS Biol. 7, e1000056 (2009).

26. R. J. Schley, et al., Is Amazonia a ‘museum’for Neotropical trees? The evolution of the Brownea clade (Detarioideae, Leguminosae). Mol. Phylogenet. Evol. 126, 279–292 (2018).

27. M. G. Harvey, et al., The evolution of a tropical biodiversity hotspot. Science 370, 1343–1348 (2020).

28. A. Antonelli, et al., Geological and climatic influences on mountain biodiversity. Nat. Geosci. 11, 718–725 (2018).

29. J. Haffer, Speciation in Amazon forest birds. Science 165, 131–137 (1969).

30. J.-E. Richardson, T.-D. Pennington, P.-M. Hollingsworth, Rapid diversification of a species-rich genus of Neotropical rain forest trees. Science 293, 2242–2245 (2001).

31. R. H. J. Erkens, L. W. Chatrou, J. W. Maas, T. van der Niet, V. Savolainen, A rapid diversification of rainforest trees (Guatteria; Annonaceae) following dispersal from Central into South America. Mol. Phylogenet. Evol. 44, 399–411 (2007).

32. C. Hughes, R. Eastwood, Island radiation on a continental scale: Exceptional rates of plant diversification after uplift of the Andes. Proc. Natl. Acad. Sci. U. S. A. 103, 10334–10339 (2006).

33. D. Esquerré, I. G. Brennan, R. A. Catullo, F. Torres-Pérez, J. S. Keogh, How mountains shape biodiversity: The role of the Andes in biogeography, diversification, and reproductive biology in South America’s most species-rich lizard radiation (Squamata: Liolaemidae). Evolution 73, 214–230 (2019).

34. C. S. Drummond, R. J. Eastwood, S. T. S. Miotto, C. E. Hughes, Multiple continental radiations and correlates of diversification in Lupinus (Leguminosae): testing for key innovation with incomplete taxon sampling. Syst. Biol. 61, 443–460 (2012).

35. L. P. Lagomarsino, F. L. Condamine, A. Antonelli, A. Mulch, C. C. Davis, The abiotic and biotic drivers of rapid diversification in Andean bellflowers (Campanulaceae). New Phytol. 210, 1430–1442 (2016).

36. O.A. Pérez-Escobar, et al., Recent origin and rapid speciation of Neotropical orchids in the world’s richest plant biodiversity hotspot. New Phytol. 215, 891–905 (2017).

37. L. J. Musher, M. Ferreira, A. L. Auerbach, J. McKay, J. Cracraft, Why is Amazonia a ‘source’of biodiversity? Climate-mediated dispersal and synchronous speciation across the Andes in an avian group (Tityrinae). Proc. R. Soc. B 286, 20182343 (2019).

38. M. Olave, L. J. Avila, J. W. Sites, M. Morando, How important is it to consider lineage diversification heterogeneity in macroevolutionary studies? Lessons from the lizard family Liolaemidae. J. Biogeogr. 47, 1286–1297 (2020).

39. D. L. Rabosky, Ecological limits and diversification rate: alternative paradigms to explain the variation in species richness among clades and regions. Ecol. Lett. 12, 735–743 (2009).

40. H. V Cornell, Is regional species diversity bounded or unbounded? Biol. Rev. 88, 140–165 (2013).

41. H. Morlon, M. D. Potts, J. B. Plotkin, Inferring the dynamics of diversification: a coalescent approach. PLOS Biol. 8, e1000493 (2010).

42. F. L. Condamine, J. Rolland, H. Morlon, Assessing the causes of diversification slowdowns: temperature-dependent and diversity-dependent models receive equivalent support. Ecol. Lett. 22, 1900–1912 (2019).

43. A. B. Phillimore, T. D. Price, Density-dependent cladogenesis in birds. PLOS Biol. 6, e71 (2008).

44. P. V. A. Fine, F. Zapata, D. C. Daly, Investigating processes of Neotropical rain forest tree diversification by examining the evolution and historical biogeography of the Protieae (Burseraceae). Evolution 68, 1988–2004 (2014).

45. C. D. Cadena, Testing the role of interspecific competition in the evolutionary origin of elevational zonation: an example with Buarremon brush-finches (Aves, Emberizidae) in the Neotropical mountains. Evol. Int. J. Org. Evol. 61, 1120–1136 (2007).

46. J. T. Weir, Divergent timing and patterns of species accumulation in lowland and highland neotropical birds. Evolution 60, 842–855 (2006).

47. A. S. Meseguer, F. L. Condamine, Ancient tropical extinctions at high latitudes contributed to the latitudinal diversity gradient. Evolution (2020).

48. T. B. Quental, C. R. Marshall, How the Red Queen Drives Terrestrial Mammals to Extinction. Science 341, 290 LP –292 (2013).

49. M. Foote, et al., Rise and fall of species occupancy in Cenozoic fossil mollusks. Science 318, 1131–1134 (2007).

50. S. B. Archibald, W. H. Bossert, D. R. Greenwood, B. D. Farrell, Seasonality, the latitudinal gradient of diversity, and Eocene insects. Paleobiology 36, 374–398 (2010).

51. R. Salas-Gismondi, et al., A Miocene hyperdiverse crocodylian community reveals peculiar trophic dynamics in proto-Amazonian mega-wetlands. Proc. R. Soc. B Biol. Sci. 282, 20142490 (2015).

52. S. A. Jansa, F. K. Barker, R. S. Voss, The early diversification history of didelphid marsupials: A window into South America’s “splendid isolation.” Evolution 68, 684–695 (2014).

53. J. D. Carrillo, et al., Disproportionate extinction of South American mammals drove the asymmetry of the Great American Biotic Interchange. Proc. Natl. Acad. Sci. 117, 26281–26287 (2020).

54. P. Wilf, et al., Eocene plant diversity at Laguna del Hunco and Río Pichileufú, Patagonia, Argentina. Am. Nat. 165, 634–650 (2005).

55. H. Kreft, W. Jetz, Global patterns and determinants of vascular plant diversity. Proc. Natl. Acad. Sci. 104, 5925–5930 (2007).

56. W. Jetz, G. H. Thomas, J. B. Joy, K. Hartmann, A. O. Mooers, The global diversity of birds in space and time. Nature 491, 444–448 (2012).

57. T. F. Rangel, et al., Modeling the ecology and evolution of biodiversity: biogeographical cradles, museums, and graves. Science 361, eaar5452 (2018).

58. I. Quintero, W. Jetz, Global elevational diversity and diversification of birds. Nature 555, 246–250 (2018).

59. T. N. C. Vasconcelos, et al., Fast diversification through a mosaic of evolutionary histories characterizes the endemic flora of ancient Neotropical mountains. Proc. R. Soc. B 287, 20192933 (2020).

60. C. Pouchon, et al., Phylogenomic analysis of the explosive adaptive radiation of the Espeletia Complex (Asteraceae) in the Tropical Andes. Syst. Biol. 67, 1041–1060 (2018).

61. S. Madriñán, A. Cortés, J. Richardson, Páramo is the world’s fastest evolving and coolest biodiversity hotspot. Front. Genet. 4, 192 (2013).

62. O. M. Vargas, B. Goldston, D. L. Grossenbacher, K. M. Kay, Patterns of speciation are similar across mountainous and lowland regions for a Neotropical plant radiation (Costaceae: Costus). Evolution 74, 2644–2661 (2020).

63. B. T. Smith, et al., The drivers of tropical speciation. Nature 515, 406–409 (2014).

64. W. L. Eiserhardt, T. L. P. Couvreur, W. J. Baker, Plant phylogeny as a window on the evolution of hyperdiversity in the tropical rainforest biome. New Phytol. 214, 1408–1422 (2017).

65. S. Nee, E. C. Holmes, R. M. May, P. H. Harvey, Extinction rates can be estimated from molecular phylogenies. Philos. Trans. - R. Soc. London, B 344, 77–82 (1994).

66. D. L. Rabosky, Extinction rates should not be estimated from molecular phylogenies. Evolution 6, 1816–1824 (2010).

67. T. Stadler, How can we improve accuracy of macroevolutionary rate estimates? Syst. Biol. 62, 321–329 (2013).

68. G. Burin, L. R. V Alencar, J. Chang, M. E. Alfaro, T. B. Quental, How well can we estimate diversity dynamics for clades in diversity decline? Syst. Biol. 68, 47–62 (2018).

69. S. Louca, M. W. Pennell, Extant timetrees are consistent with a myriad of diversification histories. Nature 580, 502–505 (2020).

70. H. Morlon, F. Hartig, S. Robin, Prior hypotheses or regularization allow inference of diversification histories from extant timetrees. bioRxiv doi:10.1101/2020.07.03.185074 (2020) https://doi.org/10.1101/2020.07.03.185074.

71. A. S. Meseguer, P. Antoine, A. Fouquet, F. Delsuc, F. L. Condamine, The role of the Neotropics as a source of world tetrapod biodiversity. Glob. Ecol. Biogeogr. 29, 1565–1578 (2020).

72. J. C. Zachos, G. R. Dickens, R. E. Zeebe, An early Cenozoic perspective on greenhouse warming and carbon-cycle dynamics. Nature 451, 279–283 (2008).

73. J. Hansen, M. Sato, G. Russell, P. Kharecha, Climate sensitivity, sea level and atmospheric carbon dioxide. Philos. Trans. R. Soc. A Math. Phys. Eng. Sci. 371, 20120294 (2013).

74. J. Veizer, A. Prokoph, Temperatures and oxygen isotopic composition of Phanerozoic oceans. Earth-Science Rev. 146, 92–104 (2015).

75. S. Louca, et al., Bacterial diversification through geological time. Nat. Ecol. Evol. 2, 1458–1467 (2018).

76. M. A. McPeek, The ecological dynamics of clade diversification and community assembly. Am. Nat. 172, E270–E284 (2008).

77. V. E. Luzuriaga-Aveiga, J. T. Weir, Elevational differentiation accelerates trait evolution but not speciation rates in Amazonian birds. Ecol. Lett. 22, 624–633 (2019).

78. H. Morlon, T. L. Parsons, J. B. Plotkin, Reconciling molecular phylogenies with the fossil record. Proc. Natl. Acad. Sci. 108, 16327–16332 (2011).

79. M. P. Davis, P. E. Midford, W. Maddison, Exploring power and parameter estimation of the BiSSE method for analyzing species diversification. BMC Evol. Biol. 13, 38 (2013).

80. N. Cusimano, S. Renner, Slowdowns in diversification rates from real phylogenies may not be real. Syst. Biol. 59, 458–464 (2010).

81. O. Hagen, A. Skeels, R. E. Onstein, W. Jetz, L. Pellissier, Earth history events shaped the evolution of uneven biodiversity across tropical moist forests. Proc. Natl. Acad. Sci. 118 (2021).

82. M. S. Springer, et al., Macroevolutionary dynamics and historical biogeography of primate diversification inferred from a species supermatrix. PLoS One 7, e49521 (2012).

83. R. S. Etienne, et al., Diversity-dependence brings molecular phylogenies closer to agreement with the fossil record. Proc. R. Soc. London B 279, 1300–1309 (2012).

84. D. L. Rabosky, S. C. Donnellan, M. Grundler, I. J. Lovette, Analysis and visualization of complex macroevolutionary dynamics: an example from Australian scincid lizards. Syst. Biol. 63, 610–627 (2014).

85. H. Morlon, Phylogenetic approaches for studying diversification. Ecol. Lett. 17, 508–525 (2014).

86. T. Pannetier, C. Martinez, L. Bunnefeld, R. S. Etienne, Branching patterns in phylogenies cannot distinguish diversity-dependent diversification from time-dependent diversification. Evolution 75, 25–38 (2021).

87. D. Moen, H. Morlon, Why does diversification slow down? Trends Ecol. Evol. 29, 190–197 (2014).

88. J. F. Gillooly, J. H. Brown, G. B. West, V. M. Savage, E. L. Charnov, Effects of size and temperature on metabolic rate. Science 293, 2248–2251 (2001).

89. A. P. Allen, J. F. Gillooly, V. M. Savage, J. H. Brown, Kinetic effects of temperature on rates of genetic divergence and speciation. Proc. Natl. Acad. Sci. 103, 9130–9135 (2006).

90. P. J. Mayhew, M. A. Bell, T. G. Benton, A. J. McGowan, Biodiversity tracks temperature over time. Proc. Natl. Acad. Sci. 109, 15141–15145 (2012).

91. G. Chomicki, M. Weber, A. Antonelli, J. Bascompte, E. T. Kiers, The Impact of Mutualisms on Species Richness. Trends Ecol. Evol. 34, 698–711 (2019).

92. A. Fouquet, C. Santana Cassini, C. Fernando Baptista Haddad, N. Pech, M. Trefaut Rodrigues, Species delimitation, patterns of diversification and historical biogeography of the Neotropical frog genus Adenomera (Anura, Leptodactylidae). J. Biogeogr. 41, 855–870 (2014).

93. D. S. Moen, J. J. Wiens, Microhabitat and climatic niche change explain patterns of diversification among frog families. Am. Nat. 190, 29–44 (2017).

94. K. H. Kozak, J. J. Wiens, Accelerated rates of climatic-niche evolution underlie rapid species diversification. Ecol. Lett. 13, 1378–1389 (2010).

95. C. R. Hutter, S. M. Lambert, J. J. Wiens, Rapid diversification and time explain Amphibian richness at different scales in the tropical Andes, Earth’s most biodiverse Hotspot. Am. Nat. 190, 828–843 (2017).

96. T. A. Kursar, et al., The evolution of antiherbivore defenses and their contribution to species coexistence in the tropical tree genus Inga. Proc. Natl. Acad. Sci. 106, 18073–18078 (2009).

97. I. Sanmartín, F. Ronquist, Southern Hemisphere biogeography inferred by event-based models: Plant versus animal patterns. Syst. Biol. 53, 216–243 (2004).

98. A. F. Magee, S. Höhna, T. I. Vasylyeva, A.D. Leaché, V. N. Minin, Locally adaptive Bayesian birth-death model successfully detects slow and rapid rate shifts. PLoS Comput. Biol. 16, e1007999 (2020).

99. H. Walter, E. Box, Global classification of natural terrestrial ecosystems. Vegetatio 32, 75–81 (1976).

100. D. M. Olson, et al., Terrestrial Ecoregions of the World: A New Map of Life on Earth. Bioscience 51, 933 (2001).

101. T. Escalante, J. J. Morrone, G. Rodríguez-Tapia, Biogeographic regions of North American mammals based on endemism. Biol. J. Linn. Soc. 110, 485–499 (2013).

102. J. J. Morrone, Biogeographical regionalisation of the Neotropical region. Zootaxa 3782, 1–110 (2014).

103. B. G. Holt, et al., An update of Wallace’s zoogeographic regions of the world. Science 339, 74–78 (2013).

104. H. Kreft, W. Jetz, A framework for delineating biogeographical regions based on species distributions. J. Biogeogr. 37, 2029–2053 (2010).

105. D. E. Prado, P. E. Gibbs, Patterns of species distributions in the dry seasonal forests of South America. Ann. Missouri Bot. Gard. 80, 902–927 (1993).

106. F. Luebert, The two South American dry diagonals. Front. Biogeogr. 13.4, e51267 (2021).

107. R. T. Pennington, J. E. Richardson, M. Lavin, Insights into the historical construction of species-rich biomes from dated plant phylogenies, neutral ecological theory and phylogenetic community structure. New Phytol. 172, 605–616 (2006).

108. D. J. Beerling, C. P. Osborne, The origin of the savanna biome. Glob. Chang. Biol. 12, 2023–2031 (2006).

109. J. X. Becerra, Timing the origin and expansion of the Mexican tropical dry forest. Proc. Natl. Acad. Sci. 102, 10919–10923 (2005).

110. H. Kreft, W. Jetz, Comment on “An update of Wallace’s zoogeographic regions of the world.” Science 341, 343 (2013).

111. C. Rahbek, et al., Building mountain biodiversity: Geological and evolutionary processes. Science 365, 1114–1119 (2019).

112. R. Armijo, R. Lacassin, A. Coudurier-Curveur, D. Carrizo, Coupled tectonic evolution of Andean orogeny and global climate. Earth-Science Rev. 143, 1–35 (2015).

113. P. M. Blisniuk, L. A. Stern, C. P. Chamberlain, B. Idleman, P. K. Zeitler, Climatic and ecologic changes during Miocene surface uplift in the Southern Patagonian Andes. Earth Planet. Sci. Lett. 230, 125–142 (2005).

114. S. G. A. Flantua, A. O’dea, R. E. Onstein, C. Giraldo, H. Hooghiemstra, The flickering connectivity system of the north Andean páramos. J. Biogeogr. 46, 1808–1825 (2019).

115. T. H. G. Ezard, T. B. Quental, M. J. Benton, The challenges to inferring the regulators of biodiversity in deep time. Philos. Trans. R. Soc. B-Biological Sci. 371, 20150216 (2016).

116. A. J. Helmstetter, et al., Pulled diversification rates, lineage-through-time plots and modern macroevolutionary modelling. bioRxiv (2021).

117. T. L. P. Couvreur, Odd man out: why are there fewer plant species in African rain forests? Plant Syst. Evol. 301, 1299–1313 (2015).

118. A. Antonelli, The rise and fall of Neotropical biodiversity. Bot. J. Linnaean Soc. (in press) (2021).

119. R. A. Pyron, F. T. Burbrink, J. J. Wiens, A phylogeny and revised classification of Squamata, including 4161 species of lizards and snakes. BMC Evol. Biol. 13, 93 (2013).

120. R. A. Pyron, Biogeographic analysis reveals ancient continental vicariance and recent oceanic dispersal in amphibians. Syst. Biol. 63, 779–797 (2014).

121. R. A. Pyron, F. T. Burbrink, Early origin of viviparity and multiple reversions to oviparity in squamate reptiles. Ecol. Lett. 17, 13–21 (2014).

122. O. R. P. Bininda-Emonds, et al., The delayed rise of present-day mammals. Nature 446, 507–512 (2007).

123. T. S. Kuhn, A. Mooers, G. H. Thomas, A simple polytomy resolver for dated phylogenies. Methods Ecol. Evol. 2, 427–436 (2011).

124. A. E. Zanne, et al., Three keys to the radiation of angiosperms into freezing environments. Nature 506, 89–92 (2014).

125. M. Töpel, et al., SpeciesGeoCoder: Fast categorization of species occurrences for analyses of biodiversity, biogeography, ecology, and evolution in Systematic Biology, (2017), pp. 145–151.

126. R. Jansson, G. Rodríguez-Castañeda, L. E. Harding, What can multiple phylogenies say about the latitudinal diversity gradient? A new look at the tropical conservatism, out of the tropics, and diversification rate hypotheses. Evolution 67, 1741–1755 (2013).

127. H. Morlon, et al., RPANDA: an R package for macroevolutionary analyses on phylogenetic trees. Methods Ecol. Evol. 7, 589–597 (2016).

128. F. L. Condamine, J. Rolland, H. Morlon, Macroevolutionary perspectives to environmental change. Ecol. Lett. 16, 72–85 (2013).

129. A. Prokoph, G. A. Shields, J. Veizer, Compilation and time-series analysis of a marine carbonate δ18O, δ13C, 87Sr/86Sr and δ34S database through Earth history. Earth-Science Rev. 87, 113–133 (2008).

130. S. Louca, M. Doebeli, Efficient comparative phylogenetics on large trees. Bioinformatics 34, 1053–1055 (2018).

131. S. Chamberlain, K. Ram, V. Barve, D. Mcglinn, M. S. Chamberlain, Package ‘rgbif’ (2017).

132. A. Zizka, et al., CoordinateCleaner: Standardized cleaning of occurrence records from biological collection databases. Methods Ecol. Evol. 10, 744–751 (2019).

133. C. Maldonado, et al., Estimating species diversity and distribution in the era of big data: to what extent can we trust public databases? Glob. Ecol. Biogeogr. 24, 973–984 (2015).

134. J. Oksanen, et al., Package ‘vegan.’ Community Ecol. Packag. version 2, 1–295 (2013).

135. A. Kassambara, F. Mundt, Package ‘factoextra.’ Extr. Vis. results Multivar. data Anal. 76 (2017).

136. R. Borges, J. P. Machado, C. Gomes, A. P. Rocha, A. Antunes, Measuring phylogenetic signal between categorical traits and phylogenies. Bioinformatics 35, 1862–1869 (2019).

137. S. Kumar, G. Stecher, M. Suleski, S. B. Hedges, TimeTree: a resource for timelines, timetrees, and divergence times. Mol. Biol. Evol. 34, 1812–1819 (2017).

138. S. P. Blomberg, T. Garland Jr, A. R. Ives, Testing for phylogenetic signal in comparative data: behavioral traits are more labile. Evolution 57, 717–745 (2003).

139. L. J. Revell, phytools: an R package for phylogenetic comparative biology (and other things). Methods Ecol. Evol. 3, 217–223 (2012).

140. Y. Benjamini, Y. Hochberg, Controlling the false discovery rate: a practical and powerful approach to multiple testing. JR Stat. Soc. B 57, 289–300 (1995).

