## Supplementary Information for "The Origins and Drivers of Neotropical Diversity"

**This PDF file includes:**

Supporting Information (SI) text

Supporting Information Tables: S1 to S13

References for SI text

Supporting Information Figures: S1 to S11

### Supporting Information (SI) text

***Mean global paleotemperatures.*** Recent investigations derived different paleotemperature curves for the Cenozoic (1–3). The estimates mostly differing in the magnitude of the temperature changes but sharing the same overall trend (Fig. S4; Table S12). To account for the uncertainty on global paleotemperatures on our results, we performed diversification analyses in RPANDA using three different global temperature curves; ***(i)*** the global temperature curve estimated by Prokoph *et al.* (4), Veizer & Prokoph (3), and Zachos *et al.* (5, 6) that used deep-sea oxygen benthic foraminifera (bf) $\delta^{18}O_{bf}$ isotopic ratio to provide estimates for the last 540 million years (Myrs), thus spanning the full time range over which Neotropical lineages diversified; ***(ii)*** the temperature curve by Cramer *et al.* (2, 7), which is similar to the more widely used previous curve but accounts for fluctuations in sea water (sw) $\delta^{18}O_{sw}$ through time and correct for ice volume. This curve provides temperature estimates for the last 62.4 Myrs; and ***(iii)*** the paleotemperature curve estimated by Hanssen *et al.* (1) for the last 65.6 Myrs, which accounts for ice volume and deep ocean temperature changes, and provides estimates of surface and deep-water temperature changes. All these curves reflect planetary-scale climatic trends that are expected to have led to temporally coordinated changes in Neotropical lineages. We compare diversification models based on the different paleotemperature estimates only for groups overlapping the isotope record of the tree paleotemperature curves (< 62.4 Myrs; resulting in 128 phylogenies).

We found that diversification analyses based on the different paleotemperature curves produced almost identical results, in terms of model selection, parameter estimates and main diversification trends (Fig. S4). Therefore, we present and discuss in the main text the results based on the curve of Veizer & Prokoph (3), as this is the only curve spanning the full time range of all the Neotropical lineages included in our dataset (150 phylogenies).

***Bioregionalization analyses.*** The Neotropical region was divided into 13 operational areas based on the WWF biome classification (8) (Fig. 6) as follows:

1. North America: Nearctic regions ranging from north and central Mexico through the southern US and north through Canada.
2. Mesoamerica: Panama through the Neotropical pine-oak forests, eastern Sinaloan dry forests, and western Veracruz moist forests of Mexico.
3. Bahamian-Antilles: From the Everglades in the southern tip of Florida south through the Caribbean Islands to Grenada.
4. Chocó: encompassing the moist and dry forests of the Chocó-Darien, Ecuador, and Magdalena-Uraba, western Pacific and northern Atlantic Colombian mangroves, and northern Colombian xeric scrub.
5. Northern Andes: Northern Andean Páramo and montane forests of the Eastern Cordillera Real, Magdalena Valley, Cordillera Oriental, and Venezuelan Andes.
6. Central Andes: western Peru through southern Bolivia, northern Chile, and the northeast Andes of Argentina, Peruvian Yungas, Páramo, and wet and dry Puna and the Sechura and Atacama Deserts.
7. Temperate South America: From the Chilean Matorral and Low Monte in the east, bordered by the dry central Chaco, to the dry savannahs of Uruguay in the west, and south to the southern tip of South America.
8. Guiana Shield: Guianan moist forests, Guianan savannah, Llanos, and nearby dry forests and xeric shrublands to Atlantic mangroves in the north and Trinidad and Tobago.

9. Amazonia: moist forests of the Amazon basin, bordered by the Northern and Central Andes in the east and Guiana Shield in the north, across to the Atlantic coast, bordered in the south and southeast by the dry regions of the Caatinga and Cerrado.
10. Cerrado: grasslands and dry forests of eastern Brazil, including the Pantanal savannah and Chiquitano forest.
11. Caatinga: grasslands and dry forests of north-eastern Brazil to the coast of the Atlantic Ocean.
12. Atlantic Forest: eastern Brazilian coast into the moist forests of Bahia, Alta Parana, Araucaria, and along the coast at Serra do Mar, almost to the Uruguayan border in the south.
13. Chaco: Argentinian, Bolivian, and Paraguayan dry and humid Chaco and Parana flooded savannah, north of the temperate grasslands.

***Diversification correlates.*** We evaluated if diversification trends estimated in this study (*i.e.* increasing, constant, decreasing) differ between lineages characterized by different geographic, and elevations. ***(1)*** For the elevation ranges, we consider three dominant patterns, and classified the phylogenies according to the elevation zone in which the species of the clades mainly occur, although single species in large clades may be exceptions: **lowland** = 0-1000 m, including lowland rainforest in Amazonia and the Chocó in western Colombia and Ecuador, as well as rainforest in the flanking lowland and pre-montane areas along the eastern side of the Andes; **montane** = 1,000-3,500 m, including mid-elevation montane forests (*e.g.* cloud and elfin forests); and **highland** = >3,500 m, including alpine-altitude grasslands (Páramo). **Mixed** category includes lineages that show a mixed preference between lowland, montane and highland.

***Phylogenetic reconstruction and dating of Caviomorpha***

*Molecular dataset and sequencing.* We obtained a time-calibrated species-level phylogeny for Caviomorpha following the Woods and Kilpatrick (9) taxonomic list, and including recent taxonomic updates for newly described genera and species (10). Five molecular markers proven valuable to resolve caviomorph phylogenies were considered in this study (11–13): two mitochondrial genes (cytochrome *b* apoenzyme: *cyt b* and 12S ribosomal unit: *12S rRNA*) and three nuclear genes (growth hormone receptor exon 10: *GHR*; interphotoreceptor retinoid binding protein exon 1: *IRPB,* and Recombinating Activating protein 1: *RAG1*). DNA cytb and *12S rRNA* sequences were newly generated for the following species: *Proechimys brevicauda, Proechimys cuvieri, Proechimys echinothrix, Proechimys gardneri, Proechimys goeldii, Proechimys gorgonae, Proechimys gularis, Proechimys guyannensis, Proechimys hoplomyoides, Proechimys kulinae, Proechimys longicaudatus, Proechimys mincae, Proechimys pattoni, Proechimys poliopus, Proechimys quadruplicatus, Proechimys roberti, Proechimys semispinosus, Proechimys simonsi, Proechimys steerei, Proechimys trinitatus, Proechimys warreni,*, *Diplomys caniceps* and *Olallamys edax*. Ctenohystrica sequences from previous studies were downloaded from public databanks (Table S13). The final dataset includes 199 caviomorpha species representing 82% of the species and all the genera described (9). We included 37 representatives of Diatomyidae, Ctenodactylidae, Hystricidae, Heterocephalidae, Bathyergidae, Petromuridae and Thryonomyidae, Pedetidae, Dipodidae and Muridae as outgroups (14–16).

Fresh tissues were extracted using the QIAGEN extraction kit protocol. Four museum skin samples of rare unsequenced species (*Proechimys gorgonae, Proechimys trinitatus, Diplomys caniceps* and *Olallamys edax*) were stored in Eppendorf tubes. They were processed in the "Degraded DNA Facility" in Montpellier, France (dedicated to processing low quality/quantity DNA tissue samples). These samples were extracted using the

QUIAGEN extraction kit protocol, in small batches (of 3 samples maximum), and a negative control was included in each batch to monitor possible contamination. For all these samples, libraries were prepared following Tilak et al. (17) protocol, in order to obtain the complete mitochondrial genome. These seven libraries were pooled and sequenced without enrichment as single end reads on Illumina HiSeq 2000 lanes at the GATC-Biotech company (Konstanz, Germany). Raw 101-nt reads were imported in Geneious R6 (18) and adaptor fragments were removed by the "trim ends" utility. The mapping of the reads on the phylogenetically closest available mitochondrial genome was performed for each species. The following mapping parameters were used in Geneious read mapper: a minimum of 24 consecutive nucleotides (nt) perfectly matching the reference, a maximum 5% of single nt mismatch over the read length, a minimum of 95%-nt similarity in overlap region, and a maximum of 3% of gaps with a maximum gap size of 3-nt. Iterative mapping cycles were performed in order to elongate the sequence when the complete mitogenome was not recovered after the initial mapping round. A high-quality consensus was generated and the circularity of the mitogenome was verified by the exact superimposition of the 100-nt at the assembly extremities. We extracted the cytb and *12S rRNA* from these mitogenomes in order to perform our phylogenetic analyses We used SEAVIEW (19) and MACSE v2 (20) to align the sequences. They were translated into peptide sequences to exclude putative NUMt copies and to ensure sequence orthology. From these individual alignments, we built four gene matrices; *cyt b* (224 taxa and 1140 sites), *12S* rRNA (152 taxa and 775 sites), *GHR* (95 taxa and 927 sites), *RAG1* (63 taxa and 1064 sites), vWF (83 taxa and 1269 sites) and a nuclear + mitochondrial supermatrix (232 taxa and 5174 sites).

*Phylogenetic analyses.* Phylogenetic trees were reconstructed using a maximum likelihood (ML) method. ML analyses were first carried out on each marker independently and on the supermatrix using RAxML 8.0 (21, 22). Each gene considered separately does not result in a robust phylogeny for Caviomorpha: mitochondrial marker helps to resolve terminal nodes, while nuclear genes lend support to deepest ones. Since the 5 different markers yielded consistent, compatible topologies, sequences were concatenated and phylogenetic analyses were then carried out using the combined dataset. We used PartitionFinder v2.1.1 (23) to find the best partition schemes and nucleotide substitution models. Robustness of each gene tree was assessed using the rapid bootstrap (Bp) procedure (option –f a) with 1,000 replications (option -# numberOfRuns) (21, 22). Divergence times were subsequently estimated from the mitochondrial + nuclear nucleotide supermatrices to provide a temporal framework of the caviomorpha radiation. A Bayesian relaxed molecular clock method was used to estimate divergence dates whilst accounting for changes in evolutionary rates through time and allowing for independent models of sequence evolution for each gene partition. The best fitting substitution models for each partition were selected according to PartitionFinder results (23). We used Beast v1.10.2 (24) for phylogenetic analyses, assuming the Birth Death model of speciation and an uncorrelated log-normal distribution molecular clock as tree priors. Clock models were unlinked across 5 genes and codon partitions of the exons in order to account for missing data (25). We ran MCMC chains for 250 million generations, with trees sampled every 10,000 generations. We performed the analyses 4 times to check for convergence of model parameter estimates, and Tracer (26) was used to assess algorithm convergence. We removed the first 25% of trees as burn in. Trees from each of the 4 independent runs were combined into a maximum clade credibility tree with mean node heights calculated using TreeCombiner and TreeAnnotator. All these analyses were computed on the CIPRES science gateway. To calibrate the phylogeny we selected 17 fossil constraints as described from previous studies (11). In order to take into account uncertainties in the phylogenetic position of these fossils, all constraints were set using hard minimum bounds and soft upper bounds under a lognormal prior, as suggested by recent paleontological studies (27, 28).

### Supporting Tables

**Table S1**. Dataset of plant phylogenies, including the taxonomic level (Tax. level), the crown age (in million years ago), the number of species on the clade (#spp), the number of Neotropical species (#spp Neotrop) and proportion, as well as the sampling fraction (*i.e.* the number of species sampled in the tree from the total number of species described in the group).

| ID | Order | Family | Clade name | Tax. level | Crown Age | #spp | #spp Neotrop | % Neotrop | #spp sampled | Samplig fraction | Tree origin | #spp reference |
|---|---|---|---|---|---|---|---|---|---|---|---|---|
| P1 | Asparagales | Orchidaceae | *Prosthechea* | genus | 2,88 | 112 | 112 | 100,00 | 13 | 0,12 | (29) | (30) |
| P2 | Asparagales | Orchidaceae | *Camaridium* | genus | 0,93 | 140 | 140 | 100,00 | 39 | 0,28 | (29) | (30) |
| P3 | Zingiberales | Zingiberaceae | *Renealmia* | genus | 43,02 | 76 | 61 | 80,26 | 12 | 0,16 | (29) | (31) |
| P4 | Zingiberales | Costaceae | *Costus*_clade 1 | section | 13,8 | 80 | 40 | 50,00 | 10 | 0,13 | (29) | (32) |
| P5 | Zingiberales | Costaceae | *Costus*_clade 2 | section | 7,32 | 80 | 40 | 50,00 | 10 | 0,13 | (29) | (32) |
| P6 | Magnoliales | Annonaceae | *Cremastosperma* | genus | 5,81 | 31 | 31 | 100,00 | 14 | 0,45 | (29) | (33) |
| P7 | Magnoliales | Annonaceae | Annonaceae-clade-2 | genus | 12,27 | 51 | 51 | 100,00 | 29 | 0,57 | (29) | (33) |
| P8 | Myrtales | Onagraceae | *Fuchsia* | genus | 11,87 | 108 | 103 | 95,37 | 15 | 0,14 | (29) | (34) |
| P9 | Myrtales | Melastomataceae | Melastomataceae-clade-1 | tribe | 19,62 | 72 | 72 | 100,00 | 11 | 0,15 | (29) | (35) |
| P10 | Rosales | Moraceae | Castilleae group | tribe | 25,52 | 59 | 59 | 100,00 | 24 | 0,41 | (29) | (30, 36) |
| P11 | Fabales | Fabaceae | *Andira*-clade | genus | 15,15 | 44 | 43 | 97,73 | 30 | 0,68 | (37) | (37) |
| P12 | Caryophyllales | Polygonaceae | Polygonaceae-clade-1 | genus | 13,04 | 43 | 43 | 100,00 | 15 | 0,35 | (29) | (30) |
| P13 | Ericales | Sapotaceae | *Sideroxylon* | genus | 56,3 | 81 | 49 | 60,49 | 26 | 0,32 | (38) | (30) |
| P14 | Ericales | Lecythidaceae | Lecythidaceae-clade-1 | genus | 26 | 50 | 50 | 100,00 | 10 | 0,20 | (29) | (30) |
| P15 | Ericales | Lecythidaceae | *Lecythis* | genus | 19,87 | 27 | 27 | 100,00 | 17 | 0,63 | (29) | (30, 39) |
| P16 | Ericales | Lecythidaceae | *Eschweilera* | genus | 7 | 92 | 92 | 100,00 | 19 | 0,21 | (29) | (30, 39) |
| P17 | Gentianales | Rubiaceae | Condamineeae | genus | 22,42 | 99 | 99 | 100,00 | 18 | 0,18 | (29) | (30) |
| P18 | Fabales | Fabaceae | *Amicia* | genus | 10,8 | 8 | 8 | 100,00 | 7 | 0,88 | (40) | (30) |

| | | | | | | | | | | | | |
|---|---|---|---|---|---|---|---|---|---|---|---|---|
| P19 | Gentianales | Rubiaceae | Cinchoneideae-clade | Genus | 28,06 | 122 | 117 | 95,90 | 22 | 0,18 | (41) | (30) |
| P20 | Fabales | Fabaceae | *Coursetia* | Genus | 20,25 | 40 | 40 | 100,00 | 29 | 0,73 | (42) | (30) |
| P21 | Magnoliales | Annonaceae | *Guatteria* | genus | 20,76 | 254 | 254 | 100,00 | 103 | 0,41 | (43) | (30) |
| P22 | Chloranthales | Chlorantaceae | *Hedyosmum* | genus | 17,63 | 44 | 44 | 100,00 | 19 | 0,43 | (44) | (30) |
| P23 | Gentianales | Rubiaceae | Isertieae | tribe | 25,05 | 16 | 16 | 100,00 | 9 | 0,56 | (41) | (30) |
| P24 | Caryophyllales | Campanulaceae | Centropogonid_clade | tribe | 17 | 550 | 550 | 100,00 | 200 | 0,36 | (45) | (45) |
| P25 | Cycadales | Zamiaceae | *Ceratozamia* | genus | 19,2 | 27 | 27 | 100,00 | 24 | 0,89 | (46) | (46) |
| P26 | Cycadales | Zamiaceae | *Zamia* | genus | 14,63 | 71 | 71 | 100,00 | 43 | 0,61 | (46) | (46) |
| P27 | Fabales | Fabaceae | *Lupinus*-clade-1 | subgenus | 4,67 | 35 | 35 | 100,00 | 19 | 0,54 | (47) | (47) |
| P28 | Fabales | Fabaceae | *Lupinus*-clade-2 | subgenus | 2,75 | 130 | 130 | 100,00 | 37 | 0,28 | (47) | (47) |
| P29 | Lamales | Bignoniaceae | *Tynanthus* | genus | 15,28 | 15 | 15 | 100,00 | 14 | 0,93 | (48) | (30) |
| P30 | Arecales | Arecaceae | *Sabal* | genus | 37,32 | 16 | 16 | 100,00 | 14 | 0,88 | (49) | (30) |
| P31 | Arecales | Arecaceae | Cryosophileae | tribe | 30,96 | 80 | 80 | 100,00 | 78 | 0,98 | (49) | (30) |
| P32 | Arecales | Arecaceae | *Copernicia* | genus | 20,51 | 28 | 28 | 100,00 | 22 | 0,79 | (49) | (30) |
| P33 | Arecales | Arecaceae | *Livistoninae*-clade-1 | tribe | 19,96 | 16 | 16 | 100,00 | 16 | 1,00 | (49) | (30) |
| P34 | Arecales | Arecaceae | Iriarteeae | tribe | 29,67 | 32 | 32 | 100,00 | 32 | 1,00 | (49) | (30) |
| P35 | Arecales | Arecaceae | Chamaedoreeae | tribe | 38,07 | 115 | 115 | 100,00 | 114 | 0,99 | (49) | (30) |
| P36 | Arecales | Arecaceae | Attaleinae | tribe | 27,09 | 156 | 156 | 100,00 | 154 | 0,99 | (49) | (30) |
| P37 | Arecales | Arecaceae | Elaeidinae+Bactridinae | tribe | 43,62 | 180 | 180 | 100,00 | 178 | 0,99 | (49) | (30) |
| P38 | Arecales | Arecaceae | Geonomateae | tribe | 26,92 | 102 | 102 | 100,00 | 102 | 1,00 | (49) | (30) |
| P39 | Arecales | Arecaceae | Euterpeae | tribe | 34,2 | 22 | 22 | 100,00 | 22 | 1,00 | (49) | (30) |
| P40 | Fabales | Fabaceae | *Piptadenia*-group | genus | 20,39 | 594 | 554 | 93,27 | 271 | 0,46 | (37) | (50) |
| P41 | Asparagales | Orchidaceae | Pleurothallidinae | tribe | 18,34 | 5100 | 5100 | 100,00 | 670 | 0,13 | (51) | (51) |
| P42 | Asparagales | Orchidaceae | Cymbidieae | tribe | 34,7 | 3700 | 3330 | 90,00 | 789 | 0,21 | (51) | (51) |
| P43 | Poales | Poaceae (Panicoideaea) | Paspaleae | tribe | 21 | 680 | 680 | 100,00 | 168 | 0,25 | (52) | (30) |
| P44 | Fabales | Fabaceae | *Inga* | genus | 6 | 381 | 381 | 100,00 | 126 | 0,33 | (53) | (30) |
| P45 | Lamiales | Gesneriaceae | Gesnerioideae | tribe | 48,44 | 1200 | 1200 | 100,00 | 588 | 0,49 | (54) | (54) |
| P46 | Sapindales | Burseraceae | Proteiae | tribe | 28,91 | 140 | 134 | 95,71 | 111 | 0,79 | (55) | (55) |
| P47 | Solanales | Solanaceae | Schizanthoideae-Goetzeoideae | tribe | 28,8 | 21 | 20 | 95,24 | 21 | 1,00 | (56, 57) | (56, 57) |

| | | | | | | | | | | | | |
|---|---|---|---|---|---|---|---|---|---|---|---|---|
| P48 | Solanales | Solanaceae | Cestroideae | tribe | 32,21 | 221 | 221 | 100,00 | 50 | 0,23 | (56, 57) | (56, 57) |
| P49 | Solanales | Solanaceae | Petunioideae | tribe | 30,92 | 146 | 146 | 100,00 | 44 | 0,30 | (56, 57) | (56, 57) |
| P50 | Solanales | Solanaceae | Nolaneae | tribe | 7,2 | 89 | 89 | 100,00 | 63 | 0,71 | (56, 57) | (56, 57) |
| P51 | Solanales | Solanaceae | Physaleae | tribe | 21,78 | 448 | 371 | 82,81 | 198 | 0,44 | (56, 57) | (56, 57) |
| P52 | Solanales | Solanaceae | Solaneae | tribe | 20,79 | 1550 | 1000 | 64,52 | 495 | 0,32 | (56, 57) | (58) |
| P53 | Fabales | Detarioideae | *Brownea*-clade | tribe | 30,27 | 111 | 111 | 100,00 | 86 | 0,78 | (59) | (59) |
| P54 | Lamiales | Plantaginaceae | Angeloniae | tribe | 45,01 | 68 | 68 | 100,00 | 38 | 0,56 | (60) | (60) |
| P55 | Asparagales | Orchidaceae | Catasetinae | tribe | 19,5 | 262 | 262 | 100,00 | 120 | 0,46 | (51) | (30) |
| P56 | Myrtales | Myrtaceae | *Myrcia* s.l. | genus | 27,87 | 690 | 690 | 100,00 | 173 | 0,25 | (61) | (62) |
| P57 | Dioscoreales | Dioscoreaceae | Dioscoreaceae | family | 48,28 | 637 | 321 | 50,39 | 161 | 0,25 | (63) | (30, 63) |
| P58 | Malpighiales | Euphorbiaceae | *Croton* | genus | 42,54 | 1300 | 900 | 69,23 | 312 | 0,24 | (64) | (64, 65) |
| P59 | Magnoliales | Annonaceae | *Duguetia-Fusaea* | genus | 24,16 | 97 | 93 | 95,88 | 34 | 0,35 | (29) | (30) |
| P60 | Brassicales | Tropaeolaceae | *Tropaeolum* | genus | 35,74 | 88 | 88 | 100,00 | 16 | 0,18 | (29) | (30) |
| P61 | Malvales | Bombacoideae | *Eriotheca-Pachira* | genus | 8,68 | 110 | 109 | 99,09 | 14 | 0,13 | (29) | (30) |
| P62 | Sapindales | Simaroubaceae | *Simaba* | genus | 1,64 | 25 | 25 | 100,00 | 10 | 0,40 | (29) | (30) |
| P63 | Fabales | Fabaceae | *Leucaena* | genus | 0,87 | 24 | 24 | 100,00 | 10 | 0,42 | (29) | (30) |
| P64 | Caryophyllales | Cactaceae | *Pereskia* | genus | 4,56 | 17 | 17 | 100,00 | 10 | 0,59 | (29) | (30) |
| P65 | Ericales | Symplocaceae | *Symplocos* | genus | 21,5 | 300 | 105 | 35,00 | 32 | 0,11 | (29) | (66) |
| P66 | Gentianales | Apocynaceae | *Mandevilla* | genus | 0,49 | 174 | 174 | 100,00 | 48 | 0,28 | (29) | (30) |

**Table S2**. Dataset of mammal phylogenies, including the taxonomic level (Tax. level), the crown age (in million years ago), the number of species on the clade (#spp), the number of Neotropical species (#spp Neotrop) and proportion, as well as the sampling fraction (*i.e.* the number of species sampled in the tree from the total number of species described in the group).

| ID | Order | Family | Clade name | Tax. level | Crown Age | #spp | #spp Neotrop | % Neotrop | #spp sampled | Sampling fraction | Tree origin | #spp reference |
|---|---|---|---|---|---|---|---|---|---|---|---|---|
| M1 | Xenarthra | Xenarthra | Xenarthra | order | 67,95 | 32 | 32 | 100 | 32 | 1,00 | (67) | (68) |
| M2 | Chiroptera | Phyllostomidae | Phyllostomidae | family | 48,47 | 194 | 194 | 100 | 194 | 1,00 | (69) | (68) |
| M3 | Chiroptera | Molossidae | Molossini | subgenus | 22,5 | 29 | 29 | 100 | 20 | 0,69 | (70) | (68) |
| M4 | Platyrrhini | Ceboidea | Ceboidea | superfam | 21,53 | 199 | 199 | 100 | 95 | 0,48 | (71) | (72) |
| M5 | Didelphimorphia | Didelphidae | Didelphidae | Family | 25,49 | 103 | 92 | 90 | 43 | 0,42 | (73) | (68) |
| M6 | Rodentia (Myomorpha) | Cricetidae | Sigmodontinae | subfam | 12,65 | 413 | 400 | 96,85 | 279 | 0,68 | (74) | (68) |
| M7 | Rodentia (Castorimorpha) | Heteromyidae | *Heteromys* | genus | 10,6 | 16 | 16 | 100 | 11 | 0,69 | (70) | (68) |
| M8 | Rodentia (Myomorpha) | Cricetidae | Neotominae-*Reithrodontomys* | genus | 13,8 | 22 | 18 | 81,82 | 10 | 0,45 | (70) | (68) |
| M9 | Chiroptera | Emballonuroidea | Diclidurini | tribe | 34,9 | 22 | 22 | 100 | 20 | 0,91 | (70) | (75) |
| M10 | Artiodactyla | Cervidae | Odocoileini | tribe | 8,54 | 19 | 17 | 89,47 | 11 | 0,58 | (76) | (68) |
| M11 | Carnivora | Mephitidae | New World Mephitidae | family | 16 | 10 | 10 | 100 | 10 | 1,00 | (77) | (68) |
| M12 | Rodentia (Caviomorpha) | Caviomorpha | Caviomorpha | parvorder | 35,27 | 244 | 244 | 100 | 199 | 0,82 | This study | (78) |

**Table S3**. Dataset of bird phylogenies, including the taxonomic level (Tax. level), the crown age (in million years ago), the number of species on the clade (#spp), the number of Neotropical species (#spp Neotrop) and proportion, as well as the sampling fraction (*i.e.* the number of species sampled in the tree from the total number of species described in the group).

| ID | Order | Family | Clade name | Tax. level | Crown Age | #spp | #spp Neotrop | % Neotrop | #spp sampled | Sampling fraction | Tree origin | #spp reference |
|---|---|---|---|---|---|---|---|---|---|---|---|---|
| B1 | Galliformes | Cracidae | Cracidae | family | 11,82 | 55 | 55 | 100 | 39 | 0,71 | (79) | (80) |
| B2 | Apodiformes | Trochilidae | Hummingbirds | family | 26,09 | 338 | 338 | 100 | 233 | 0,69 | (79) | (81) |
| B3 | Falconiformes | Falconidae | Caracarinae+*Spiziapteryx* | Subfam | 24,78 | 11 | 11 | 100 | 8 | 0,73 | (79) | (80) |
| B4 | Psittaciformes | Psittacidae | Psittacidae | Family | 28,95 | 167 | 157 | 94,01 | 118 | 0,71 | (79) | (80) |
| B5 | Passeriformes | Grallariidae-Rhinocryptidae | Grallariidae-Rhinocryptidae | family | 25,16 | 112 | 112 | 100 | 55 | 0,49 | (79) | (80) |
| B6 | Passeriformes | Melanopareiidae-Conopophagidae | Melanopareiidae-Conopophagidae | family | 23,42 | 15 | 15 | 100 | 9 | 0,60 | (79) | (80) |
| B7 | Passeriformes | Formicariidae | Formicariidae | family | 18,54 | 11 | 11 | 100 | 7 | 0,64 | (79) | (80) |
| B8 | Passeriformes | Thamnophilidae | Thamnophilidae | family | 16,82 | 234 | 234 | 100 | 165 | 0,71 | (79) | (80) |
| B9 | Passeriformes | Funariidae | Funariidae | family | 32,58 | 301 | 301 | 100 | 284 | 0,94 | (82) | (80) |
| B10 | Passeriformes | Tyrannoidea | Tyrannoidea | superfam | 25,5 | 400 | 400 | 100 | 316 | 0,79 | (79) | (80) |
| B11 | Passeriformes | Pipridae-Cotingidae | Pipridae-Cotingidae | family | 23,51 | 118 | 118 | 100 | 80 | 0,68 | (79) | (80) |
| B12 | Passeriformes | Vireonidae | Vireonidae | family | 20,15 | 52 | 42 | 80,77 | 19 | 0,37 | (79) | (83) |
| B13 | Passeriformes | Corvidae | Neot-Corvidae | family | 17,38 | 38 | 36 | 94,74 | 36 | 0,95 | (79) | (80) |
| B14 | Passeriformes | Turdidae | *Turdus* | genus | 8,19 | 46 | 44 | 95,65 | 39 | 0,85 | (79) | (80) |
| B15 | Passeriformes | Turdidae | Neotrop-Turdidae | genus | 13,10 | 18 | 17 | 94,44 | 17 | 0,94 | (79) | (80) |
| B16 | Passeriformes | Certhioidea | Polioptilidae-Troglodytidae | family | 24,50 | 108 | 100 | 92,59 | 67 | 0,62 | (79) | (80) |

| | | | | | | | | | | | | |
|---|---|---|---|---|---|---|---|---|---|---|---|---|
| B17 | Passeriformes | Fringillidae | *Euphonia* | genus | 13,1 | 27 | 27 | 100 | 10 | 0,37 | (79) | (80) |
| B18 | Passeriformes | Passerellidae | Passerellidae | family | 11,2 | 90 | 60 | 66,67 | 89 | 0,99 | (79) | (80) |
| B19 | Passeriformes | Icteridae | Icteridae | family | 12,02 | 103 | 103 | 100 | 92 | 0,89 | (79) | (80) |
| B20 | Passeriformes | Cardinalidae | Cardinalidae | family | 18,03 | 50 | 50 | 100 | 41 | 0,82 | (79) | (80) |
| B21 | Passeriformes | Thraupidae | Thraupidae | family | 15,56 | 400 | 400 | 100 | 309 | 0,77 | (79) | (80) |
| B22 | Trogoniformes | Trogonidae | *Trogon-Priotelus* | genus | 24,66 | 23 | 23 | 100 | 20 | 0,87 | (79) | (80) |
| B23 | Piciformes | Picidae | Neotrop-*Picini* | tribe | 9,36 | 33 | 23 | 69,70 | 25 | 0,76 | (79) | (80) |
| B24 | Piciformes | Picidae | *Campephilus* (Neotrop-*Megapicini*) | genus | 6,01 | 11 | 11 | 100 | 10 | 0,91 | (79) | (80) |
| B25 | Piciformes | Picidae | *Melanerpes* | genus | 10,53 | 24 | 20 | 83,33 | 12 | 0,50 | (79) | (80) |
| B26 | Piciformes | Picidae | *Veniliornis* | genus | 5,38 | 14 | 14 | 100 | 12 | 0,86 | (79) | (80) |
| B27 | Piciformes | Picidae | *Picumnus* | genus | 12,17 | 28 | 28 | 100 | 8 | 0,29 | (79) | (80) |
| B28 | Piciformes | Ramphastides | Capitonidae, Semnornithidae,Ramphastidae | infraord | 20,11 | 59 | 59 | 100 | 44 | 0,75 | (79) | (80) |
| B29 | Galbuliformes | Bucconidae | Neotrop-Bucconidae | family | 42,09 | 37 | 37 | 100 | 10 | 0,27 | (79) | (80) |
| B30 | Accipitriformes | Accipitridae | *Buteogallus* | genus | 9,64 | 9 | 9 | 100 | 9 | 1,00 | (79) | (80) |
| B31 | Columbiformes | Columbidae | *Patagioenas* | genus | 11,05 | 17 | 17 | 100 | 12 | 0,71 | (79) | (80) |
| B32 | Columbiformes | Columbidae | Neotrop-Columbinae | tribe | 20,68 | 17 | 17 | 100 | 13 | 0,76 | (79) | (80) |

**Table S4**. Dataset of Squamata phylogenies, including the taxonomic level (Tax. level), the crown age (in million years ago), the number of species on the clade (#spp), the number of Neotropical species (#spp Neotrop) and proportion, as well as the sampling fraction (*i.e.* the number of species sampled in the tree from the total number of species described in the group).

| ID | Order | Clade name | Tax. level | Crown Age | #spp | #spp Neotrop | % Neotrop | #spp sampled | Sampling fraction | Tree origin | #spp reference |
|---|---|---|---|---|---|---|---|---|---|---|---|
| S1 | Squamata | Colubridae (sf: Dipsadidae) | tribe | 34,35 | 154 | 154 | 100,00 | 76 | 0,49 | (84) | (85) |
| S2 | Squamata | Colubridae (sf: Colubrinae) | tribe | 26,03 | 39 | 28 | 71,79 | 15 | 0,38 | (84) | (85) |
| S3 | Squamata | Elapidae | family | 21,78 | 80 | 80 | 100,00 | 19 | 0,24 | (84) | (85) |
| S4 | Squamata | Viperidae (sf. Crotalinae) | tribe | 21,14 | 21 | 21 | 100,00 | 13 | 0,62 | (84) | (85) |
| S5 | Squamata | Viperidae (sf. Crotalinae) | tribe | 21,41 | 71 | 71 | 100,00 | 43 | 0,61 | (84) | (85) |
| S6 | Squamata | Boidae | family | 44,84 | 32 | 32 | 100,00 | 15 | 0,47 | (84) | (85) |
| S7 | Squamata | Aniliidae, Tropidophiidae | family | 80,54 | 35 | 35 | 100,00 | 9 | 0,26 | (84) | (85) |
| S8 | Squamata | Typhlopidae | family | 43,74 | 59 | 59 | 100,00 | 29 | 0,49 | (84) | (85) |
| S9 | Squamata | Corytophanidae, Dactyloidae | family | 83,4 | 436 | 436 | 100,00 | 214 | 0,49 | (84) | (85) |
| S10 | Squamata | Liolaemidae | family | 71,14 | 308 | 308 | 100,00 | 119 | 0,39 | (84) | (85) |
| S11 | Squamata | Leiosauridae | family | 42,19 | 33 | 33 | 100,00 | 11 | 0,33 | (84) | (85) |
| S12 | Squamata | Polychrotidae, Hoplocercidae | family | 81,21 | 27 | 27 | 100,00 | 12 | 0,44 | (84) | (85) |
| S13 | Squamata | Phrynosomatidae | family | 19,79 | 102 | 50 | 49,02 | 44 | 0,43 | (84) | (85) |
| S14 | Squamata | Iguanidae | family | 34,56 | 29 | 29 | 100,00 | 22 | 0,76 | (84) | (85) |
| S15 | Squamata | Tropiduridae | family | 88,47 | 136 | 136 | 100,00 | 78 | 0,57 | (84) | (85) |
| S16 | Squamata | Anguidae (Gerrhonotinae) | tribe | 26,47 | 43 | 43 | 100,00 | 18 | 0,42 | (84) | (85) |
| S17 | Squamata | Amphisbaenidae | family | 48,84 | 95 | 95 | 100,00 | 37 | 0,39 | (84) | (85) |
| S18 | Squamata | Teiidae, Alopoglossidae, Gymnophthalmidae | family | 86,27 | 337 | 292 | 86,65 | 144 | 0,43 | (84) | (85) |
| S19 | Squamata | Scincidae (Mabuyinae) | tribe | 25,22 | 60 | 60 | 100,00 | 20 | 0,33 | (84) | (85) |
| S20 | Squamata | Xantusiidae | family | 36,33 | 19 | 19 | 100,00 | 16 | 0,84 | (84) | (85) |

| | | | | | | | | | | | |
|---|---|---|---|---|---|---|---|---|---|---|---|
| S21 | Squamata | Phyllodactylidae | family | 66,91 | 65 | 65 | 100,00 | 20 | 0,31 | (84) | (85) |
| S22 | Squamata | Sphaerodactylidae | family | 70,88 | 170 | 170 | 100,00 | 69 | 0,41 | (84) | (85) |
| S23 | Squamata | Colubridae (sf: Dipsadidae) | tribe | 34,42 | 123 | 123 | 100,00 | 60 | 0,49 | (84) | (85) |
| S24 | Squamata | Colubridae (sf: Dipsadidae) | tribe | 29,25 | 313 | 313 | 100,00 | 45 | 0,14 | (84) | (85) |

**Table S5**. Dataset of Amphibia phylogenies, including the taxonomic level (Tax. level), the crown age (in million years ago), the number of species on the clade (#spp), the number of Neotropical species (#spp Neotrop) and proportion, as well as the sampling fraction (*i.e.* the number of species sampled in the tree from the total number of species described in the group).

| ID | Order | Clade name | Tax. level | Crown Age | #spp | #spp Neotrop | % Neotrop | #spp sampled | Sampling fraction | Tree origin | #spp reference |
|---|---|---|---|---|---|---|---|---|---|---|---|
| A1 | Anura | Aromobatidae | family | 67,18 | 127 | 127 | 100 | 118 | 0,93 | (86) | (87) |
| A2 | Anura | Dendrobatidae | family | 61,2 | 136 | 136 | 100 | 136 | 1,00 | (86) | (87) |
| A3 | Anura | Hemiphractidae | family | 80,69 | 109 | 109 | 100 | 86 | 0,79 | (86) | (87) |
| A4 | Anura | Eleutherodactylidae | family | 72,08 | 217 | 212 | 97,7 | 170 | 0,78 | (86) | (87) |
| A5 | Anura | Craugastoridae (sf: Craugastorinae, excluding Haddadus) | family | 69,87 | 131 | 126 | 96,2 | 61 | 0,47 | (86) | (88) |
| A6 | Anura | Craugastoridae (Ceuthomantinae and (part of) Holoadeninae (other is paraphyletic)) | family | 68,16 | 625 | 625 | 100 | 295 | 0,47 | (86) | (88) |
| A7 | Anura | Hylidae (Phyllomedusinae) | tribe | 54,78 | 63 | 63 | 100 | 50 | 0,79 | (86) | (89) |
| A8 | Anura | Hylidae (Hylinae) | tribe | 69,93 | 183 | 183 | 100 | 117 | 0,64 | (86) | (87) |
| A9 | Anura | Hylidae (Hylinae) | tribe | 72,8 | 252 | 252 | 100 | 147 | 0,58 | (86) | (87) |
| A10 | Anura | Hylidae (Lophyohylinae) | tribe | 43,23 | 85 | 85 | 100 | 60 | 0,71 | (86) | (87) |
| A11 | Anura | Odontophrynidae, Ceratophryidae, Rhinodermatidae, Telmatobiidae, Cycloramphidae, Hylodidae, Batrichylidae, Alsodidae | family | 87,37 | 259 | 259 | 100 | 127 | 0,49 | (86) | (87) |
| A12 | Anura | Leptodactylidae | family | 78,07 | 211 | 211 | 100 | 186 | 0,88 | (86) | (87) |
| A13 | Anura | Centrolenidae | family | 33,39 | 158 | 158 | 100 | 128 | 0,81 | (86) | (87) |
| A14 | Anura | Bufonidae | family | 53,95 | 119 | 119 | 100 | 54 | 0,45 | (86) | (87) |
| A15 | Anura | Bufonidae | family | 33,89 | 157 | 132 | 84 | 110 | 0,70 | (86) | (88, 89) |
| A16 | Caudata | Plethodontidae (sf: Hemidactyliinae) | family | 73,05 | 307 | 307 | 100 | 159 | 0,52 | (90) | (87) |

**Table S6.** Summary results showing the best-fit model (among constant and time-dependent models) for each clade and the derived species richness pattern (gradual increase [Sc. 1], exponential increase [Sc. 2], saturated increase [Sc. 3] and decline [Sc. 4]). For each model category, we fitted three models in which speciation (Birth "B") and/or extinction (Death "D") remain constant ("cst"), or vary ("Var.") continuously with time. The value of speciation (lambda) and extinction (mu) rates is provided, as well as the value of the dependence between speciation and extinction rates with time (alpha and beta values, respectively). For time-dependent models, $\alpha$ and $\beta > 0$ reflect decreasing speciation and extinction towards the present, respectively, while $\alpha$ and $\beta < 0$ indicate the opposite, increasing speciation and extinction towards the present. Species richness patterns are derived from the diversification trend through time. For each phylogeny, the sampling fraction (*i.e.* number of species sampled from the total described), crown age (in million years ago), and number of species (#spp) is provided together with the main geographic (Andean-centered, Amazonian-centered, or other), altitudinal (lowland, montane, highland) and habitat distribution (tropical open vegetation, tropical forest vegetation or non-tropical) of the species in the clade. Mixed patterns could be observed in all categories. Abbreviations for model type; B = Birth; D = Death; cst = constant; Var. = variable.

| Taxa | Clade | # spp. | Sampling fraction | Crown age | Distribution | Habitat | Altitude | Model type | Lambda | Alpha | Mu | Beta | Richness pattern |
|---|---|---|---|---|---|---|---|---|---|---|---|---|---|
| Plants | P1 | 13 | 0.12 | 2.88 | Mixed | Tropical forest | Lowland-Montane | BcstDTimeVar | 2.08 | NA | 0.02 | 2.52 | Exponential |
| Plants | P2 | 38 | 0.28 | 0.94 | Andean | Tropical forest | Lowland-Montane | BcstDTimeVar | 6.43 | NA | 0.15 | 5.64 | Exponential |
| Plants | P3 | 12 | 0.16 | 43.02 | Andean | Tropical forest | Lowland-Montane | BcstDcst | 0.15 | NA | 0.09 | NA | Gradual |
| Plants | P4 | 10 | 0.12 | 13.8 | Amazonian | Tropical forest | Lowland-Montane | BcstDcst | 1.02 | NA | 0.89 | NA | Gradual |
| Plants | P5 | 10 | 0.12 | 7.32 | Amazonian | Tropical forest | Lowland-Montane | BcstDcst | 1.35 | NA | 1.04 | NA | Gradual |
| Plants | P6 | 14 | 0.45 | 5.82 | Andean | Open tropical | Lowland-Montane | BcstDcst | 0.47 | NA | 0 | NA | Gradual |
| Plants | P7 | 29 | 0.57 | 12.27 | Amazonian | Tropical forest | Lowland-Montane | BcstDcst | 0.29 | NA | 0 | NA | Gradual |
| Plants | P8 | 15 | 0.14 | 11.88 | Andean | Tropical forest | Lowland-Montane | BTimeVarDcst | 1.12 | -0.27 | 0 | NA | Exponential |
| Plants | P9 | 10 | 0.15 | 19.63 | Amazonian | Tropical forest | Lowland-Montane | BcstDcst | 0.18 | NA | 0 | NA | Gradual |

| | | | | | | | | | | | | | |
|---|---|---|---|---|---|---|---|---|---|---|---|---|---|
| Plants | P10 | 24 | 0.41 | 25.52 | Amazonian | Tropical forest | Lowland-Montane | BcstDcst | 0.14 | NA | 0 | NA | Gradual |
| Plants | P11 | 42 | 0.68 | 15.15 | Mixed | Tropical forest | Lowland-Montane | BcstDcst | 0.43 | NA | 0.24 | NA | Gradual |
| Plants | P12 | 15 | 0.35 | 13.04 | Mixed | Tropical forest | Lowland-Montane | BcstDcst | 0.3 | NA | 0.08 | NA | Gradual |
| Plants | P13 | 46 | 0.32 | 56.31 | Mixed | Tropical forest | Lowland-Montane | BTimeVarDcst | 0.24 | 0.01 | 0.29 | NA | Declining |
| Plants | P14 | 10 | 0.2 | 26 | Andean | Tropical forest | Lowland-Montane | BcstDcst | 0.43 | NA | 0.38 | NA | Gradual |
| Plants | P15 | 19 | 0.63 | 19.88 | Amazonian | Tropical forest | Lowland-Montane | BcstDcst | 0.19 | NA | 0.05 | NA | Gradual |
| Plants | P16 | 19 | 0.21 | 7.08 | Mixed | Tropical forest | Lowland-Montane | BcstDcst | 1.14 | NA | 0.69 | NA | Gradual |
| Plants | P17 | 18 | 0.18 | 22.42 | Mixed | Tropical forest | Lowland-Montane | BcstDcst | 0.27 | NA | 0.14 | NA | Gradual |
| Plants | P18 | 7 | 0.88 | 10.8 | Andean | Tropical forest | Lowland-Montane | BcstDcst | 0.11 | NA | 0 | NA | Gradual |
| Plants | P19 | 22 | 0.18 | 28.02 | Andean | Tropical forest | Lowland-Montane | BcstDTimeVar | 0.21 | NA | 0 | 0.39 | Exponential |
| Plants | P20 | 29 | 0.72 | 20.25 | Andean | Tropical forest | Lowland-Montane | BcstDcst | 0.14 | NA | 0 | NA | Gradual |
| Plants | P21 | 103 | 0.41 | 20.77 | Amazonian | Tropical forest | Lowland-Montane | BTimeVarDcst | 0.09 | 0.11 | 0 | NA | Saturated |
| Plants | P22 | 19 | 0.43 | 17.64 | Andean | Tropical forest | Montane-Highland | BcstDcst | 0.17 | NA | 0 | NA | Gradual |
| Plants | P23 | 9 | 0.56 | 25.05 | Amazonian | Tropical forest | Lowland-Montane | BcstDcst | 0.13 | NA | 0.07 | NA | Gradual |
| Plants | P24 | 200 | 0.36 | 17.03 | Andean | Mixed | Mixed | BTimeVarDcst | 1.64 | -0.29 | 0 | NA | Exponential |
| Plants | P25 | 24 | 0.89 | 19.21 | Other | Tropical forest | Lowland-Montane | BcstDcst | 0.14 | NA | 0 | NA | Gradual |
| Plants | P26 | 43 | 0.61 | 14.63 | Other | Tropical forest | Lowland-Montane | BcstDcst | 0.23 | NA | 0 | NA | Gradual |
| Plants | P27 | 19 | 0.54 | 4.67 | Other | Temperate | Lowland-Montane | BcstDcst | 0.74 | NA | 0.17 | NA | Gradual |
| Plants | P28 | 37 | 0.28 | 2.75 | Andean | Open tropical | Montane-Highland | BcstDTimeVar | 2.18 | NA | 0 | 4.34 | Exponential |
| Plants | P29 | 13 | 0.93 | 15.29 | Amazonian | Tropical forest | Lowland-Montane | BcstDcst | 0.11 | NA | 0 | NA | Gradual |
| Plants | P30 | 14 | 0.88 | 37.32 | Other | Tropical forest | Lowland-Montane | BcstDcst | 0.1 | NA | 0.06 | NA | Gradual |
| Plants | P31 | 78 | 0.98 | 30.97 | Other | Tropical forest | Lowland-Montane | BcstDcst | 0.19 | NA | 0.08 | NA | Gradual |
| Plants | P32 | 22 | 0.79 | 20.51 | Other | Tropical forest | Lowland-Montane | BcstDcst | 0.14 | NA | 0 | NA | Gradual |
| Plants | P33 | 16 | 1 | 19.96 | Other | Mixed | Lowland-Montane | BcstDcst | 0.09 | NA | 0 | NA | Gradual |
| Plants | P34 | 32 | 1 | 29.68 | Andean | Tropical forest | Lowland-Montane | BcstDcst | 0.21 | NA | 0.14 | NA | Gradual |

| | | | | | | | | | | | | | |
|---|---|---|---|---|---|---|---|---|---|---|---|---|---|
| Plants | P35 | 114 | 0.99 | 38.07 | Andean | Tropical forest | Lowland-Montane | BcstDcst | 0.13 | NA | 0 | NA | Gradual |
| Plants | P36 | 154 | 0.99 | 27.09 | Mixed | Mixed | Lowland-Montane | BcstDcst | 0.2 | NA | 0 | NA | Gradual |
| Plants | P37 | 178 | 0.99 | 43.62 | Mixed | Tropical forest | Lowland-Montane | BcstDTimeVar | 0.19 | NA | 0 | 0.14 | Exponential |
| Plants | P38 | 102 | 1 | 26.92 | Andean | Tropical forest | Lowland-Montane | BcstDTimeVar | 0.25 | NA | 0.01 | 0.18 | Exponential |
| Plants | P39 | 22 | 1 | 34.21 | Mixed | Tropical forest | Lowland-Montane | BcstDcst | 0.26 | NA | 0.26 | NA | Gradual |
| Plants | P40 | 271 | 0.46 | 20.39 | Mixed | Mixed | Lowland-Montane | BTimeVarDcst | 0.76 | -0.05 | 0.25 | NA | Exponential |
| Plants | P41 | 670 | 0.13 | 17.87 | Andean | Tropical forest | Montane-Highland | BcstDcst | 3.08 | NA | 2.76 | NA | Gradual |
| Plants | P42 | 789 | 0.21 | 31.48 | Andean | Tropical forest | Mixed | BTimeVarDcst | 0.75 | -0.02 | 0.41 | NA | Exponential |
| Plants | P43 | 168 | 0.25 | 21.01 | Mixed | Open tropical | Lowland-Montane | BcstDTimeVar | 0.71 | NA | 0.31 | 0.05 | Exponential |
| Plants | P44 | 126 | 0.33 | 6 | Mixed | Tropical forest | Lowland-Montane | BTimeVarDcst | 3.76 | -0.68 | 0 | NA | Exponential |
| Plants | P45 | 588 | 0.49 | 48.45 | Andean | Tropical forest | Lowland-Montane | BTimeVarDTimeVar | 0.16 | 0.01 | 0.01 | 0.09 | Exponential |
| Plants | P46 | 111 | 0.79 | 28.92 | Amazonian | Tropical forest | Lowland-Montane | BcstDcst | 0.16 | NA | 0 | NA | Gradual |
| Plants | P47 | 21 | 1 | 28.86 | Other | Temperate | Lowland-Montane | BcstDcst | 0.34 | NA | 0.34 | NA | Gradual |
| Plants | P48 | 50 | 0.23 | 32.21 | Andean | Tropical forest | Lowland-Montane | BcstDcst | 3.13 | NA | 3.13 | NA | Gradual |
| Plants | P49 | 44 | 0.3 | 30.99 | Other | Temperate | Montane-Highland | BcstDcst | 1.18 | NA | 1.14 | NA | Gradual |
| Plants | P50 | 63 | 0.71 | 7.21 | Other | Temperate | Lowland-Montane | BcstDcst | 0.59 | NA | 0 | NA | Gradual |
| Plants | P51 | 189 | 0.44 | 21.79 | Mixed | Mixed | Lowland-Montane | BcstDcst | 0.59 | NA | 0.38 | NA | Gradual |
| Plants | P52 | 495 | 0.32 | 20.79 | Andean | Tropical forest | Mixed | BTimeVarDcst | 0.59 | -0.07 | 0 | NA | Exponential |
| Plants | P53 | 86 | 0.77 | 30.28 | Amazonian | Tropical forest | Lowland-Montane | BcstDcst | 0.15 | NA | 0 | NA | Gradual |
| Plants | P54 | 38 | 0.56 | 45.01 | Other | Mixed | Lowland-Montane | BcstDcst | 0.19 | NA | 0.13 | NA | Gradual |
| Plants | P55 | 120 | 0.46 | 19.5 | Amazonian | Tropical forest | Lowland-Montane | BTimeVarDcst | 1.01 | -0.21 | 0 | NA | Exponential |
| Plants | P56 | 150 | 0.25 | 27.88 | Mixed | Mixed | Lowland-Montane | BcstDcst | 0.25 | NA | 0.05 | NA | Gradual |
| Plants | P57 | 161 | 0.25 | 48.28 | Mixed | Mixed | Lowland-Montane | BcstDcst | 0.18 | NA | 0.07 | NA | Gradual |
| Plants | P58 | 307 | 0.24 | 42.38 | Mixed | Mixed | Lowland-Montane | BcstDcst | 0.53 | NA | 0.4 | NA | Gradual |
| Plants | P59 | 34 | 0.35 | 24.17 | Amazonian | Tropical forest | Lowland-Montane | BcstDTimeVar | 0.43 | NA | 0.08 | 0.15 | Exponential |

| | | | | | | | | | | | | | |
|---|---|---|---|---|---|---|---|---|---|---|---|---|---|
| Plants | P60 | 16 | 0.18 | 35.74 | Andean | Open tropical | Lowland-Montane | BcstDTimeVar | 0.17 | NA | 0 | 0.25 | Exponential |
| Plants | P61 | 14 | 0.13 | 8.68 | Amazonian | Tropical forest | Lowland-Montane | BcstDcst | 0.51 | NA | 0.07 | NA | Gradual |
| Plants | P62 | 10 | 0.4 | 1.65 | Mixed | Tropical forest | Lowland-Montane | BcstDcst | 1.7 | NA | 0 | NA | Gradual |
| Plants | P63 | 10 | 0.42 | 0.88 | Other | Open tropical | Lowland-Montane | BcstDcst | 2.23 | NA | 0 | NA | Gradual |
| Plants | P64 | 10 | 0.59 | 4.57 | Other | Mixed | Lowland-Montane | BcstDcst | 0.42 | NA | 0 | NA | Gradual |
| Plants | P65 | 32 | 0.11 | 21.51 | Andean | Tropical forest | Montane-Highland | BcstDcst | 0.71 | NA | 0.55 | NA | Gradual |
| Plants | P66 | 48 | 0.28 | 0.5 | Mixed | Mixed | Lowland-Montane | BcstDcst | 8.82 | NA | 0 | NA | Gradual |
| Mammals | M1 | 32 | 1 | 67.96 | Amazonian | Mixed | Lowland-Montane | BcstDcst | 0.08 | NA | 0.05 | NA | Gradual |
| Mammals | M2 | 192 | 0.86 | 41.86 | Amazonian | Mixed | Lowland-Montane | BcstDcst | 0.12 | NA | 0 | NA | Gradual |
| Mammals | M3 | 20 | 0.69 | 22.5 | Mixed | Tropical forest | Lowland-Montane | BTimeVarDTimeVar | 0.01 | 0.36 | 0.03 | 0.33 | Declining |
| Mammals | M4 | 95 | 0.48 | 21.54 | Amazonian | Tropical forest | Lowland-Montane | BcstDcst | 1.07 | NA | 0.97 | NA | Gradual |
| Mammals | M5 | 43 | 0.42 | 25.49 | Amazonian | Tropical forest | Lowland-Montane | BcstDcst | 0.25 | NA | 0.12 | NA | Gradual |
| Mammals | M6 | 279 | 0.68 | 12.66 | Andean | Mixed | Mixed | BTimeVarDcst | 0.39 | 0.05 | 0 | NA | Saturated |
| Mammals | M7 | 11 | 0.69 | 10.6 | Other | Mixed | Lowland-Montane | BcstDcst | 0.17 | NA | 0 | NA | Gradual |
| Mammals | M8 | 10 | 0.46 | 13.81 | Andean | Mixed | Lowland-Montane | BcstDcst | 0.45 | NA | 0.41 | NA | Gradual |
| Mammals | M9 | 20 | 0.91 | 34.9 | Amazonian | Tropical forest | Lowland-Montane | BTimeVarDcst | 0.02 | 0.06 | 0 | NA | Saturated |
| Mammals | M10 | 11 | 0.58 | 8.54 | Andean | Mixed | Mixed | BcstDcst | 0.23 | NA | 0 | NA | Gradual |
| Mammals | M11 | 10 | 1 | 16 | Amazonian | Mixed | Lowland-Montane | BcstDcst | 0.1 | NA | 0 | NA | Gradual |
| Mammals | M12 | 199 | 0.82 | 35.28 | Amazonian | Mixed | Lowland-Montane | BTimeVarDcst | 0.24 | -0.04 | 0 | NA | Exponential |
| Birds | B1 | 39 | 0.71 | 11.82 | Amazonian | Tropical forest | Lowland-Montane | BcstDcst | 0.33 | NA | 0 | NA | Gradual |
| Birds | B2 | 233 | 0.69 | 26.1 | Andean | Mixed | Mixed | BcstDcst | 0.18 | NA | 0 | NA | Gradual |
| Birds | B3 | 8 | 0.73 | 24.79 | Mixed | Mixed | Lowland-Montane | BcstDcst | 0.3 | NA | 0.36 | NA | Gradual |
| Birds | B4 | 118 | 0.71 | 28.95 | Amazonian | Tropical forest | Mixed | BcstDcst | 0.18 | NA | 0.02 | NA | Gradual |
| Birds | B5 | 55 | 0.49 | 25.17 | Andean | Tropical forest | Lowland-Montane | BcstDcst | 0.15 | NA | 0 | NA | Gradual |
| Birds | B6 | 9 | 0.6 | 23.42 | Andean | Mixed | Lowland-Montane | BcstDcst | 0.09 | NA | 0 | NA | Gradual |

| | | | | | | | | | | | | | |
|---|---|---|---|---|---|---|---|---|---|---|---|---|---|
| Birds | B7 | 7 | 0.64 | 18.55 | Andean | Tropical forest | Lowland-Montane | BcstDcst | 0.2 | NA | 0.2 | NA | Gradual |
| Birds | B8 | 165 | 0.71 | 16.82 | Amazonian | Tropical forest | Lowland-Montane | BTimeVarDcst | 0.12 | 0.11 | 0 | NA | Saturated |
| Birds | B9 | 292 | 0.94 | 32.59 | Mixed | Mixed | Mixed | BcstDcst | 0.15 | NA | 0 | NA | Gradual |
| Birds | B10 | 316 | 0.79 | 25.5 | Mixed | Mixed | Mixed | BTimeVarDcst | 0.15 | 0.02 | 0 | NA | Saturated |
| Birds | B11 | 80 | 0.68 | 23.52 | Amazonian | Tropical forest | Lowland-Montane | BcstDcst | 0.18 | NA | 0 | NA | Gradual |
| Birds | B12 | 19 | 0.37 | 20.15 | Other | Tropical forest | Lowland-Montane | BcstDcst | 0.2 | NA | 0.04 | NA | Gradual |
| Birds | B13 | 36 | 0.95 | 17.38 | Andean | Mixed | Lowland-Montane | BTimeVarDcst | 0.1 | 0.06 | 0 | NA | Saturated |
| Birds | B14 | 39 | 0.85 | 8.2 | Andean | Mixed | Mixed | BTimeVarDcst | 0.06 | 0.37 | 0 | NA | Saturated |
| Birds | B15 | 17 | 0.94 | 13.11 | Amazonian | Tropical forest | Lowland-Montane | BcstDcst | 0.14 | NA | 0 | NA | Gradual |
| Birds | B16 | 67 | 0.62 | 24.51 | Amazonian | Tropical forest | Lowland-Montane | BTimeVarDcst | 0.12 | 0.05 | 0 | NA | Saturated |
| Birds | B17 | 10 | 0.37 | 13.1 | Mixed | Mixed | Lowland-Montane | BcstDcst | 0.19 | NA | 0 | NA | Gradual |
| Birds | B18 | 89 | 0.99 | 11.21 | Mixed | Mixed | Lowland-Montane | BTimeVarDcst | 0.11 | 0.19 | 0 | NA | Saturated |
| Birds | B19 | 92 | 0.89 | 12.02 | Andean | Mixed | Lowland-Montane | BTimeVarDcst | 0.18 | 0.11 | 0 | NA | Saturated |
| Birds | B20 | 41 | 0.82 | 18.04 | Andean | Mixed | Lowland-Montane | BTimeVarDcst | 0.05 | 0.13 | 0 | NA | Saturated |
| Birds | B21 | 309 | 0.77 | 13.57 | Other | Open tropical | Lowland-Montane | BTimeVarDcst | 0.18 | 0.12 | 0.08 | NA | Saturated |
| Birds | B22 | 20 | 0.87 | 24.67 | Mixed | Mixed | Mixed | BcstDcst | 0.1 | NA | 0 | NA | Gradual |
| Birds | B23 | 25 | 0.76 | 9.37 | Amazonian | Tropical forest | Lowland-Montane | BTimeVarDcst | 0.13 | 0.18 | 0 | NA | Saturated |
| Birds | B24 | 10 | 0.91 | 6.01 | Amazonian | Tropical forest | Lowland-Montane | BcstDcst | 0.26 | NA | 0 | NA | Gradual |
| Birds | B25 | 12 | 0.5 | 10.54 | Mixed | Tropical forest | Lowland-Montane | BcstDcst | 0.21 | NA | 0 | NA | Gradual |
| Birds | B26 | 12 | 0.86 | 5.38 | Amazonian | Mixed | Lowland-Montane | BcstDcst | 0.32 | NA | 0 | NA | Gradual |
| Birds | B27 | 8 | 0.29 | 12.17 | Mixed | Tropical forest | Lowland-Montane | BcstDcst | 0.62 | NA | 0.56 | NA | Gradual |
| Birds | B28 | 44 | 0.75 | 20.12 | Amazonian | Mixed | Lowland-Montane | BcstDcst | 0.22 | NA | 0.04 | NA | Gradual |
| Birds | B29 | 10 | 0.27 | 42.09 | Amazonian | Tropical forest | Lowland-Montane | BcstDcst | 0.12 | NA | 0.08 | NA | Gradual |
| Birds | B30 | 9 | 1 | 9.65 | Amazonian | Mixed | Lowland-Montane | BcstDcst | 0.15 | NA | 0 | NA | Gradual |
| Birds | B31 | 12 | 0.71 | 11.06 | Andean | Mixed | Mixed | BcstDcst | 0.2 | NA | 0 | NA | Gradual |

| | | | | | | | | | | | | | |
|---|---|---|---|---|---|---|---|---|---|---|---|---|---|
| Birds | B32 | 13 | 0.76 | 20.68 | Andean | Open tropical | Lowland-Montane | BcstDcst | 0.1 | NA | 0 | NA | Gradual |
| Squamata | S1 | 76 | 0.49 | 34.35 | Amazonian | Mixed | Lowland-Montane | BTimeVarDcst | 0.06 | 0.04 | 0 | NA | Saturated |
| Squamata | S2 | 15 | 0.38 | 26.03 | Andean | Mixed | Lowland-Montane | BTimeVarDcst | 0.04 | 0.07 | 0 | NA | Saturated |
| Squamata | S3 | 19 | 0.24 | 21.78 | Andean | Mixed | Lowland-Montane | BcstDcst | 0.28 | NA | 0.13 | NA | Gradual |
| Squamata | S4 | 13 | 0.62 | 21.14 | Andean | Tropical forest | Lowland-Montane | BcstDcst | 0.1 | NA | 0 | NA | Gradual |
| Squamata | S5 | 43 | 0.61 | 21.41 | Andean | Mixed | Lowland-Montane | BcstDcst | 0.16 | NA | 0 | NA | Gradual |
| Squamata | S6 | 15 | 0.47 | 44.84 | Mixed | Mixed | Lowland-Montane | BcstDcst | 0.06 | NA | 0 | NA | Gradual |
| Squamata | S7 | 9 | 0.26 | 80.54 | Amazonian | Tropical forest | Lowland-Montane | BcstDcst | 0.23 | NA | 0.24 | NA | Gradual |
| Squamata | S8 | 29 | 0.49 | 43.74 | Amazonian | Tropical forest | Lowland-Montane | BTimeVarDTimeVar | 0.58 | -0.06 | 1.65 | -0.29 | Declining |
| Squamata | S9 | 214 | 0.49 | 83.4 | Other | Tropical forest | Lowland-Montane | BTimeVarDcst | 0.06 | 0.01 | 0 | NA | Saturated |
| Squamata | S10 | 119 | 0.39 | 71.14 | Andean | Temperate | Montane-Highland | BTimeVarDcst | 0.23 | -0.04 | 0 | NA | Exponential |
| Squamata | S11 | 11 | 0.33 | 42.19 | Other | Mixed | Lowland-Montane | BcstDcst | 0.06 | NA | 0 | NA | Gradual |
| Squamata | S12 | 12 | 0.44 | 81.21 | Amazonian | Tropical forest | Lowland-Montane | BcstDcst | 0.03 | NA | 0 | NA | Gradual |
| Squamata | S13 | 44 | 0.43 | 19.79 | Other | Mixed | Mixed | BTimeVarDcst | 0.12 | 0.05 | 0 | NA | Saturated |
| Squamata | S14 | 22 | 0.76 | 34.56 | Andean | Tropical forest | Lowland-Montane | BcstDcst | 0.12 | NA | 0.06 | NA | Gradual |
| Squamata | S15 | 78 | 0.57 | 88.47 | Andean | Mixed | Lowland-Montane | BcstDcst | 0.05 | NA | 0 | NA | Gradual |
| Squamata | S16 | 18 | 0.42 | 26.47 | Other | Tropical forest | Lowland-Montane | BcstDcst | 0.11 | NA | 0.01 | NA | Gradual |
| Squamata | S17 | 37 | 0.39 | 48.84 | Amazonian | Mixed | Lowland-Montane | BTimeVarDcst | 0.02 | 0.04 | 0 | NA | Saturated |
| Squamata | S18 | 144 | 0.43 | 86.27 | Mixed | Mixed | Mixed | BcstDcst | 0.06 | NA | 0 | NA | Gradual |
| Squamata | S19 | 20 | 0.33 | 25.22 | Andean | Mixed | Lowland-Montane | BcstDcst | 0.12 | NA | 0 | NA | Gradual |
| Squamata | S20 | 16 | 0.84 | 36.33 | Andean | Tropical forest | Lowland-Montane | BTimeVarDcst | 0.02 | 0.06 | 0 | NA | Saturated |
| Squamata | S21 | 20 | 0.31 | 66.91 | Andean | Mixed | Lowland-Montane | BcstDcst | 0.06 | NA | 0 | NA | Gradual |
| Squamata | S22 | 69 | 0.41 | 70.89 | Amazonian | Tropical forest | Lowland-Montane | BTimeVarDTimeVar | 0.1 | 0 | 4.01 | -2.25 | Declining |
| Squamata | S23 | 60 | 0.49 | 34.42 | Other | Mixed | Mixed | BcstDcst | 0.11 | NA | 0 | NA | Gradual |
| Squamata | S24 | 45 | 0.14 | 29.25 | Andean | Tropical forest | Lowland-Montane | BTimeVarDcst | 0.11 | 0.03 | 0 | NA | Saturated |

| | | | | | | | | | | | | | |
|---|---|---|---|---|---|---|---|---|---|---|---|---|---|
| Amphibia | A1 | 118 | 0.93 | 67.18 | Mixed | Tropical forest | Lowland-Montane | BTimeVarDcst | 0.05 | 0.01 | 0 | NA | Saturated |
| Amphibia | A2 | 136 | 1 | 67.35 | Andean | Tropical forest | Mixed | BcstDcst | 0.06 | NA | 0 | NA | Gradual |
| Amphibia | A3 | 86 | 0.79 | 80.69 | Andean | Tropical forest | Lowland-Montane | BcstDcst | 0.04 | NA | 0 | NA | Gradual |
| Amphibia | A4 | 170 | 0.78 | 72.08 | Other | Tropical forest | Mixed | BTimeVarDTimeVar | 0.04 | 0.05 | 0.01 | 0.08 | Exponential |
| Amphibia | A5 | 61 | 0.47 | 69.87 | Other | Tropical forest | Lowland-Montane | BTimeVarDcst | 0.04 | 0.01 | 0 | NA | Saturated |
| Amphibia | A6 | 295 | 0.47 | 68.16 | Andean | Tropical forest | Mixed | BTimeVarDcst | 0.07 | 0.01 | 0 | NA | Saturated |
| Amphibia | A7 | 50 | 0.79 | 54.78 | Amazonian | Tropical forest | Lowland-Montane | BcstDcst | 0.06 | NA | 0 | NA | Gradual |
| Amphibia | A8 | 117 | 0.64 | 69.93 | Amazonian | Tropical forest | Lowland-Montane | BcstDcst | 0.09 | NA | 0.03 | NA | Gradual |
| Amphibia | A9 | 147 | 0.58 | 72.8 | Amazonian | Tropical forest | Lowland-Montane | BcstDcst | 0.07 | NA | 0 | NA | Gradual |
| Amphibia | A10 | 60 | 0.71 | 43.33 | Amazonian | Tropical forest | Lowland-Montane | BcstDcst | 0.08 | NA | 0 | NA | Gradual |
| Amphibia | A11 | 127 | 0.49 | 87.37 | Mixed | Mixed | Lowland-Montane | BTimeVarDTimeVar | 0.28 | -0.02 | 0.37 | -0.05 | Declining |
| Amphibia | A12 | 186 | 0.88 | 78.07 | Mixed | Mixed | Lowland-Montane | BcstDcst | 0.06 | NA | 0 | NA | Gradual |
| Amphibia | A13 | 128 | 0.81 | 33.4 | Andean | Tropical forest | Lowland-Montane | BTimeVarDcst | 0.07 | 0.05 | 0 | NA | Saturated |
| Amphibia | A14 | 54 | 0.45 | 53.95 | Andean | Mixed | Mixed | BcstDcst | 0.43 | NA | 0.4 | NA | Gradual |
| Amphibia | A15 | 110 | 0.7 | 33.89 | Andean | Mixed | Mixed | BcstDcst | 0.16 | NA | 0.06 | NA | Gradual |
| Amphibia | A16 | 159 | 0.52 | 73.05 | Andean | Tropical forest | Lowland-Montane | BcstDcst | 0.07 | NA | 0 | NA | Gradual |

**Table S7.** Summary results showing the best-fit model for each clade (among constant, time-, temperature- and Andean uplift-dependent models) and the derived species richness pattern (gradual increase [Sc. 1], exponential increase [Sc. 2], saturated increase [Sc. 3] and decline [Sc. 4]). For each model category, we fitted three models in which speciation (Birth "B") and/or extinction (Death "D") remain constant ("cst"), or vary ("Var.") continuously with time, with temperature changes, or with the elevation of the Andes. The value of speciation (lambda) and extinction (mu) rates is provided, as well as the value of the dependence between speciation and extinction rates with the environmental variables (alpha and beta values, respectively). For time-dependent models, $\alpha$ and $\beta > 0$ reflect decreasing speciation and extinction towards the present, respectively, while $\alpha$ and $\beta <0$ indicate the opposite, increasing speciation and extinction towards the present. For temperature models, $\alpha$ and $\beta > 0$ reflect decreasing speciation and extinction with decreasing temperatures, respectively, and conversely. For the uplift models, $\alpha$ and $\beta$ $< 0$ reflect increasing speciation and extinction with increasing Andean elevations, respectively, and conversely. Species richness patterns are derived from the diversification trend through time. For each phylogeny, the sampling fraction (*i.e.* number of species sampled from the total described), crown age (in million years ago), and number of species (#spp) is provided together with the main geographic (Andean-centered, Amazonian-centered, or other), altitudinal (lowland, montane, highland) and habitat distribution (tropical open vegetation, tropical forest vegetation or non-tropical) of the species in the clade. Mixed patterns could be observed in all categories. Abbreviations for model type; B = Birth; D = Death; cst = constant; Var. = variable; Temp. = Temperature; Ande = Andean Uplift model.

| **Taxa** | **Clade** | **# spp.** | **Sampling frac.** | **Crown age** | **Distribution** | **Habitat** | **Altitude** | **Model type** | **Model category** | **Lambda** | **Alpha** | **Mu** | **Beta** | **Richness pattern** |
|---|---|---|---|---|---|---|---|---|---|---|---|---|---|---|
| **Plants** | P1 | 13 | 0.12 | 2.88 | Mixed | Tropical forest | Lowland-Montane | BcstDTimeVar | Time | 2.08 | NA | 0.02 | 2.52 | Exponential |
| **Plants** | P2 | 38 | 0.28 | 0.94 | Andean | Tropical forest | Lowland-Montane | BcstDTimeVar | Time | 6.43 | NA | 0.15 | 5.64 | Exponential |
| **Plants** | P3 | 12 | 0.16 | 43.02 | Andean | Tropical forest | Lowland-Montane | BcstDcst | Constant | 0.15 | NA | 0.09 | NA | Gradual |
| **Plants** | P4 | 10 | 0.12 | 13.8 | Amazonian | Tropical forest | Lowland-Montane | BcstDcst | Constant | 1.02 | NA | 0.89 | NA | Gradual |
| **Plants** | P5 | 10 | 0.12 | 7.32 | Amazonian | Tropical forest | Lowland-Montane | BcstDcst | Constant | 1.35 | NA | 1.04 | NA | Gradual |

| | | | | | | | | | | | | | | |
|---|---|---|---|---|---|---|---|---|---|---|---|---|---|---|
| **Plants** | P6 | 14 | 0.45 | 5.82 | Andean | Open tropical | Lowland-Montane | BcstDcst | Constant | 0.47 | NA | 0 | NA | Gradual |
| **Plants** | P7 | 29 | 0.57 | 12.27 | Amazonian | Tropical forest | Lowland-Montane | BcstDcst | Constant | 0.29 | NA | 0 | NA | Gradual |
| **Plants** | P8 | 15 | 0.14 | 11.88 | Andean | Tropical forest | Lowland-Montane | BcstDTemp.Var | Temperature | 0.94 | NA | 0.13 | 0.41 | Exponential |
| **Plants** | P9 | 10 | 0.15 | 19.63 | Amazonian | Tropical forest | Lowland-Montane | BcstDcst | Constant | 0.18 | NA | 0 | NA | Gradual |
| **Plants** | P10 | 24 | 0.41 | 25.52 | Amazonian | Tropical forest | Lowland-Montane | BcstDcst | Constant | 0.14 | NA | 0 | NA | Gradual |
| **Plants** | P11 | 42 | 0.68 | 15.15 | Mixed | Tropical forest | Lowland-Montane | BcstDcst | Constant | 0.43 | NA | 0.24 | NA | Gradual |
| **Plants** | P12 | 15 | 0.35 | 13.04 | Mixed | Tropical forest | Lowland-Montane | BcstDcst | Constant | 0.3 | NA | 0.08 | NA | Gradual |
| **Plants** | P13 | 46 | 0.32 | 56.31 | Mixed | Tropical forest | Lowland-Montane | BTemp.VarDcst | Temperature | 0.21 | 0.06 | 0.31 | NA | Declining |
| **Plants** | P14 | 10 | 0.2 | 26 | Andean | Tropical forest | Lowland-Montane | BcstDcst | Constant | 0.43 | NA | 0.38 | NA | Gradual |
| **Plants** | P15 | 19 | 0.63 | 19.88 | Amazonian | Tropical forest | Lowland-Montane | BcstDcst | Constant | 0.19 | NA | 0.05 | NA | Gradual |
| **Plants** | P16 | 19 | 0.21 | 7.08 | Mixed | Tropical forest | Lowland-Montane | BcstDcst | Constant | 1.14 | NA | 0.69 | NA | Gradual |
| **Plants** | P17 | 18 | 0.18 | 22.42 | Mixed | Tropical forest | Lowland-Montane | BcstDcst | Constant | 0.27 | NA | 0.14 | NA | Gradual |
| **Plants** | P18 | 7 | 0.88 | 10.8 | Andean | Tropical forest | Lowland-Montane | BcstDcst | Constant | 0.11 | NA | 0 | NA | Gradual |
| **Plants** | P19 | 22 | 0.18 | 28.02 | Andean | Tropical forest | Lowland-Montane | BcstDTimeVar | Time | 0.21 | NA | 0 | 0.39 | Exponential |
| **Plants** | P20 | 29 | 0.72 | 20.25 | Andean | Tropical forest | Lowland-Montane | BcstDcst | Constant | 0.14 | NA | 0 | NA | Gradual |
| **Plants** | P21 | 103 | 0.41 | 20.77 | Amazonian | Tropical forest | Lowland-Montane | BTemp.VarDTemp.Var | Temperature | 0.27 | 0.07 | 8 | -1.26 | Declining |
| **Plants** | P22 | 19 | 0.43 | 17.64 | Andean | Tropical forest | Montane-Highland | BcstDcst | Constant | 0.17 | NA | 0 | NA | Gradual |
| **Plants** | P23 | 9 | 0.56 | 25.05 | Amazonian | Tropical forest | Lowland-Montane | BcstDcst | Constant | 0.13 | NA | 0.07 | NA | Gradual |
| **Plants** | P24 | 200 | 0.36 | 17.03 | Andean | Mixed | Mixed | BTimeVarDcst | Time | 1.64 | -0.29 | 0 | NA | Exponential |
| **Plants** | P25 | 24 | 0.89 | 19.21 | Other | Tropical forest | Lowland-Montane | BcstDcst | Constant | 0.14 | NA | 0 | NA | Gradual |
| **Plants** | P26 | 43 | 0.61 | 14.63 | Other | Tropical forest | Lowland-Montane | BcstDcst | Constant | 0.23 | NA | 0 | NA | Gradual |
| **Plants** | P27 | 19 | 0.54 | 4.67 | Other | Temperate | Lowland-Montane | BcstDcst | Constant | 0.74 | NA | 0.17 | NA | Gradual |
| **Plants** | P28 | 37 | 0.28 | 2.75 | Andean | Open tropical | Montane-Highland | BcstDTimeVar | Time | 2.18 | NA | 0 | 4.34 | Exponential |
| **Plants** | P29 | 13 | 0.93 | 15.29 | Amazonian | Tropical forest | Lowland-Montane | BTemp.VarDcst | Temperature | 0.02 | 0.32 | 0 | NA | Saturated |
| **Plants** | P30 | 14 | 0.88 | 37.32 | Other | Tropical forest | Lowland-Montane | BcstDcst | Constant | 0.1 | NA | 0.06 | NA | Gradual |

| | | | | | | | | | | | | | | |
|---|---|---|---|---|---|---|---|---|---|---|---|---|---|---|
| **Plants** | P31 | 78 | 0.98 | 30.97 | Other | Tropical forest | Lowland-Montane | BcstDcst | Constant | 0.19 | NA | 0.08 | NA | Gradual |
| **Plants** | P32 | 22 | 0.79 | 20.51 | Other | Tropical forest | Lowland-Montane | BcstDcst | Constant | 0.14 | NA | 0 | NA | Gradual |
| **Plants** | P33 | 16 | 1 | 19.96 | Other | Mixed | Lowland-Montane | BcstDcst | Constant | 0.09 | NA | 0 | NA | Gradual |
| **Plants** | P34 | 32 | 1 | 29.68 | Andean | Tropical forest | Lowland-Montane | BcstDcst | Constant | 0.21 | NA | 0.14 | NA | Gradual |
| **Plants** | P35 | 114 | 0.99 | 38.07 | Andean | Tropical forest | Lowland-Montane | BTemp.VarDcst | Temperature | 0.09 | 0.07 | 0 | NA | Saturated |
| **Plants** | P36 | 154 | 0.99 | 27.09 | Mixed | Mixed | Lowland-Montane | BcstDTemp.Var | Temperature | 0.21 | NA | 0 | 1.44 | Exponential |
| **Plants** | P37 | 178 | 0.99 | 43.62 | Mixed | Tropical forest | Lowland-Montane | BTemp.VarDTemp.Var | Temperature | 0.11 | 0.17 | 0 | 0.65 | Declining |
| **Plants** | P38 | 102 | 1 | 26.92 | Andean | Tropical forest | Lowland-Montane | BcstDTimeVar | Time | 0.25 | NA | 0.01 | 0.18 | Exponential |
| **Plants** | P39 | 22 | 1 | 34.21 | Mixed | Tropical forest | Lowland-Montane | BcstDcst | Constant | 0.26 | NA | 0.26 | NA | Gradual |
| **Plants** | P40 | 271 | 0.46 | 20.39 | Mixed | Mixed | Lowland-Montane | BTemp.VarDcst | Temperature | 0.9 | -0.19 | 0.01 | NA | Exponential |
| **Plants** | P41 | 670 | 0.13 | 17.87 | Andean | Tropical forest | Montane-Highland | BcstDcst | Constant | 3.08 | NA | 2.76 | NA | Gradual |
| **Plants** | P42 | 789 | 0.21 | 31.48 | Andean | Tropical forest | Mixed | BTemp.VarDcst | Temperature | 0.83 | -0.19 | 0 | NA | Exponential |
| **Plants** | P43 | 168 | 0.25 | 21.01 | Mixed | Open tropical | Lowland-Montane | BcstDTimeVar | Time | 0.71 | NA | 0.31 | 0.05 | Exponential |
| **Plants** | P44 | 126 | 0.33 | 6 | Mixed | Tropical forest | Lowland-Montane | BTimeVarDcst | Time | 3.76 | -0.68 | 0 | NA | Exponential |
| **Plants** | P45 | 588 | 0.49 | 48.45 | Andean | Tropical forest | Lowland-Montane | BcstDAnde.Var | Uplift | 0.17 | NA | 0.2 | 0 | Exponential |
| **Plants** | P46 | 111 | 0.79 | 28.92 | Amazonian | Tropical forest | Lowland-Montane | BTemp.VarDcst | Temperature | 0.09 | 0.11 | 0 | NA | Saturated |
| **Plants** | P47 | 21 | 1 | 28.86 | Other | Temperate | Lowland-Montane | BcstDcst | Constant | 0.34 | NA | 0.34 | NA | Gradual |
| **Plants** | P48 | 50 | 0.23 | 32.21 | Andean | Tropical forest | Lowland-Montane | BcstDcst | Constant | 3.13 | NA | 3.13 | NA | Gradual |
| **Plants** | P49 | 44 | 0.3 | 30.99 | Other | Temperate | Montane-Highland | BcstDcst | Constant | 1.18 | NA | 1.14 | NA | Gradual |
| **Plants** | P50 | 63 | 0.71 | 7.21 | Other | Temperate | Lowland-Montane | BcstDcst | Constant | 0.59 | NA | 0 | NA | Gradual |
| **Plants** | P51 | 189 | 0.44 | 21.79 | Mixed | Mixed | Lowland-Montane | BTemp.VarDcst | Temperature | 0.61 | -0.17 | 0 | NA | Exponential |
| **Plants** | P52 | 495 | 0.32 | 20.79 | Andean | Tropical forest | Mixed | BTimeVarDcst | Time | 0.59 | -0.07 | 0 | NA | Exponential |
| **Plants** | P53 | 86 | 0.77 | 30.28 | Amazonian | Tropical forest | Lowland-Montane | BcstDcst | Constant | 0.15 | NA | 0 | NA | Gradual |
| **Plants** | P54 | 38 | 0.56 | 45.01 | Other | Mixed | Lowland-Montane | BcstDTemp.Var | Temperature | 0.14 | NA | 0 | 0.79 | Declining |
| **Plants** | P55 | 120 | 0.46 | 19.5 | Amazonian | Tropical forest | Lowland-Montane | BTemp.VarDcst | Temperature | 1.7 | -0.38 | 0 | NA | Exponential |

| | | | | | | | | | | | | | | |
|---|---|---|---|---|---|---|---|---|---|---|---|---|---|---|
| **Plants** | P56 | 150 | 0.25 | 27.88 | Mixed | Mixed | Lowland-Montane | BTemp.VarDTemp.Var | Temperature | 0.09 | 0.47 | 0.1 | 0.46 | Declining |
| **Plants** | P57 | 161 | 0.25 | 48.28 | Mixed | Mixed | Lowland-Montane | BcstDcst | Constant | 0.18 | NA | 0.07 | NA | Gradual |
| **Plants** | P58 | 307 | 0.24 | 42.38 | Mixed | Mixed | Lowland-Montane | BTemp.VarDcst | Temperature | 0.51 | -0.13 | 0.08 | NA | Exponential |
| **Plants** | P59 | 34 | 0.35 | 24.17 | Amazonian | Tropical forest | Lowland-Montane | BcstDTimeVar | Time | 0.43 | NA | 0.08 | 0.15 | Exponential |
| **Plants** | P60 | 16 | 0.18 | 35.74 | Andean | Open tropical | Lowland-Montane | BcstDTimeVar | Time | 0.17 | NA | 0 | 0.25 | Exponential |
| **Plants** | P61 | 14 | 0.13 | 8.68 | Amazonian | Tropical forest | Lowland-Montane | BcstDcst | Constant | 0.51 | NA | 0.07 | NA | Gradual |
| **Plants** | P62 | 10 | 0.4 | 1.65 | Mixed | Tropical forest | Lowland-Montane | BcstDcst | Constant | 1.7 | NA | 0 | NA | Gradual |
| **Plants** | P63 | 10 | 0.42 | 0.88 | Other | Open tropical | Lowland-Montane | BcstDcst | Constant | 2.23 | NA | 0 | NA | Gradual |
| **Plants** | P64 | 10 | 0.59 | 4.57 | Other | Mixed | Lowland-Montane | BcstDcst | Constant | 0.42 | NA | 0 | NA | Gradual |
| **Plants** | P65 | 32 | 0.11 | 21.51 | Andean | Tropical forest | Montane-Highland | BcstDcst | Constant | 0.71 | NA | 0.55 | NA | Gradual |
| **Plants** | P66 | 48 | 0.28 | 0.5 | Mixed | Mixed | Lowland-Montane | BcstDcst | Constant | 8.82 | NA | 0 | NA | Gradual |
| **Mammals** | M1 | 32 | 1 | 67.96 | Amazonian | Mixed | Lowland-Montane | BcstDcst | Constant | 0.08 | NA | 0.05 | NA | Gradual |
| **Mammals** | M2 | 192 | 0.86 | 41.86 | Amazonian | Mixed | Lowland-Montane | BTemp.VarDTemp.Var | Temperature | 0.06 | 0.3 | 0.06 | 0.3 | Saturated |
| **Mammals** | M3 | 20 | 0.69 | 22.5 | Mixed | Tropical forest | Lowland-Montane | BTimeVarDTimeVar | Time | 0.01 | 0.36 | 0.03 | 0.33 | Declining |
| **Mammals** | M4 | 95 | 0.48 | 21.54 | Amazonian | Tropical forest | Lowland-Montane | BTemp.VarDcst | Temperature | 1.19 | -0.34 | 0 | NA | Exponential |
| **Mammals** | M5 | 43 | 0.42 | 25.49 | Amazonian | Tropical forest | Lowland-Montane | BcstDcst | Constant | 0.25 | NA | 0.12 | NA | Gradual |
| **Mammals** | M6 | 279 | 0.68 | 12.66 | Andean | Mixed | Mixed | BTemp.VarDcst | Temperature | 0.33 | 0.1 | 0 | NA | Saturated |
| **Mammals** | M7 | 11 | 0.69 | 10.6 | Other | Mixed | Lowland-Montane | BcstDcst | Constant | 0.17 | NA | 0 | NA | Gradual |
| **Mammals** | M8 | 10 | 0.46 | 13.81 | Andean | Mixed | Lowland-Montane | BcstDcst | Constant | 0.45 | NA | 0.41 | NA | Gradual |
| **Mammals** | M9 | 20 | 0.91 | 34.9 | Amazonian | Tropical forest | Lowland-Montane | BTimeVarDcst | Time | 0.02 | 0.06 | 0 | NA | Saturated |
| **Mammals** | M10 | 11 | 0.58 | 8.54 | Andean | Mixed | Mixed | BcstDcst | Constant | 0.23 | NA | 0 | NA | Gradual |
| **Mammals** | M11 | 10 | 1 | 16 | Amazonian | Mixed | Lowland-Montane | BcstDcst | Constant | 0.1 | NA | 0 | NA | Gradual |
| **Mammals** | M12 | 199 | 0.82 | 35.28 | Amazonian | Mixed | Lowland-Montane | BTemp.VarDTemp.Var | Temperature | 0.14 | 0.36 | 0.18 | 0.33 | Declining |
| **Birds** | B1 | 39 | 0.71 | 11.82 | Amazonian | Tropical forest | Lowland-Montane | BcstDcst | Constant | 0.33 | NA | 0 | NA | Gradual |
| **Birds** | B2 | 233 | 0.69 | 26.1 | Andean | Mixed | Mixed | BTemp.VarDcst | Temperature | 0.14 | 0.07 | 0.01 | NA | Saturated |

| | | | | | | | | | | | | | | |
|---|---|---|---|---|---|---|---|---|---|---|---|---|---|---|
| **Birds** | B3 | 8 | 0.73 | 24.79 | Mixed | Mixed | Lowland-Montane | BcstDcst | Constant | 0.3 | NA | 0.36 | NA | Gradual |
| **Birds** | B4 | 118 | 0.71 | 28.95 | Amazonian | Tropical forest | Mixed | BcstDcst | Constant | 0.18 | NA | 0.02 | NA | Gradual |
| **Birds** | B5 | 55 | 0.49 | 25.17 | Andean | Tropical forest | Lowland-Montane | BTemp.VarDcst | Temperature | 0.08 | 0.12 | 0 | NA | Saturated |
| **Birds** | B6 | 9 | 0.6 | 23.42 | Andean | Mixed | Lowland-Montane | BcstDcst | Constant | 0.09 | NA | 0 | NA | Gradual |
| **Birds** | B7 | 7 | 0.64 | 18.55 | Andean | Tropical forest | Lowland-Montane | BcstDcst | Constant | 0.2 | NA | 0.2 | NA | Gradual |
| **Birds** | B8 | 165 | 0.71 | 16.82 | Amazonian | Tropical forest | Lowland-Montane | BTemp.VarDcst | Temperature | 0.07 | 0.25 | 0 | NA | Saturated |
| **Birds** | B9 | 292 | 0.94 | 32.59 | Mixed | Mixed | Mixed | BTemp.VarDcst | Temperature | 0.11 | 0.07 | 0 | NA | Saturated |
| **Birds** | B10 | 316 | 0.79 | 25.5 | Mixed | Mixed | Mixed | BTemp.VarDcst | Temperature | 0.13 | 0.06 | 0 | NA | Saturated |
| **Birds** | B11 | 80 | 0.68 | 23.52 | Amazonian | Tropical forest | Lowland-Montane | BcstDcst | Constant | 0.18 | NA | 0 | NA | Gradual |
| **Birds** | B12 | 19 | 0.37 | 20.15 | Other | Tropical forest | Lowland-Montane | BcstDcst | Constant | 0.2 | NA | 0.04 | NA | Gradual |
| **Birds** | B13 | 36 | 0.95 | 17.38 | Andean | Mixed | Lowland-Montane | BTemp.VarDcst | Temperature | 0.06 | 0.2 | 0 | NA | Saturated |
| **Birds** | B14 | 39 | 0.85 | 8.2 | Andean | Mixed | Mixed | BTimeVarDcst | Time | 0.06 | 0.37 | 1 | NA | Saturated |
| **Birds** | B15 | 17 | 0.94 | 13.11 | Amazonian | Tropical forest | Lowland-Montane | BcstDcst | Constant | 0.14 | NA | 0 | NA | Gradual |
| **Birds** | B16 | 67 | 0.62 | 24.51 | Amazonian | Tropical forest | Lowland-Montane | BTemp.VarDcst | Temperature | 0.05 | 0.21 | 0 | NA | Saturated |
| **Birds** | B17 | 10 | 0.37 | 13.1 | Mixed | Mixed | Lowland-Montane | BcstDcst | Constant | 0.19 | NA | 0 | NA | Gradual |
| **Birds** | B18 | 89 | 0.99 | 11.21 | Mixed | Mixed | Lowland-Montane | BTemp.VarDcst | Temperature | 0.05 | 0.41 | 0.08 | NA | Declining |
| **Birds** | B19 | 92 | 0.89 | 12.02 | Andean | Mixed | Lowland-Montane | BTemp.VarDcst | Temperature | 0.1 | 0.27 | 0 | NA | Saturated |
| **Birds** | B20 | 41 | 0.82 | 18.04 | Andean | Mixed | Lowland-Montane | BTemp.VarDcst | Temperature | 0.02 | 0.34 | 0 | NA | Saturated |
| **Birds** | B21 | 309 | 0.77 | 13.57 | Other | Open tropical | Lowland-Montane | BTemp.VarDcst | Temperature | 0.15 | 0.3 | 0.37 | NA | Declining |
| **Birds** | B22 | 20 | 0.87 | 24.67 | Mixed | Mixed | Mixed | BcstDcst | Constant | 0.1 | NA | 0 | NA | Gradual |
| **Birds** | B23 | 25 | 0.76 | 9.37 | Amazonian | Tropical forest | Lowland-Montane | BTemp.VarDcst | Temperature | 0.06 | 0.38 | 0 | NA | Saturated |
| **Birds** | B24 | 10 | 0.91 | 6.01 | Amazonian | Tropical forest | Lowland-Montane | BcstDcst | Constant | 0.26 | NA | 0 | NA | Gradual |
| **Birds** | B25 | 12 | 0.5 | 10.54 | Mixed | Tropical forest | Lowland-Montane | BcstDcst | Constant | 0.21 | NA | 0 | NA | Gradual |
| **Birds** | B26 | 12 | 0.86 | 5.38 | Amazonian | Mixed | Lowland-Montane | BcstDcst | Constant | 0.32 | NA | 0 | NA | Gradual |
| **Birds** | B27 | 8 | 0.29 | 12.17 | Mixed | Tropical forest | Lowland-Montane | BcstDcst | Constant | 0.62 | NA | 0.56 | NA | Gradual |

| | | | | | | | | | | | | | | |
|---|---|---|---|---|---|---|---|---|---|---|---|---|---|---|
| **Birds** | B28 | 44 | 0.75 | 20.12 | Amazonian | Mixed | Lowland-Montane | BcstDcst | Constant | 0.22 | NA | 0.04 | NA | Gradual |
| **Birds** | B29 | 10 | 0.27 | 42.09 | Amazonian | Tropical forest | Lowland-Montane | BcstDcst | Constant | 0.12 | NA | 0.08 | NA | Gradual |
| **Birds** | B30 | 9 | 1 | 9.65 | Amazonian | Mixed | Lowland-Montane | BcstDcst | Constant | 0.15 | NA | 0 | NA | Gradual |
| **Birds** | B31 | 12 | 0.71 | 11.06 | Andean | Mixed | Mixed | BcstDcst | Constant | 0.2 | NA | 0 | NA | Gradual |
| **Birds** | B32 | 13 | 0.76 | 20.68 | Andean | Open tropical | Lowland-Montane | BcstDcst | Constant | 0.1 | NA | 0 | NA | Gradual |
| **Squamata** | S1 | 76 | 0.49 | 34.35 | Amazonian | Mixed | Lowland-Montane | BTemp.VarDcst | Temperature | 0.04 | 0.16 | 0 | NA | Saturated |
| **Squamata** | S2 | 15 | 0.38 | 26.03 | Andean | Mixed | Lowland-Montane | BTimeVarDcst | Time | 0.04 | 0.07 | 0 | NA | Saturated |
| **Squamata** | S3 | 19 | 0.24 | 21.78 | Andean | Mixed | Lowland-Montane | BcstDcst | Constant | 0.28 | NA | 0.13 | NA | Gradual |
| **Squamata** | S4 | 13 | 0.62 | 21.14 | Andean | Tropical forest | Lowland-Montane | BcstDcst | Constant | 0.1 | NA | 0 | NA | Gradual |
| **Squamata** | S5 | 43 | 0.61 | 21.41 | Andean | Mixed | Lowland-Montane | BTemp.VarDcst | Temperature | 0.08 | 0.16 | 0 | NA | Saturated |
| **Squamata** | S6 | 15 | 0.47 | 44.84 | Mixed | Mixed | Lowland-Montane | BcstDcst | Constant | 0.06 | NA | 0 | NA | Gradual |
| **Squamata** | S7 | 9 | 0.26 | 80.54 | Amazonian | Tropical forest | Lowland-Montane | BcstDcst | Constant | 0.23 | NA | 0.24 | NA | Gradual |
| **Squamata** | S8 | 29 | 0.49 | 43.74 | Amazonian | Tropical forest | Lowland-Montane | BTimeVarDTimeVar | Time | 0.58 | -0.06 | 1.65 | -0.29 | Declining |
| **Squamata** | S9 | 214 | 0.49 | 83.4 | Other | Tropical forest | Lowland-Montane | BAnde.VarDcst | Uplift | 0.14 | 0 | 0 | NA | Saturated |
| **Squamata** | S10 | 119 | 0.39 | 71.14 | Andean | Temperate | Montane-Highland | BAnde.VarDAnde.Var | Uplift | 0.1 | 0 | 0.18 | 0 | Exponential |
| **Squamata** | S11 | 11 | 0.33 | 42.19 | Other | Mixed | Lowland-Montane | BAnde.VarDcst | Uplift | 0.17 | 0 | 0 | NA | Saturated |
| **Squamata** | S12 | 12 | 0.44 | 81.21 | Amazonian | Tropical forest | Lowland-Montane | BAnde.VarDAnde.Var | Uplift | 0.3 | 0 | 0.66 | 0 | Exponential |
| **Squamata** | S13 | 44 | 0.43 | 19.79 | Other | Mixed | Mixed | BTimeVarDcst | Time | 0.12 | 0.05 | 0 | NA | Saturated |
| **Squamata** | S14 | 22 | 0.76 | 34.56 | Andean | Tropical forest | Lowland-Montane | BcstDcst | Constant | 0.12 | NA | 0.06 | NA | Gradual |
| **Squamata** | S15 | 78 | 0.57 | 88.47 | Andean | Mixed | Lowland-Montane | BAnde.VarDAnde.Var | Uplift | 0.09 | 0 | 0.28 | -0.01 | Exponential |
| **Squamata** | S16 | 18 | 0.42 | 26.47 | Other | Tropical forest | Lowland-Montane | BcstDcst | Constant | 0.11 | NA | 0.01 | NA | Gradual |
| **Squamata** | S17 | 37 | 0.39 | 48.84 | Amazonian | Mixed | Lowland-Montane | BTemp.VarDcst | Temperature | 0.01 | 0.22 | 0 | NA | Saturated |
| **Squamata** | S18 | 144 | 0.43 | 86.27 | Mixed | Mixed | Mixed | BAnde.VarDAnde.Var | Uplift | 0.2 | 1 | 0.17 | 0 | Exponential |
| **Squamata** | S19 | 20 | 0.33 | 25.22 | Andean | Mixed | Lowland-Montane | BTemp.VarDcst | Temperature | 0.03 | 0.29 | 0.06 | NA | Declining |
| **Squamata** | S20 | 16 | 0.84 | 36.33 | Andean | Tropical forest | Lowland-Montane | BAnde.VarDcst | Uplift | 0.17 | 0 | 0 | NA | Saturated |

| | | | | | | | | | | | | | | |
|---|---|---|---|---|---|---|---|---|---|---|---|---|---|---|
| **Squamata** | S21 | 20 | 0.31 | 66.91 | Andean | Mixed | Lowland-Montane | BcstDcst | Constant | 0.06 | NA | 0 | NA | Gradual |
| **Squamata** | S22 | 69 | 0.41 | 70.89 | Amazonian | Tropical forest | Lowland-Montane | BAnde.VarDcst | Uplift | 0.08 | 0 | 0 | NA | Saturated |
| **Squamata** | S23 | 60 | 0.49 | 34.42 | Other | Mixed | Mixed | BTemp.VarDcst | Temperature | 0.05 | 0.12 | 0 | NA | Saturated |
| **Squamata** | S24 | 45 | 0.14 | 29.25 | Andean | Tropical forest | Lowland-Montane | BTemp.VarDcst | Temperature | 0.06 | 0.15 | 0 | NA | Saturated |
| **Amphibia** | A1 | 118 | 0.93 | 67.18 | Mixed | Tropical forest | Lowland-Montane | BAnde.VarDAnde.Var | Uplift | 0.18 | 0 | 0.16 | 0 | Exponential |
| **Amphibia** | A2 | 136 | 1 | 67.35 | Andean | Tropical forest | Mixed | BTemp.VarDcst | Temperature | 0.04 | 0.06 | 0 | NA | Saturated |
| **Amphibia** | A3 | 86 | 0.79 | 80.69 | Andean | Tropical forest | Lowland-Montane | BcstDcst | Constant | 0.04 | NA | 0 | NA | Gradual |
| **Amphibia** | A4 | 170 | 0.78 | 72.08 | Other | Tropical forest | Mixed | BAnde.VarDAnde.Var | Uplift | 0.76 | 0 | 0.8 | 0 | Exponential |
| **Amphibia** | A5 | 61 | 0.47 | 69.87 | Other | Tropical forest | Lowland-Montane | BAnde.VarDAnde.Var | Uplift | 0.22 | 0 | 0.21 | 0 | Exponential |
| **Amphibia** | A6 | 295 | 0.47 | 68.16 | Andean | Tropical forest | Mixed | BAnde.VarDAnde.Var | Uplift | 0.17 | 0 | 0.11 | 0 | Exponential |
| **Amphibia** | A7 | 50 | 0.79 | 54.78 | Amazonian | Tropical forest | Lowland-Montane | BAnde.VarDcst | Uplift | 0.08 | 0 | 0 | NA | Saturated |
| **Amphibia** | A8 | 117 | 0.64 | 69.93 | Amazonian | Tropical forest | Lowland-Montane | BcstDcst | Constant | 0.09 | NA | 0.03 | NA | Gradual |
| **Amphibia** | A9 | 147 | 0.58 | 72.8 | Amazonian | Tropical forest | Lowland-Montane | BcstDcst | Constant | 0.07 | NA | 0 | NA | Gradual |
| **Amphibia** | A10 | 60 | 0.71 | 43.33 | Amazonian | Tropical forest | Lowland-Montane | BTemp.VarDcst | Temperature | 0.04 | 0.16 | 0.06 | NA | Declining |
| **Amphibia** | A11 | 127 | 0.49 | 87.37 | Mixed | Mixed | Lowland-Montane | BcstDAnde.Var | Uplift | 0.25 | NA | 3 | 0 | Declining |
| **Amphibia** | A12 | 186 | 0.88 | 78.07 | Mixed | Mixed | Lowland-Montane | BAnde.VarDAnde.Var | Uplift | 0.1 | 1 | 0.08 | 0 | Exponential |
| **Amphibia** | A13 | 128 | 0.81 | 33.4 | Andean | Tropical forest | Lowland-Montane | BTemp.VarDcst | Temperature | 0.03 | 0.2 | 0 | NA | Saturated |
| **Amphibia** | A14 | 54 | 0.45 | 53.95 | Andean | Mixed | Mixed | BcstDcst | Constant | 0.43 | NA | 0.4 | NA | Gradual |
| **Amphibia** | A15 | 110 | 0.7 | 33.89 | Andean | Mixed | Mixed | BcstDcst | Constant | 0.16 | NA | 0.06 | NA | Gradual |
| **Amphibia** | A16 | 159 | 0.52 | 73.05 | Andean | Tropical forest | Lowland-Montane | BAnde.VarDAnde.Var | Uplift | 0.19 | 0 | 0.17 | 0 | Exponential |

**Table S8.** Number of phylogenies and species supporting different speciation trends (among constant, decreasing, increasing), species richness dynamics (gradual increase [Sc. 1], exponential increase [Sc. 2], saturated increase [Sc. 3] and decline [Sc. 4]), and diversification drivers (time, temperature, Andean uplift) in this study.

| | | All | Plants | Mammals | Birds | Squamata | Amphibia |
|---|---|---|---|---|---|---|---|
| | Total # of clades | 150 | 66 | 12 | 32 | 24 | 16 |
| | Total # of species | 12512 | 6222 | 922 | 2216 | 1148 | 2004 |
| **Diversification trend** | # clades constant | 76 | 39 | 6 | 19 | 7 | 5 |
| | # clades time-variable | 74 | 27 | 6 | 13 | 17 | 11 |
| | # species constant | 2989 | 1785 | 117 | 457 | 116 | 514 |
| | # species time-variable | 9523 | 4437 | 805 | 1759 | 1032 | 1490 |
| **Speciation trend** | # clades decreasing | 39 | 7 | 5 | 13 | 10 | 4 |
| | # clades increasing | 22 | 8 | 1 | 0 | 7 | 6 |
| | # clades lambda constant (mu varies) | 13 | 12 | 0 | 0 | 0 | 1 |
| **Species richness dynamic** | Gradual increase | 76 | 39 | 6 | 19 | 7 | 5 |
| | Exponential increase | 30 | 19 | 1 | 0 | 4 | 6 |
| | Saturated increase | 31 | 3 | 3 | 11 | 11 | 3 |
| | Declining & Waning | 13 | 5 | 2 | 2 | 2 | 2 |
| **Environmental driver** | Time-dependent models | 17 | 11 | 2 | 1 | 3 | 0 |
| | Temperature-dependent models | 40 | 15 | 4 | 12 | 6 | 3 |
| | Uplift-dependent models | 17 | 1 | 0 | 0 | 8 | 8 |

**Table S9.** Summary results showing the best-fit model for each clade based on Pulled diversification rates (PDR) in *castor*. For each model category, we compared a constant model (1-time interval) with a model in which PDR values are allowed to vary independently on a grid of 3-time intervals (PDR values are presented for the successive intervals going back in time.). We selected the model that best explains the LTT of the Neotropical time trees based on AIC values. Rholambda ($\rho\lambda o$) value is the product of present-day speciation ($\lambda o$) and sampling fraction ($\rho$). Knowing $\rho$, $\lambda_0$ could be derived as follows: $\lambda_0 = \lambda_0\rho/\rho$. Pulled extinction rates "PER" ($\mu_p$) for each time interval could be derived as follows: $\mu_p := \lambda_0 - r_p$. If the estimated $\mu_p(0)$ is negative, this is evidence that speciation (*lambda*) is currently decreasing over time (91, 92).

| | | **Constant model** | | | | **Gridded model** | | | | | | | | | | |
|---|---|---|---|---|---|---|---|---|---|---|---|---|---|---|---|---|
| **Taxa** | **Clade** | **PDR** | **rholambda0** | **Lambda** | **AIC** | **PDR1.present** | **PDR2** | **PDR3.past** | **rholambda0** | **lambda0** | **PER1.present** | **PER2** | **PER3.past** | **Lambda trend** | **AIC** | **Best model** |
| **Plants** | P1 | 1.83 | 0.21 | 1.79 | 33.1 | 20.44 | -0.14 | 1.89 | 0 | 0 | -20.44 | 0.14 | -1.89 | lambda descreases | 30.99 | time-variable |
| **Plants** | P2 | 5.63 | 1.76 | 6.32 | -12.61 | 20.24 | 1.91 | 5.8 | 0.54 | 1.94 | -19.7 | -1.36 | -5.26 | lambda descreases | -12.97 | time-variable |
| **Plants** | P3 | 0.09 | 0.02 | 0.13 | 93.34 | 0.52 | -0.06 | 0.26 | 0 | 0.02 | -0.52 | 0.07 | -0.25 | lambda descreases | 94.92 | constant |
| **Plants** | P4 | 0.22 | 0.1 | 0.82 | 56.25 | 1.92 | -0.43 | 0.9 | 0.01 | 0.08 | -1.92 | 0.44 | -0.89 | lambda descreases | 56.68 | constant |
| **Plants** | P5 | 0.45 | 0.13 | 1.07 | 46.05 | 1.67 | -0.08 | 1.1 | 0.06 | 0.47 | -1.61 | 0.14 | -1.05 | lambda descreases | 49.35 | constant |
| **Plants** | P6 | 0.67 | 0.14 | 0.32 | 56.55 | 3.92 | -0.45 | 2.08 | 0.02 | 0.04 | -3.9 | 0.47 | -2.06 | lambda descreases | 57.5 | constant |
| **Plants** | P7 | 0.34 | 0.14 | 0.25 | 141.95 | 0.5 | 0.27 | 0.43 | 0.12 | 0.22 | -0.38 | -0.15 | -0.3 | lambda descreases | 145.86 | constant |
| **Plants** | P8 | 0.23 | 0.22 | 1.62 | 73.25 | 1.72 | -0.45 | 0.95 | 0.06 | 0.42 | -1.66 | 0.51 | -0.89 | lambda descreases | 73.35 | constant |
| **Plants** | P9 | 0.23 | 0.02 | 0.12 | 63.98 | 0.56 | 0.09 | 0.44 | 0.01 | 0.06 | -0.55 | -0.08 | -0.42 | lambda descreases | 67.63 | constant |
| **Plants** | P10 | 0.17 | 0.04 | 0.1 | 157.29 | 0.44 | 0.08 | 0.27 | 0.02 | 0.05 | -0.42 | -0.06 | -0.24 | lambda descreases | 160.61 | constant |
| **Plants** | P11 | 0.21 | 0.28 | 0.41 | 201.43 | 0.25 | 0.15 | 0.38 | 0.28 | 0.4 | 0.02 | 0.12 | -0.1 | unknown | 205.24 | constant |
| **Plants** | P12 | 0.27 | 0.09 | 0.26 | 81.99 | 0.82 | 0.02 | 0.61 | 0.05 | 0.14 | -0.77 | 0.03 | -0.56 | lambda descreases | 85.27 | constant |
| **Plants** | P13 | 0.04 | 0.05 | 0.16 | 361.3 | 0.02 | -0.01 | 0.23 | 0.06 | 0.2 | 0.04 | 0.07 | -0.17 | unknown | 359.39 | time-variable |
| **Plants** | P14 | 0.1 | 0.07 | 0.34 | 67 | 0.52 | -0.09 | 0.31 | 0.03 | 0.14 | -0.49 | 0.12 | -0.28 | lambda descreases | 69.92 | constant |

| | | | | | | | | | | | | | | | | |
|---|---|---|---|---|---|---|---|---|---|---|---|---|---|---|---|---|
| **Plants** | P15 | 0.17 | 0.1 | 0.17 | 112.7 | 0.55 | 0 | 0.35 | 0.06 | 0.09 | -0.5 | 0.06 | -0.29 | lambda descreases | 115.72 | constant |
| **Plants** | P16 | 0.53 | 0.21 | 1.01 | 77.97 | 1.08 | 0.39 | 0.51 | 0.14 | 0.7 | -0.94 | -0.24 | -0.37 | lambda descreases | 81.69 | constant |
| **Plants** | P17 | 0.15 | 0.04 | 0.24 | 119.47 | 0.09 | 0.03 | 0.59 | 0.06 | 0.36 | -0.03 | 0.03 | -0.52 | lambda descreases | 120.69 | constant |
| **Plants** | P18 | 0.46 | 0.02 | 0.02 | 37.11 | 5.23 | 0.02 | 0.66 | 0 | 0 | -5.23 | -0.02 | -0.66 | lambda descreases | 39.55 | constant |
| **Plants** | P19 | 0.18 | 0.04 | 0.21 | 144.08 | 0.62 | 0.1 | 0.13 | 0.01 | 0.06 | -0.61 | -0.09 | -0.12 | lambda descreases | 145.99 | constant |
| **Plants** | P20 | 0.18 | 0.09 | 0.12 | 174.5 | 0.17 | 0.13 | 0.35 | 0.1 | 0.13 | -0.07 | -0.04 | -0.25 | lambda descreases | 178.09 | constant |
| **Plants** | P21 | 0.38 | 0.02 | 0.04 | 584.39 | -0.19 | 0.52 | 0.19 | 0.05 | 0.12 | 0.24 | -0.47 | -0.14 | unknown | 583.32 | time-variable |
| **Plants** | P22 | 0.27 | 0.04 | 0.09 | 113.05 | 0.68 | 0.17 | 0.32 | 0.02 | 0.04 | -0.66 | -0.15 | -0.31 | lambda descreases | 116.51 | constant |
| **Plants** | P23 | 0.11 | 0.06 | 0.1 | 60.65 | 0.24 | 0.07 | 0.12 | 0.04 | 0.07 | -0.2 | -0.03 | -0.08 | lambda descreases | 64.56 | constant |
| **Plants** | P24 | 0.35 | 1 | 2.76 | 550.77 | 1.18 | 0.03 | -0.01 | 0.68 | 1.87 | -0.5 | 0.65 | 0.69 | lambda descreases | 533.76 | time-variable |
| **Plants** | P25 | 0.22 | 0.08 | 0.09 | 140.25 | 0.99 | -0.02 | 0.42 | 0.02 | 0.02 | -0.97 | 0.04 | -0.4 | lambda descreases | 140.97 | constant |
| **Plants** | P26 | 0.3 | 0.09 | 0.15 | 228.57 | 0.63 | 0.12 | 0.7 | 0.07 | 0.11 | -0.56 | -0.06 | -0.63 | lambda descreases | 231.23 | constant |
| **Plants** | P27 | 0.69 | 0.36 | 0.66 | 64.12 | 0.81 | 0.39 | 1.58 | 0.38 | 0.69 | -0.44 | -0.02 | -1.21 | lambda descreases | 67.7 | constant |
| **Plants** | P28 | 2.1 | 0.56 | 1.97 | 62.72 | 9.06 | 0.55 | 1.89 | 0.1 | 0.36 | -8.96 | -0.44 | -1.79 | lambda descreases | 59.82 | time-variable |
| **Plants** | P29 | 0.32 | 0.03 | 0.03 | 71.56 | 3.16 | 0.21 | -0.14 | 0 | 0 | -3.16 | -0.21 | 0.14 | lambda descreases | 70.36 | time-variable |
| **Plants** | P30 | 0.06 | 0.08 | 0.08 | 99.9 | 0.3 | -0.1 | 0.31 | 0.04 | 0.05 | -0.26 | 0.14 | -0.26 | lambda descreases | 101.97 | constant |
| **Plants** | P31 | 0.12 | 0.18 | 0.18 | 454.1 | 0.37 | -0.05 | 0.41 | 0.12 | 0.12 | -0.25 | 0.17 | -0.29 | lambda descreases | 453.02 | time-variable |
| **Plants** | P32 | 0.2 | 0.08 | 0.1 | 131.65 | 0.64 | 0.08 | 0.27 | 0.03 | 0.04 | -0.6 | -0.04 | -0.24 | lambda descreases | 134.44 | constant |
| **Plants** | P33 | 0.15 | 0.06 | 0.06 | 103.23 | -0.26 | 0.2 | 0.32 | 0.14 | 0.14 | 0.4 | -0.06 | -0.18 | unknown | 104.87 | constant |
| **Plants** | P34 | 0.09 | 0.19 | 0.19 | 190.01 | -0.1 | 0.18 | -0.01 | 0.25 | 0.25 | 0.35 | 0.07 | 0.26 | unknown | 193.08 | constant |
| **Plants** | P35 | 0.18 | 0.09 | 0.09 | 685.12 | 0.37 | 0.16 | 0 | 0.06 | 0.06 | -0.31 | -0.1 | 0.06 | lambda descreases | 682.02 | time-variable |
| **Plants** | P36 | 0.2 | 0.19 | 0.19 | 810.35 | 0.38 | 0.15 | 0.13 | 0.14 | 0.15 | -0.24 | 0 | 0.01 | lambda descreases | 811.29 | constant |
| **Plants** | P37 | 0.19 | 0.17 | 0.17 | 970.84 | 0.46 | 0.11 | 0 | 0.1 | 0.1 | -0.36 | 0 | 0.1 | lambda descreases | 959.66 | time-variable |
| **Plants** | P38 | 0.21 | 0.25 | 0.25 | 501.35 | 0.64 | 0.07 | 0.13 | 0.14 | 0.14 | -0.49 | 0.08 | 0.01 | lambda descreases | 496.5 | time-variable |
| **Plants** | P39 | 0.04 | 0.24 | 0.24 | 129.42 | 0.28 | -0.15 | 0.34 | 0.17 | 0.17 | -0.11 | 0.32 | -0.17 | lambda descreases | 131.19 | constant |

| | | | | | | | | | | | | | | | |
|---|---|---|---|---|---|---|---|---|---|---|---|---|---|---|---|
| **Plants** | P40 | 0.29 | 0.4 | 0.88 | 1112.4 | 0.5 | 0.26 | 0 | 0.34 | 0.74 | -0.16 | 0.08 | 0.34 | lambda descreases | 1110.57 | time-variable |
| **Plants** | P41 | 0.32 | 0.4 | 3.07 | 2712.49 | 0.28 | 0.32 | 0.35 | 0.42 | 3.16 | 0.13 | 0.09 | 0.06 | unknown | 2716.24 | constant |
| **Plants** | P42 | 0.24 | 0.18 | 0.87 | 4017.59 | 0.5 | 0.15 | 0.24 | 0.13 | 0.62 | -0.36 | -0.02 | -0.11 | lambda descreases | 4001.58 | time-variable |
| **Plants** | P43 | 0.29 | 0.2 | 0.83 | 805.84 | 0.48 | 0.25 | 0.16 | 0.16 | 0.66 | -0.32 | -0.09 | 0 | lambda descreases | 807.6 | constant |
| **Plants** | P44 | 1.25 | 1.58 | 4.77 | 164.19 | 5.17 | -0.48 | 1.14 | 0.69 | 2.08 | -4.49 | 1.17 | -0.46 | lambda descreases | 137.61 | time-variable |
| **Plants** | P45 | 0.2 | 0.06 | 0.13 | 3643.96 | 0.49 | 0.12 | 0.11 | 0.03 | 0.06 | -0.46 | -0.09 | -0.08 | lambda descreases | 3602.6 | time-variable |
| **Plants** | P46 | 0.24 | 0.08 | 0.1 | 645.59 | 1.12 | 0.02 | 0.21 | 0.01 | 0.01 | -1.11 | -0.01 | -0.2 | lambda descreases | 617.31 | time-variable |
| **Plants** | P47 | 0.03 | 0.32 | 0.32 | 117.12 | 0.3 | -0.36 | 0.73 | 0.25 | 0.25 | -0.05 | 0.6 | -0.48 | lambda descreases | 115.93 | time-variable |
| **Plants** | P48 | 0.02 | 0.69 | 3.05 | 229.64 | -0.19 | 0.06 | 0.19 | 0.83 | 3.67 | 1.02 | 0.78 | 0.64 | unknown | 229.73 | constant |
| **Plants** | P49 | 0.06 | 0.34 | 1.13 | 232.38 | 0.25 | -0.11 | 0.38 | 0.28 | 0.92 | 0.02 | 0.39 | -0.1 | unknown | 233.53 | constant |
| **Plants** | P50 | 0.65 | 0.39 | 0.55 | 211.5 | 1.36 | 0.4 | 0.74 | 0.28 | 0.39 | -1.09 | -0.13 | -0.46 | lambda descreases | 214.26 | constant |
| **Plants** | P51 | 0.21 | 0.26 | 0.58 | 929.61 | 0.6 | 0 | 0.48 | 0.17 | 0.39 | -0.43 | 0.17 | -0.31 | lambda descreases | 923.56 | time-variable |
| **Plants** | P52 | 0.35 | 0.22 | 0.7 | 2245.43 | 0.74 | 0.22 | 0.29 | 0.15 | 0.47 | -0.59 | -0.07 | -0.14 | lambda descreases | 2233.17 | time-variable |
| **Plants** | P53 | 0.05 | 0.17 | 0.26 | 765.15 | 0.11 | 0.02 | 0.12 | 0.15 | 0.23 | 0.04 | 0.13 | 0.03 | unknown | 767.83 | constant |
| **Plants** | P54 | 0.07 | 0.1 | 0.18 | 259.13 | 0.27 | -0.04 | 0.21 | 0.06 | 0.11 | -0.21 | 0.1 | -0.15 | lambda descreases | 260.25 | constant |
| **Plants** | P55 | 0.18 | 0.72 | 1.58 | 437.08 | 0.71 | -0.15 | 0.52 | 0.53 | 1.15 | -0.18 | 0.67 | 0 | lambda descreases | 433.25 | time-variable |
| **Plants** | P56 | 0.2 | 0.06 | 0.25 | 923.14 | 0.8 | -0.01 | 0.5 | 0.02 | 0.07 | -0.78 | 0.02 | -0.49 | lambda descreases | 906.15 | time-variable |
| **Plants** | P57 | 0.12 | 0.04 | 0.18 | 1135 | 0.27 | 0.06 | 0.23 | 0.03 | 0.12 | -0.24 | -0.03 | -0.2 | lambda descreases | 1134.35 | time-variable |
| **Plants** | P58 | 0.12 | 0.13 | 0.52 | 1887.12 | 0.33 | 0.02 | 0.33 | 0.08 | 0.35 | -0.25 | 0.07 | -0.25 | lambda descreases | 1877.14 | time-variable |
| **Plants** | P59 | 0.17 | 0.2 | 0.57 | 181.13 | 0.93 | -0.14 | 0.3 | 0.07 | 0.2 | -0.86 | 0.21 | -0.23 | lambda descreases | 177.28 | time-variable |
| **Plants** | P60 | 0.1 | 0.05 | 0.27 | 113.26 | 0.18 | 0.1 | 0.04 | 0.04 | 0.2 | -0.14 | -0.06 | -0.01 | lambda descreases | 116.97 | constant |
| **Plants** | P61 | 0.5 | 0.06 | 0.44 | 68.04 | 3.38 | -0.38 | 1.55 | 0 | 0.02 | -3.38 | 0.38 | -1.54 | lambda descreases | 68.39 | constant |
| **Plants** | P62 | 2.45 | 0.49 | 1.22 | 18.1 | 27.41 | -2.59 | 5.8 | 0 | 0.01 | -27.41 | 2.6 | -5.8 | lambda descreases | 17.43 | time-variable |
| **Plants** | P63 | 4.25 | 0.34 | 0.82 | 10.73 | 48.83 | -20.92 | 42.72 | 0 | 0.01 | -48.82 | 20.93 | -42.72 | lambda descreases | -0.26 | time-variable |
| **Plants** | P64 | 0.67 | 0.16 | 0.27 | 39.85 | 5.53 | -2.37 | 5.17 | 0.02 | 0.03 | -5.51 | 2.39 | -5.15 | lambda descreases | 37.54 | time-variable |

| | | | | | | | | | | | | | | | | |
|---|---|---|---|---|---|---|---|---|---|---|---|---|---|---|---|---|
| **Plants** | P65 | 0.18 | 0.07 | 0.66 | 198.46 | 0.12 | 0.14 | 0.34 | 0.08 | 0.79 | -0.04 | -0.06 | -0.26 | lambda descreases | 201.77 | constant |
| **Plants** | P66 | 9.67 | 2.13 | 7.71 | -60.39 | 21.97 | 4.33 | 18.19 | 1.3 | 4.72 | -20.67 | -3.02 | -16.89 | lambda descreases | -57.8 | constant |
| **Mammals** | M1 | 0.04 | 0.07 | 0.07 | 247.54 | 0.03 | 0.02 | 0.09 | 0.08 | 0.08 | 0.05 | 0.06 | -0.01 | unknown | 251.07 | constant |
| **Mammals** | M2 | 0.11 | 0.11 | 0.11 | 1245.12 | 0.3 | 0.04 | 0.17 | 0.07 | 0.07 | -0.23 | 0.03 | -0.1 | lambda descreases | 1240.03 | time-variable |
| **Mammals** | M3 | 0.19 | 0.02 | 0.03 | 133.22 | 0.99 | -0.12 | 0.57 | 0 | 0 | -0.99 | 0.12 | -0.56 | lambda descreases | 133.08 | time-variable |
| **Mammals** | M4 | 0.11 | 0.5 | 1.05 | 422.38 | 0.5 | -0.23 | 0.77 | 0.38 | 0.8 | -0.12 | 0.61 | -0.39 | lambda descreases | 416.13 | time-variable |
| **Mammals** | M5 | 0.14 | 0.1 | 0.24 | 265.58 | 0.46 | -0.04 | 0.45 | 0.06 | 0.14 | -0.4 | 0.1 | -0.39 | lambda descreases | 266.33 | constant |
| **Mammals** | M6 | 0.59 | 0.2 | 0.3 | 1103.75 | 1.06 | 0.47 | 0.48 | 0.14 | 0.21 | -0.92 | -0.33 | -0.34 | lambda descreases | 1102.91 | time-variable |
| **Mammals** | M7 | 0.4 | 0.04 | 0.06 | 57.98 | 3.73 | -0.42 | 1.3 | 0 | 0 | -3.73 | 0.42 | -1.3 | lambda descreases | 58.11 | constant |
| **Mammals** | M8 | 0.12 | 0.17 | 0.37 | 56.46 | 0.42 | -0.36 | 1.18 | 0.16 | 0.34 | -0.26 | 0.52 | -1.02 | lambda descreases | 57.66 | constant |
| **Mammals** | M9 | 0.14 | 0.01 | 0.01 | 146.61 | 1.79 | -0.14 | 0.43 | 0 | 0 | -1.79 | 0.14 | -0.43 | lambda descreases | 141.37 | time-variable |
| **Mammals** | M10 | 0.52 | 0.04 | 0.08 | 53.91 | -0.69 | 0.78 | 0.57 | 0.15 | 0.26 | 0.84 | -0.62 | -0.42 | unknown | 56.87 | constant |
| **Mammals** | M11 | 0.14 | 0.09 | 0.09 | 56.08 | 1.69 | -0.25 | 0.27 | 0 | 0 | -1.68 | 0.26 | -0.27 | lambda descreases | 56.15 | constant |
| **Mammals** | M12 | 0.14 | 0.21 | 0.26 | 1091.66 | 0.32 | 0.05 | 0.21 | 0.16 | 0.2 | -0.16 | 0.12 | -0.05 | lambda descreases | 1090.5 | time-variable |
| **Birds** | B1 | 0.4 | 0.2 | 0.29 | 174.19 | 2.11 | -0.17 | 0.68 | 0.04 | 0.06 | -2.07 | 0.21 | -0.64 | lambda descreases | 168.55 | time-variable |
| **Birds** | B2 | 0.22 | 0.1 | 0.14 | 1337.43 | 0.36 | 0.15 | 0.32 | 0.08 | 0.12 | -0.28 | -0.07 | -0.24 | lambda descreases | 1339.49 | constant |
| **Birds** | B3 | 0.03 | 0.16 | 0.22 | 50.77 | 0.17 | -0.24 | 0.57 | 0.15 | 0.21 | -0.01 | 0.39 | -0.41 | lambda descreases | 53.32 | constant |
| **Birds** | B4 | 0.17 | 0.13 | 0.18 | 681.82 | 0.45 | 0.06 | 0.26 | 0.08 | 0.12 | -0.37 | 0.02 | -0.18 | lambda descreases | 681.19 | time-variable |
| **Birds** | B5 | 0.24 | 0.04 | 0.08 | 340.15 | 0.58 | 0.17 | 0.23 | 0.02 | 0.03 | -0.56 | -0.15 | -0.22 | lambda descreases | 342.05 | constant |
| **Birds** | B6 | 0.15 | 0.04 | 0.06 | 59.97 | 1.32 | -0.15 | 0.4 | 0 | 0 | -1.32 | 0.16 | -0.4 | lambda descreases | 61.16 | constant |
| **Birds** | B7 | 0.1 | 0.09 | 0.14 | 44.06 | 0.36 | -0.13 | 0.47 | 0.07 | 0.11 | -0.29 | 0.2 | -0.4 | lambda descreases | 47.6 | constant |
| **Birds** | B8 | 0.39 | 0.06 | 0.08 | 860.41 | 0.71 | 0.26 | 0.74 | 0.04 | 0.06 | -0.67 | -0.22 | -0.7 | lambda descreases | 861.51 | constant |
| **Birds** | B9 | 0.19 | 0.11 | 0.12 | 1701.47 | 0.48 | 0.07 | 0.34 | 0.07 | 0.07 | -0.41 | 0 | -0.27 | lambda descreases | 1692.72 | time-variable |
| **Birds** | B10 | 0.22 | 0.11 | 0.14 | 1790.59 | 0.35 | 0.16 | 0.31 | 0.09 | 0.11 | -0.27 | -0.07 | -0.22 | lambda descreases | 1792.32 | constant |
| **Birds** | B11 | 0.2 | 0.11 | 0.16 | 455.91 | 0.23 | 0.19 | 0.14 | 0.1 | 0.15 | -0.13 | -0.09 | -0.04 | lambda descreases | 459.78 | constant |

| | | | | | | | | | | | | | | | | |
|---|---|---|---|---|---|---|---|---|---|---|---|---|---|---|---|---|
| **Birds** | B12 | 0.18 | 0.07 | 0.18 | 117.74 | 0.66 | -0.05 | 0.52 | 0.03 | 0.08 | -0.63 | 0.08 | -0.49 | lambda descreases | 119.92 | constant |
| **Birds** | B13 | 0.29 | 0.06 | 0.07 | 202.86 | 1.35 | -0.02 | 0.56 | 0.01 | 0.01 | -1.34 | 0.03 | -0.55 | lambda descreases | 200.96 | time-variable |
| **Birds** | B14 | 0.7 | 0.03 | 0.03 | 173.07 | 1.33 | 0.3 | 1.56 | 0.02 | 0.02 | -1.31 | -0.27 | -1.54 | lambda descreases | 174.58 | constant |
| **Birds** | B15 | 0.32 | 0.05 | 0.05 | 94.43 | 0.5 | 0.21 | 0.52 | 0.04 | 0.04 | -0.46 | -0.16 | -0.48 | lambda descreases | 98.23 | constant |
| **Birds** | B16 | 0.3 | 0.04 | 0.06 | 391.7 | 1.26 | 0.12 | 0.32 | 0 | 0.01 | -1.26 | -0.11 | -0.31 | lambda descreases | 384.19 | time-variable |
| **Birds** | B17 | 0.35 | 0.03 | 0.07 | 56.36 | 2.5 | -0.14 | 0.88 | 0 | 0 | -2.5 | 0.14 | -0.88 | lambda descreases | 58.25 | constant |
| **Birds** | B18 | 0.51 | 0.07 | 0.07 | 421.37 | 0.63 | 0.41 | 0.81 | 0.06 | 0.06 | -0.57 | -0.34 | -0.74 | lambda descreases | 424.64 | constant |
| **Birds** | B19 | 0.53 | 0.1 | 0.11 | 418.1 | 1.03 | 0.43 | 0.39 | 0.06 | 0.06 | -0.98 | -0.38 | -0.33 | lambda descreases | 419.78 | constant |
| **Birds** | B20 | 0.33 | 0.02 | 0.03 | 235.64 | 1.42 | 0.16 | 0.38 | 0 | 0 | -1.42 | -0.16 | -0.38 | lambda descreases | 235.29 | time-variable |
| **Birds** | B21 | 0.38 | 0.11 | 0.14 | 1520.64 | 0.18 | 0.34 | 0.76 | 0.14 | 0.19 | -0.04 | -0.2 | -0.62 | lambda descreases | 1516.5 | time-variable |
| **Birds** | B22 | 0.15 | 0.06 | 0.06 | 129.98 | -0.09 | 0.24 | 0.07 | 0.09 | 0.11 | 0.18 | -0.15 | 0.02 | unknown | 133.38 | constant |
| **Birds** | B23 | 0.56 | 0.05 | 0.07 | 117.59 | 1.03 | 0.45 | 0.63 | 0.03 | 0.04 | -1 | -0.42 | -0.6 | lambda descreases | 121.36 | constant |
| **Birds** | B24 | 0.63 | 0.09 | 0.1 | 40.84 | 2.44 | 0.98 | -1.52 | 0.01 | 0.01 | -2.43 | -0.97 | 1.53 | lambda descreases | 41.97 | constant |
| **Birds** | B25 | 0.53 | 0.02 | 0.04 | 60.54 | 3.11 | 0.18 | 0.75 | 0 | 0 | -3.11 | -0.18 | -0.75 | lambda descreases | 63.14 | constant |
| **Birds** | B26 | 0.85 | 0.09 | 0.1 | 47.34 | 0.58 | 0.93 | 0.76 | 0.1 | 0.12 | -0.48 | -0.83 | -0.66 | lambda descreases | 51.32 | constant |
| **Birds** | B27 | 0.17 | 0.13 | 0.46 | 44.53 | 0.64 | -0.28 | 0.96 | 0.1 | 0.34 | -0.54 | 0.37 | -0.86 | lambda descreases | 47.4 | constant |
| **Birds** | B28 | 0.19 | 0.16 | 0.21 | 240.72 | 0.86 | -0.11 | 0.54 | 0.06 | 0.09 | -0.8 | 0.18 | -0.47 | lambda descreases | 238.59 | time-variable |
| **Birds** | B29 | 0.07 | 0.03 | 0.1 | 77.49 | 0.36 | -0.06 | 0.25 | 0.01 | 0.03 | -0.35 | 0.07 | -0.24 | lambda descreases | 80.15 | constant |
| **Birds** | B30 | 0.39 | 0.07 | 0.07 | 46.45 | -0.49 | 0.75 | 0.02 | 0.14 | 0.14 | 0.64 | -0.61 | 0.12 | unknown | 49.94 | constant |
| **Birds** | B31 | 0.28 | 0.11 | 0.16 | 63.29 | 1.14 | -0.17 | 0.97 | 0.05 | 0.07 | -1.09 | 0.22 | -0.92 | lambda descreases | 65.87 | constant |
| **Birds** | B32 | 0.16 | 0.05 | 0.06 | 84.52 | 0.25 | 0.02 | 0.48 | 0.05 | 0.06 | -0.2 | 0.02 | -0.43 | lambda descreases | 87.63 | constant |
| **Squamata** | S1 | 0.15 | 0.02 | 0.04 | 532.96 | 0.73 | -0.02 | 0.37 | 0 | 0.01 | -0.73 | 0.02 | -0.37 | lambda descreases | 524.33 | time-variable |
| **Squamata** | S2 | 0.18 | 0.01 | 0.03 | 99.13 | 0.58 | 0.08 | 0.28 | 0 | 0.01 | -0.58 | -0.08 | -0.28 | lambda descreases | 102.57 | constant |
| **Squamata** | S3 | 0.17 | 0.06 | 0.25 | 120.49 | 0.48 | 0.02 | 0.4 | 0.03 | 0.14 | -0.45 | 0.02 | -0.37 | lambda descreases | 123.57 | constant |
| **Squamata** | S4 | 0.19 | 0.03 | 0.04 | 84.97 | 1.21 | -0.18 | 0.68 | 0 | 0 | -1.21 | 0.18 | -0.68 | lambda descreases | 85.49 | constant |

| | | | | | | | | | | | | | | | | |
|---|---|---|---|---|---|---|---|---|---|---|---|---|---|---|---|---|
| **Squamata** | S5 | 0.27 | 0.05 | 0.08 | 252.87 | 0.56 | 0.2 | 0.27 | 0.03 | 0.04 | -0.53 | -0.17 | -0.24 | lambda descreases | 255.92 | constant |
| **Squamata** | S6 | 0.12 | 0.01 | 0.02 | 115.61 | 0.06 | 0.15 | 0.06 | 0.01 | 0.02 | -0.05 | -0.14 | -0.05 | lambda descreases | 119.41 | constant |
| **Squamata** | S7 | 0.02 | 0.05 | 0.18 | 75.41 | 0.15 | -0.12 | 0.22 | 0.03 | 0.1 | -0.12 | 0.14 | -0.19 | lambda descreases | 77.12 | constant |
| **Squamata** | S8 | 0.07 | 0.05 | 0.1 | 216.36 | -0.24 | 0.19 | 0 | 0.13 | 0.26 | 0.36 | -0.06 | 0.13 | unknown | 215 | time-variable |
| **Squamata** | S9 | 0.12 | 0.02 | 0.04 | 1624.88 | 0.17 | 0.12 | 0.02 | 0.01 | 0.03 | -0.15 | -0.11 | -0.01 | lambda descreases | 1625.27 | constant |
| **Squamata** | S10 | 0.1 | 0.1 | 0.26 | 775.73 | 0.31 | 0.04 | 0.02 | 0.05 | 0.14 | -0.26 | 0.02 | 0.04 | lambda descreases | 764.42 | time-variable |
| **Squamata** | S11 | 0.17 | 0 | 0 | 79.97 | 1.88 | 0.12 | 0.17 | 0 | 0 | -1.88 | -0.12 | -0.17 | lambda descreases | 82.47 | constant |
| **Squamata** | S12 | 0.08 | 0 | 0.01 | 103.1 | 0.38 | 0.1 | -0.03 | 0 | 0 | -0.38 | -0.1 | 0.03 | lambda descreases | 103.28 | constant |
| **Squamata** | S13 | 0.25 | 0.04 | 0.08 | 268.07 | 0.5 | 0.12 | 0.49 | 0.02 | 0.06 | -0.48 | -0.09 | -0.46 | lambda descreases | 270.86 | constant |
| **Squamata** | S14 | 0.08 | 0.08 | 0.11 | 151.94 | 0.07 | 0.04 | 0.21 | 0.09 | 0.12 | 0.02 | 0.05 | -0.12 | unknown | 155.36 | constant |
| **Squamata** | S15 | 0.08 | 0.02 | 0.03 | 637.92 | 0.2 | 0.06 | 0.04 | 0.01 | 0.01 | -0.19 | -0.05 | -0.03 | lambda descreases | 636.97 | time-variable |
| **Squamata** | S16 | 0.11 | 0.04 | 0.1 | 126.84 | 0.13 | -0.06 | 0.58 | 0.06 | 0.14 | -0.07 | 0.12 | -0.53 | lambda descreases | 125.94 | time-variable |
| **Squamata** | S17 | 0.12 | 0 | 0.01 | 293.41 | 0.35 | 0.01 | 0.3 | 0 | 0 | -0.35 | -0.01 | -0.3 | lambda descreases | 293.58 | constant |
| **Squamata** | S18 | 0.07 | 0.02 | 0.04 | 1192.6 | 0.17 | 0.05 | 0.07 | 0.01 | 0.02 | -0.16 | -0.04 | -0.06 | lambda descreases | 1190.78 | time-variable |
| **Squamata** | S19 | 0.23 | 0.01 | 0.03 | 131.77 | 0.05 | 0.25 | 0.26 | 0.02 | 0.06 | -0.03 | -0.23 | -0.24 | lambda descreases | 135.48 | constant |
| **Squamata** | S20 | 0.15 | 0.01 | 0.01 | 117.44 | 0.45 | 0.07 | 0.23 | 0 | 0 | -0.44 | -0.07 | -0.23 | lambda descreases | 120.77 | constant |
| **Squamata** | S21 | 0.07 | 0.02 | 0.05 | 167.06 | 0.2 | 0.05 | 0.05 | 0 | 0.02 | -0.2 | -0.04 | -0.05 | lambda descreases | 169.92 | constant |
| **Squamata** | S22 | 0.09 | 0.01 | 0.02 | 577.74 | 0.07 | 0.09 | 0.07 | 0.01 | 0.02 | -0.06 | -0.08 | -0.06 | lambda descreases | 581.62 | constant |
| **Squamata** | S23 | 0.16 | 0.03 | 0.06 | 413.22 | 0.23 | 0.14 | 0.16 | 0.02 | 0.05 | -0.21 | -0.12 | -0.14 | lambda descreases | 416.97 | constant |
| **Squamata** | S24 | 0.22 | 0.01 | 0.06 | 304.42 | 0.18 | 0.22 | 0.21 | 0.01 | 0.07 | -0.17 | -0.21 | -0.2 | lambda descreases | 308.4 | constant |
| **Amphibia** | A1 | 0.08 | 0.04 | 0.04 | 908.19 | 0.1 | 0.09 | 0.05 | 0.03 | 0.03 | -0.07 | -0.05 | -0.02 | lambda descreases | 911.75 | constant |
| **Amphibia** | A2 | 0.09 | 0.05 | 0.05 | 1012.7 | 0.25 | 0.03 | 0.12 | 0.02 | 0.02 | -0.22 | -0.01 | -0.1 | lambda descreases | 1008.1 | time-variable |
| **Amphibia** | A3 | 0.06 | 0.03 | 0.03 | 716.31 | 0.14 | 0.03 | 0.09 | 0.02 | 0.02 | -0.12 | -0.02 | -0.07 | lambda descreases | 718.18 | constant |
| **Amphibia** | A4 | 0.12 | 0.02 | 0.03 | 1264.58 | 0.36 | 0.08 | 0.05 | 0.01 | 0.01 | -0.35 | -0.07 | -0.05 | lambda descreases | 1249.24 | time-variable |
| **Amphibia** | A5 | 0.09 | 0.01 | 0.03 | 498.94 | 0.23 | 0.06 | 0.08 | 0 | 0.01 | -0.23 | -0.06 | -0.07 | lambda descreases | 499.95 | constant |

| | | | | | | | | | | | | | | | | |
|---|---|---|---|---|---|---|---|---|---|---|---|---|---|---|---|---|
| **Amphibia** | A6 | 0.11 | 0.02 | 0.05 | 2243.42 | 0.16 | 0.1 | 0.07 | 0.02 | 0.04 | -0.14 | -0.09 | -0.05 | lambda descreases | 2244.92 | constant |
| **Amphibia** | A7 | 0.09 | 0.03 | 0.03 | 390.07 | 0.08 | 0.09 | 0.09 | 0.03 | 0.04 | -0.05 | -0.06 | -0.06 | lambda descreases | 394.05 | constant |
| **Amphibia** | A8 | 0.06 | 0.06 | 0.09 | 897.33 | 0.12 | 0.03 | 0.14 | 0.04 | 0.07 | -0.07 | 0.02 | -0.09 | lambda descreases | 899.1 | constant |
| **Amphibia** | A9 | 0.08 | 0.03 | 0.06 | 1142.79 | 0.14 | 0.07 | 0.06 | 0.02 | 0.04 | -0.11 | -0.05 | -0.03 | lambda descreases | 1144.8 | constant |
| **Amphibia** | A10 | 0.11 | 0.04 | 0.05 | 438.56 | 0.21 | 0.07 | 0.14 | 0.03 | 0.04 | -0.18 | -0.04 | -0.11 | lambda descreases | 441.79 | constant |
| **Amphibia** | A11 | 0.03 | 0.08 | 0.17 | 1009.67 | -0.07 | 0.05 | 0.08 | 0.13 | 0.26 | 0.2 | 0.08 | 0.05 | unknown | 998.45 | time-variable |
| **Amphibia** | A12 | 0.08 | 0.04 | 0.04 | 1443.98 | 0.16 | 0.06 | 0.06 | 0.02 | 0.03 | -0.14 | -0.03 | -0.03 | lambda descreases | 1441.95 | time-variable |
| **Amphibia** | A13 | 0.16 | 0.04 | 0.06 | 863.03 | 0.21 | 0.2 | -0.39 | 0.03 | 0.04 | -0.18 | -0.17 | 0.43 | lambda descreases | 848.46 | time-variable |
| **Amphibia** | A14 | 0.04 | 0.19 | 0.42 | 344.76 | 0.21 | -0.12 | 0.28 | 0.14 | 0.3 | -0.07 | 0.25 | -0.15 | lambda descreases | 342.17 | time-variable |
| **Amphibia** | A15 | 0.11 | 0.11 | 0.16 | 704.37 | 0.21 | 0.01 | 0.35 | 0.1 | 0.14 | -0.12 | 0.09 | -0.26 | lambda descreases | 704.01 | time-variable |
| **Amphibia** | A16 | 0.09 | 0.03 | 0.05 | 1235.92 | 0.27 | 0.04 | 0.09 | 0.01 | 0.02 | -0.26 | -0.04 | -0.08 | lambda descreases | 1226.49 | time-variable |

**Table S10**. Species-richness per clade and area. For each of the 150 phylogenetic clades considered in this study, we provide the number of species occurring in each of the 13 Neotropical ecoregions of the WWF biome classification (8). Clade numbers correspond with Table S1–5. Abbreviations: Amazonia = Amaz., Atlantic Forest = Atl.For., Bahama-Antilles = Baham., Caatinga, Central Andes = C.Andes, Cerrado, Chaco, Chocó, Guiana Shield, Mesoamerica = Mesoam., and the Northern Andes = N.Andes, temperate South America = TemSAm., and Elsewhere = Elsewh.

| Clade | Amaz. | Atl.For. | Baham. | Caatinga | C.Andes | Cerrado | Chaco | Chocó | Elsewh. | Galap. | Guiana | Mesoam. | N.Andes | TemSAm |
|---|---|---|---|---|---|---|---|---|---|---|---|---|---|---|
| **A1** | 46 | 1 | 1 | 0 | 3 | 0 | 0 | 3 | 0 | 0 | 48 | 1 | 26 | 0 |
| **A2** | 60 | 1 | 0 | 0 | 12 | 4 | 0 | 26 | 0 | 0 | 8 | 17 | 59 | 0 |
| **A3** | 5 | 13 | 0 | 1 | 25 | 1 | 1 | 6 | 0 | 0 | 18 | 2 | 31 | 1 |
| **A4** | 2 | 5 | 0 | 2 | 0 | 0 | 0 | 1 | 0 | 0 | 2 | 7 | 0 | 0 |
| **A5** | 4 | 0 | 143 | 0 | 0 | 0 | 0 | 5 | 2 | 0 | 1 | 63 | 3 | 0 |
| **A6** | 43 | 4 | 2 | 1 | 77 | 4 | 1 | 26 | 0 | 0 | 25 | 28 | 155 | 1 |
| **A7** | 12 | 15 | 0 | 4 | 3 | 9 | 4 | 3 | 0 | 0 | 6 | 11 | 6 | 2 |
| **A8** | 20 | 44 | 1 | 4 | 12 | 10 | 5 | 8 | 0 | 0 | 28 | 4 | 28 | 4 |
| **A9** | 64 | 41 | 0 | 5 | 3 | 32 | 11 | 11 | 0 | 0 | 28 | 9 | 31 | 5 |
| **A10** | 26 | 9 | 8 | 4 | 5 | 5 | 2 | 3 | 1 | 0 | 21 | 2 | 8 | 2 |
| **A11** | 2 | 37 | 0 | 4 | 33 | 4 | 8 | 0 | 0 | 0 | 1 | 0 | 2 | 46 |
| **A12** | 53 | 64 | 3 | 14 | 12 | 49 | 22 | 18 | 0 | 0 | 38 | 11 | 22 | 17 |
| **A13** | 23 | 2 | 0 | 0 | 19 | 0 | 0 | 22 | 0 | 0 | 18 | 15 | 59 | 0 |
| **A14** | 6 | 1 | 0 | 0 | 7 | 0 | 0 | 1 | 0 | 0 | 12 | 5 | 27 | 0 |
| **A15** | 19 | 14 | 0 | 5 | 13 | 13 | 5 | 5 | 25 | 0 | 10 | 30 | 8 | 6 |
| **A16** | 4 | 0 | 0 | 0 | 1 | 0 | 0 | 2 | 8 | 0 | 0 | 149 | 6 | 0 |

| | | | | | | | | | | | | | | |
|---|---|---|---|---|---|---|---|---|---|---|---|---|---|---|
| **B1** | 15 | 5 | 0 | 1 | 8 | 7 | 5 | 8 | 3 | 0 | 10 | 9 | 11 | 2 |
| **B2** | 61 | 29 | 6 | 20 | 90 | 24 | 12 | 59 | 19 | 0 | 53 | 70 | 127 | 16 |
| **B3** | 4 | 3 | 0 | 2 | 4 | 4 | 4 | 2 | 0 | 0 | 3 | 2 | 3 | 5 |
| **B4** | 47 | 24 | 11 | 19 | 32 | 33 | 16 | 27 | 16 | 0 | 36 | 19 | 31 | 13 |
| **B5** | 11 | 8 | 0 | 1 | 19 | 5 | 3 | 4 | 0 | 0 | 6 | 4 | 25 | 11 |
| **B6** | 3 | 2 | 0 | 2 | 3 | 2 | 1 | 2 | 0 | 0 | 0 | 1 | 2 | 2 |
| **B7** | 5 | 3 | 0 | 2 | 3 | 0 | 1 | 2 | 0 | 0 | 3 | 2 | 4 | 1 |
| **B8** | 112 | 32 | 0 | 20 | 63 | 34 | 9 | 35 | 0 | 0 | 55 | 26 | 68 | 4 |
| **B9** | 86 | 53 | 0 | 25 | 140 | 58 | 52 | 45 | 3 | 0 | 66 | 42 | 96 | 76 |
| **B10** | 153 | 104 | 1 | 72 | 169 | 107 | 76 | 101 | 37 | 0 | 131 | 98 | 148 | 72 |
| **B11** | 46 | 16 | 0 | 7 | 31 | 18 | 1 | 16 | 0 | 0 | 33 | 13 | 28 | 3 |
| **B12** | 4 | 0 | 1 | 0 | 4 | 1 | 0 | 3 | 13 | 0 | 4 | 16 | 7 | 0 |
| **B13** | 8 | 5 | 0 | 1 | 5 | 4 | 3 | 2 | 12 | 0 | 5 | 18 | 7 | 2 |
| **B14** | 9 | 6 | 4 | 6 | 13 | 5 | 2 | 10 | 10 | 0 | 10 | 8 | 19 | 4 |
| **B15** | 1 | 1 | 0 | 0 | 4 | 0 | 0 | 4 | 9 | 0 | 2 | 13 | 6 | 0 |
| **B16** | 15 | 8 | 0 | 4 | 17 | 8 | 5 | 21 | 17 | 0 | 14 | 46 | 23 | 4 |
| **B17** | 3 | 3 | 1 | 3 | 3 | 2 | 2 | 2 | 2 | 0 | 4 | 5 | 4 | 2 |
| **B18** | 7 | 3 | 0 | 2 | 19 | 4 | 6 | 10 | 49 | 0 | 6 | 51 | 17 | 4 |
| **B19** | 26 | 18 | 10 | 8 | 25 | 17 | 19 | 19 | 25 | 0 | 23 | 38 | 29 | 17 |
| **B20** | 8 | 5 | 1 | 2 | 9 | 5 | 4 | 12 | 19 | 0 | 12 | 32 | 15 | 3 |
| **B21** | 95 | 68 | 15 | 38 | 150 | 63 | 49 | 84 | 11 | 13 | 81 | 63 | 150 | 54 |
| **B22** | 6 | 4 | 2 | 2 | 5 | 4 | 1 | 6 | 3 | 0 | 6 | 11 | 8 | 0 |
| **B23** | 12 | 7 | 1 | 5 | 11 | 11 | 6 | 5 | 5 | 0 | 8 | 6 | 10 | 4 |
| **B24** | 2 | 1 | 0 | 1 | 5 | 2 | 2 | 3 | 3 | 0 | 2 | 3 | 5 | 2 |
| **B25** | 1 | 2 | 1 | 0 | 1 | 2 | 2 | 1 | 6 | 0 | 0 | 5 | 2 | 2 |

| | | | | | | | | | | | | | | |
|---|---|---|---|---|---|---|---|---|---|---|---|---|---|---|
| **B26** | 4 | 3 | 0 | 2 | 7 | 4 | 3 | 3 | 0 | 0 | 3 | 1 | 7 | 4 |
| **B27** | 5 | 3 | 0 | 1 | 2 | 2 | 1 | 0 | 1 | 0 | 3 | 0 | 1 | 1 |
| **B28** | 21 | 7 | 0 | 3 | 16 | 9 | 2 | 15 | 0 | 0 | 13 | 10 | 25 | 1 |
| **B29** | 10 | 1 | 0 | 2 | 3 | 3 | 1 | 1 | 0 | 0 | 6 | 1 | 4 | 0 |
| **B30** | 4 | 4 | 0 | 2 | 4 | 3 | 3 | 5 | 1 | 0 | 6 | 5 | 5 | 3 |
| **B31** | 4 | 4 | 0 | 3 | 7 | 4 | 3 | 5 | 2 | 0 | 5 | 6 | 6 | 3 |
| **B32** | 7 | 5 | 0 | 5 | 8 | 6 | 4 | 7 | 3 | 0 | 5 | 5 | 7 | 6 |
| **M1** | 14 | 10 | 2 | 5 | 13 | 11 | 13 | 9 | 3 | 0 | 13 | 10 | 11 | 9 |
| **M2** | 72 | 40 | 20 | 20 | 78 | 29 | 15 | 82 | 8 | 0 | 89 | 77 | 98 | 2 |
| **M3** | 12 | 6 | 2 | 3 | 7 | 7 | 11 | 7 | 2 | 0 | 15 | 12 | 8 | 3 |
| **M4** | 61 | 14 | 0 | 6 | 14 | 12 | 5 | 12 | 8 | 0 | 17 | 10 | 25 | 0 |
| **M5** | 21 | 17 | 1 | 5 | 19 | 13 | 10 | 8 | 3 | 0 | 17 | 8 | 15 | 6 |
| **M6** | 57 | 51 | 2 | 8 | 106 | 48 | 49 | 22 | 10 | 4 | 33 | 29 | 50 | 65 |
| **M7** | 0 | 0 | 0 | 0 | 0 | 0 | 0 | 3 | 3 | 0 | 1 | 10 | 3 | 0 |
| **M8** | 0 | 0 | 0 | 0 | 0 | 0 | 0 | 0 | 1 | 0 | 0 | 10 | 1 | 0 |
| **M9** | 13 | 5 | 0 | 4 | 6 | 5 | 0 | 12 | 0 | 0 | 16 | 12 | 9 | 0 |
| **M10** | 4 | 2 | 0 | 1 | 5 | 4 | 4 | 3 | 2 | 0 | 3 | 3 | 5 | 2 |
| **M11** | 0 | 0 | 1 | 0 | 2 | 0 | 1 | 0 | 6 | 0 | 1 | 6 | 1 | 2 |
| **M12** | 35 | 19 | 5 | 9 | 28 | 12 | 23 | 11 | 0 | 0 | 19 | 5 | 19 | 37 |
| **P1** | 3 | 3 | 3 | 3 | 3 | 0 | 0 | 3 | 0 | 0 | 4 | 13 | 3 | 0 |
| **P2** | 0 | 2 | 0 | 0 | 0 | 0 | 0 | 2 | 0 | 0 | 2 | 34 | 6 | 0 |
| **P3** | 5 | 1 | 3 | 1 | 5 | 2 | 0 | 5 | 0 | 0 | 4 | 8 | 7 | 0 |
| **P4** | 4 | 2 | 4 | 1 | 4 | 1 | 0 | 4 | 0 | 0 | 4 | 6 | 4 | 0 |
| **P5** | 6 | 1 | 0 | 1 | 1 | 1 | 0 | 2 | 0 | 0 | 3 | 3 | 8 | 0 |
| **P6** | 14 | 0 | 0 | 0 | 5 | 0 | 0 | 0 | 0 | 0 | 0 | 0 | 11 | 0 |

| | | | | | | | | | | | | | | |
|---|---|---|---|---|---|---|---|---|---|---|---|---|---|---|
| **P7** | 21 | 1 | 2 | 0 | 6 | 3 | 0 | 4 | 0 | 0 | 5 | 8 | 10 | 0 |
| **P8** | 0 | 0 | 1 | 0 | 7 | 0 | 0 | 1 | 0 | 0 | 0 | 0 | 7 | 2 |
| **P9** | 6 | 1 | 2 | 1 | 3 | 2 | 0 | 4 | 0 | 0 | 7 | 6 | 4 | 0 |
| **P10** | 19 | 2 | 1 | 1 | 13 | 2 | 0 | 11 | 0 | 0 | 11 | 11 | 15 | 0 |
| **P11** | 19 | 11 | 4 | 9 | 7 | 11 | 4 | 8 | 0 | 0 | 14 | 9 | 11 | 0 |
| **P12** | 7 | 2 | 0 | 3 | 7 | 2 | 4 | 3 | 0 | 0 | 5 | 5 | 4 | 1 |
| **P13** | 0 | 1 | 20 | 1 | 0 | 1 | 1 | 3 | 20 | 0 | 4 | 19 | 1 | 0 |
| **P14** | 5 | 0 | 1 | 1 | 2 | 0 | 0 | 6 | 0 | 0 | 3 | 4 | 7 | 0 |
| **P15** | 11 | 2 | 0 | 1 | 1 | 0 | 0 | 7 | 0 | 0 | 7 | 5 | 8 | 0 |
| **P16** | 18 | 1 | 0 | 1 | 3 | 5 | 0 | 1 | 0 | 0 | 15 | 2 | 2 | 0 |
| **P17** | 12 | 2 | 0 | 1 | 7 | 0 | 0 | 4 | 0 | 0 | 6 | 3 | 11 | 0 |
| **P18** | 1 | 0 | 0 | 0 | 6 | 0 | 0 | 0 | 0 | 0 | 0 | 1 | 1 | 0 |
| **P19** | 9 | 0 | 0 | 0 | 16 | 0 | 0 | 2 | 0 | 0 | 5 | 0 | 16 | 0 |
| **P20** | 2 | 1 | 0 | 2 | 6 | 3 | 2 | 5 | 3 | 0 | 4 | 15 | 7 | 0 |
| **P21** | 42 | 20 | 2 | 17 | 20 | 28 | 0 | 14 | 0 | 0 | 34 | 25 | 29 | 0 |
| **P22** | 0 | 1 | 1 | 0 | 10 | 1 | 0 | 2 | 0 | 0 | 1 | 6 | 12 | 0 |
| **P23** | 7 | 0 | 1 | 0 | 2 | 2 | 0 | 4 | 0 | 0 | 8 | 3 | 5 | 0 |
| **P24** | 5 | 13 | 0 | 2 | 108 | 9 | 0 | 15 | 0 | 0 | 4 | 46 | 81 | 4 |
| **P25** | 0 | 0 | 0 | 0 | 0 | 0 | 0 | 0 | 0 | 0 | 0 | 24 | 0 | 0 |
| **P26** | 3 | 0 | 9 | 0 | 1 | 0 | 0 | 7 | 0 | 0 | 1 | 30 | 2 | 0 |
| **P27** | 0 | 15 | 0 | 1 | 1 | 4 | 5 | 0 | 0 | 0 | 0 | 0 | 0 | 7 |
| **P28** | 3 | 0 | 0 | 0 | 24 | 1 | 0 | 0 | 1 | 0 | 0 | 5 | 14 | 2 |
| **P29** | 7 | 4 | 1 | 2 | 3 | 3 | 1 | 2 | 0 | 0 | 3 | 3 | 3 | 0 |
| **P30** | 0 | 0 | 6 | 0 | 0 | 0 | 0 | 1 | 4 | 0 | 1 | 8 | 0 | 0 |
| **P31** | 4 | 2 | 58 | 0 | 1 | 0 | 2 | 3 | 0 | 0 | 1 | 10 | 0 | 1 |

| | | | | | | | | | | | | | | |
|---|---|---|---|---|---|---|---|---|---|---|---|---|---|---|
| **P32** | 1 | 0 | 19 | 1 | 0 | 1 | 1 | 1 | 0 | 0 | 1 | 0 | 0 | 0 |
| **P33** | 0 | 0 | 3 | 0 | 0 | 0 | 0 | 0 | 7 | 0 | 0 | 9 | 0 | 0 |
| **P34** | 13 | 0 | 0 | 0 | 7 | 1 | 0 | 11 | 0 | 0 | 7 | 7 | 22 | 0 |
| **P35** | 4 | 0 | 3 | 0 | 8 | 0 | 0 | 11 | 0 | 0 | 1 | 102 | 10 | 0 |
| **P36** | 40 | 60 | 2 | 17 | 10 | 55 | 7 | 9 | 3 | 0 | 21 | 6 | 11 | 6 |
| **P37** | 88 | 23 | 8 | 10 | 24 | 22 | 2 | 32 | 1 | 0 | 41 | 40 | 43 | 0 |
| **P38** | 28 | 4 | 5 | 2 | 20 | 1 | 0 | 27 | 0 | 0 | 20 | 44 | 41 | 0 |
| **P39** | 8 | 1 | 2 | 1 | 8 | 1 | 0 | 8 | 0 | 0 | 9 | 10 | 12 | 0 |
| **P40** | 31 | 64 | 8 | 61 | 28 | 119 | 21 | 11 | 33 | 0 | 27 | 72 | 20 | 14 |
| **P41** | 16 | 163 | 31 | 17 | 60 | 12 | 0 | 60 | 0 | 0 | 49 | 315 | 250 | 3 |
| **P42** | 88 | 109 | 43 | 16 | 141 | 33 | 8 | 100 | 69 | 0 | 81 | 272 | 320 | 2 |
| **P43** | 39 | 96 | 42 | 48 | 37 | 84 | 67 | 22 | 18 | 8 | 51 | 54 | 36 | 34 |
| **P44** | 112 | 20 | 4 | 16 | 82 | 27 | 5 | 61 | 0 | 0 | 70 | 52 | 87 | 0 |
| **P45** | 92 | 127 | 53 | 12 | 113 | 39 | 8 | 115 | 2 | 0 | 70 | 161 | 227 | 5 |
| **P46** | 70 | 16 | 9 | 9 | 25 | 17 | 4 | 19 | 6 | 0 | 53 | 18 | 36 | 0 |
| **P47** | 1 | 2 | 5 | 2 | 1 | 0 | 0 | 0 | 1 | 0 | 0 | 0 | 0 | 12 |
| **P48** | 3 | 0 | 10 | 0 | 6 | 2 | 0 | 2 | 1 | 0 | 1 | 30 | 9 | 7 |
| **P49** | 0 | 11 | 1 | 1 | 15 | 1 | 11 | 0 | 4 | 0 | 0 | 5 | 3 | 25 |
| **P50** | 1 | 0 | 0 | 0 | 40 | 0 | 0 | 0 | 0 | 1 | 0 | 0 | 0 | 28 |
| **P51** | 21 | 18 | 17 | 7 | 65 | 8 | 13 | 23 | 43 | 3 | 16 | 84 | 59 | 9 |
| **P52** | 77 | 56 | 19 | 30 | 187 | 48 | 38 | 44 | 192 | 3 | 41 | 98 | 93 | 44 |
| **P53** | 71 | 3 | 2 | 1 | 3 | 2 | 1 | 30 | 15 | 0 | 64 | 13 | 26 | 0 |
| **P54** | 2 | 3 | 2 | 13 | 4 | 8 | 5 | 1 | 5 | 0 | 2 | 1 | 4 | 9 |
| **P55** | 54 | 29 | 0 | 10 | 9 | 26 | 2 | 14 | 0 | 0 | 21 | 47 | 14 | 0 |
| **P56** | 35 | 114 | 20 | 60 | 19 | 79 | 6 | 13 | 1 | 0 | 34 | 12 | 13 | 6 |

| | | | | | | | | | | | | | | |
|---|---|---|---|---|---|---|---|---|---|---|---|---|---|---|
| **P57** | 10 | 34 | 4 | 17 | 11 | 19 | 7 | 3 | 105 | 0 | 2 | 8 | 5 | 12 |
| **P58** | 26 | 40 | 40 | 30 | 33 | 40 | 19 | 25 | 82 | 0 | 37 | 103 | 41 | 13 |
| **P59** | 21 | 6 | 0 | 2 | 3 | 3 | 0 | 2 | 2 | 0 | 17 | 2 | 3 | 0 |
| **P60** | 0 | 1 | 0 | 0 | 5 | 0 | 1 | 2 | 0 | 0 | 1 | 2 | 8 | 8 |
| **P61** | 8 | 7 | 0 | 4 | 4 | 6 | 2 | 3 | 0 | 0 | 6 | 2 | 3 | 0 |
| **P62** | 5 | 3 | 0 | 3 | 2 | 3 | 0 | 1 | 0 | 0 | 5 | 2 | 1 | 0 |
| **P63** | 0 | 0 | 1 | 0 | 1 | 0 | 0 | 1 | 4 | 0 | 0 | 7 | 2 | 0 |
| **P64** | 0 | 4 | 3 | 2 | 1 | 5 | 2 | 1 | 0 | 0 | 0 | 2 | 0 | 0 |
| **P65** | 1 | 7 | 5 | 1 | 3 | 3 | 1 | 0 | 0 | 0 | 2 | 16 | 6 | 1 |
| **P66** | 9 | 26 | 0 | 8 | 13 | 20 | 5 | 4 | 5 | 0 | 5 | 11 | 11 | 2 |
| **S1** | 24 | 31 | 1 | 9 | 4 | 29 | 13 | 6 | 0 | 0 | 16 | 4 | 9 | 13 |
| **S2** | 7 | 7 | 0 | 1 | 3 | 5 | 0 | 5 | 3 | 0 | 6 | 5 | 8 | 0 |
| **S3** | 6 | 7 | 0 | 2 | 3 | 3 | 3 | 2 | 1 | 0 | 4 | 3 | 5 | 4 |
| **S4** | 1 | 1 | 0 | 1 | 0 | 0 | 0 | 2 | 0 | 0 | 1 | 13 | 2 | 0 |
| **S5** | 7 | 13 | 2 | 2 | 3 | 4 | 3 | 6 | 0 | 0 | 5 | 14 | 5 | 2 |
| **S6** | 6 | 4 | 9 | 2 | 3 | 4 | 4 | 3 | 0 | 0 | 6 | 2 | 4 | 0 |
| **S7** | 1 | 0 | 6 | 0 | 0 | 0 | 0 | 2 | 0 | 0 | 1 | 1 | 1 | 0 |
| **S8** | 1 | 1 | 27 | 0 | 0 | 1 | 1 | 1 | 0 | 0 | 2 | 0 | 0 | 0 |
| **S9** | 9 | 3 | 129 | 0 | 3 | 1 | 0 | 20 | 1 | 0 | 12 | 48 | 25 | 0 |
| **S10** | 0 | 2 | 0 | 0 | 25 | 0 | 8 | 0 | 0 | 0 | 0 | 0 | 0 | 90 |
| **S11** | 1 | 2 | 0 | 0 | 0 | 0 | 2 | 0 | 0 | 0 | 0 | 0 | 0 | 7 |
| **S12** | 5 | 0 | 0 | 1 | 1 | 2 | 1 | 3 | 0 | 0 | 1 | 2 | 8 | 0 |
| **S13** | 0 | 0 | 0 | 0 | 0 | 0 | 0 | 0 | 25 | 0 | 0 | 29 | 0 | 0 |
| **S14** | 0 | 0 | 8 | 0 | 0 | 0 | 0 | 0 | 1 | 0 | 0 | 14 | 0 | 0 |
| **S15** | 11 | 7 | 0 | 8 | 35 | 11 | 4 | 16 | 0 | 0 | 7 | 0 | 15 | 3 |

| | | | | | | | | | | | | | | |
|---|---|---|---|---|---|---|---|---|---|---|---|---|---|---|
| **S16** | 0 | 0 | 0 | 0 | 0 | 0 | 0 | 0 | 2 | 0 | 0 | 17 | 0 | 0 |
| **S17** | 8 | 6 | 12 | 4 | 1 | 12 | 8 | 1 | 0 | 0 | 2 | 0 | 2 | 5 |
| **S18** | 41 | 25 | 21 | 14 | 18 | 27 | 11 | 10 | 10 | 0 | 33 | 12 | 25 | 7 |
| **S19** | 6 | 5 | 2 | 5 | 1 | 3 | 3 | 1 | 0 | 0 | 5 | 1 | 2 | 1 |
| **S20** | 0 | 0 | 0 | 0 | 0 | 0 | 0 | 0 | 0 | 0 | 0 | 16 | 0 | 0 |
| **S21** | 0 | 0 | 1 | 0 | 1 | 0 | 2 | 1 | 6 | 0 | 0 | 9 | 1 | 5 |
| **S22** | 9 | 2 | 39 | 2 | 2 | 1 | 0 | 4 | 0 | 0 | 22 | 4 | 8 | 0 |
| **S23** | 8 | 12 | 28 | 4 | 3 | 11 | 13 | 7 | 0 | 0 | 10 | 3 | 5 | 13 |
| **S24** | 12 | 11 | 1 | 1 | 2 | 4 | 3 | 8 | 2 | 0 | 13 | 22 | 10 | 2 |

**Table S11.** For each of the 150 phylogenetic clades considered in this study, we provide the proportion of species occurring in the 5 main biogeographic clusters identified here based on K-means clustering algorithms: cluster 1 (including the Amazonia, Central Andes, Chocó, Guiana Shield, Mesoamerica, and Northern Andes), cluster 2 (Atlantic Forest, Caatinga, Cerrado, Chaco, and temperate South America), cluster 3 (Bahama-Antilles), cluster 4 ("elsewhere" region), or cluster 5 (Galapagos). Clades were assigned to a given cluster only if > 60% of the species in the clade are distributed in the cluster, otherwise clades are classified as 'mixed' (main cluster 0). Clade numbers correspond with Table S1–5. Cluster numbers correspond with Figure 6.

| Clade | cluster-1 | cluster-2 | cluster-3 | cluster-4 | cluster-5 | Assignation to cluster |
|---|---|---|---|---|---|---|
| **A1** | 0.98 | 0.01 | 0 | 0 | 0.01 | 1 |
| **A10** | 0.68 | 0.23 | 0.01 | 0 | 0.08 | 1 |
| **A11** | 0.28 | 0.72 | 0 | 0 | 0 | 2 |
| **A12** | 0.48 | 0.51 | 0 | 0 | 0.01 | 0 |
| **A13** | 0.99 | 0.01 | 0 | 0 | 0 | 1 |
| **A14** | 0.98 | 0.02 | 0 | 0 | 0 | 1 |
| **A15** | 0.56 | 0.28 | 0.16 | 0 | 0 | 0 |
| **A16** | 0.95 | 0 | 0.05 | 0 | 0 | 1 |
| **A2** | 0.97 | 0.03 | 0 | 0 | 0 | 1 |
| **A3** | 0.84 | 0.16 | 0 | 0 | 0 | 1 |
| **A4** | 0.63 | 0.37 | 0 | 0 | 0 | 1 |
| **A5** | 0.34 | 0 | 0.01 | 0 | 0.65 | 5 |
| **A6** | 0.96 | 0.03 | 0 | 0 | 0.01 | 1 |
| **A7** | 0.55 | 0.45 | 0 | 0 | 0 | 0 |
| **A8** | 0.6 | 0.4 | 0 | 0 | 0.01 | 1 |

| | | | | | | |
|---|---|---|---|---|---|---|
| **A9** | 0.61 | 0.39 | 0 | 0 | 0 | 1 |
| **B1** | 0.73 | 0.24 | 0.04 | 0 | 0 | 1 |
| **B10** | 0.63 | 0.34 | 0.03 | 0 | 0 | 1 |
| **B11** | 0.79 | 0.21 | 0 | 0 | 0 | 1 |
| **B12** | 0.72 | 0.02 | 0.25 | 0 | 0.02 | 1 |
| **B13** | 0.62 | 0.21 | 0.17 | 0 | 0 | 1 |
| **B14** | 0.65 | 0.22 | 0.09 | 0 | 0.04 | 1 |
| **B15** | 0.75 | 0.03 | 0.22 | 0 | 0 | 1 |
| **B16** | 0.75 | 0.16 | 0.09 | 0 | 0 | 1 |
| **B17** | 0.58 | 0.33 | 0.06 | 0 | 0.03 | 0 |
| **B18** | 0.62 | 0.11 | 0.28 | 0 | 0 | 1 |
| **B19** | 0.58 | 0.29 | 0.09 | 0 | 0.04 | 0 |
| **B2** | 0.78 | 0.17 | 0.03 | 0 | 0.01 | 1 |
| **B20** | 0.69 | 0.15 | 0.15 | 0 | 0.01 | 1 |
| **B21** | 0.67 | 0.29 | 0.01 | 0.01 | 0.02 | 1 |
| **B22** | 0.72 | 0.19 | 0.05 | 0 | 0.03 | 1 |
| **B23** | 0.57 | 0.36 | 0.05 | 0 | 0.01 | 0 |
| **B24** | 0.65 | 0.26 | 0.1 | 0 | 0 | 1 |
| **B25** | 0.4 | 0.32 | 0.24 | 0 | 0.04 | 0 |
| **B26** | 0.61 | 0.39 | 0 | 0 | 0 | 1 |
| **B27** | 0.55 | 0.4 | 0.05 | 0 | 0 | 0 |
| **B28** | 0.82 | 0.18 | 0 | 0 | 0 | 1 |
| **B29** | 0.78 | 0.22 | 0 | 0 | 0 | 1 |
| **B3** | 0.5 | 0.5 | 0 | 0 | 0 | 0 |
| **B30** | 0.64 | 0.33 | 0.02 | 0 | 0 | 1 |

| | | | | | | |
|---|---|---|---|---|---|---|
| **B31** | 0.63 | 0.33 | 0.04 | 0 | 0 | 1 |
| **B32** | 0.57 | 0.38 | 0.04 | 0 | 0 | 0 |
| **B4** | 0.59 | 0.32 | 0.05 | 0 | 0.03 | 0 |
| **B5** | 0.71 | 0.29 | 0 | 0 | 0 | 1 |
| **B6** | 0.55 | 0.45 | 0 | 0 | 0 | 0 |
| **B7** | 0.73 | 0.27 | 0 | 0 | 0 | 1 |
| **B8** | 0.78 | 0.22 | 0 | 0 | 0 | 1 |
| **B9** | 0.64 | 0.36 | 0 | 0 | 0 | 1 |
| **M1** | 0.57 | 0.39 | 0.02 | 0 | 0.02 | 0 |
| **M10** | 0.61 | 0.34 | 0.05 | 0 | 0 | 1 |
| **M11** | 0.5 | 0.15 | 0.3 | 0 | 0.05 | 0 |
| **M12** | 0.53 | 0.45 | 0 | 0 | 0.02 | 0 |
| **M2** | 0.79 | 0.17 | 0.01 | 0 | 0.03 | 1 |
| **M3** | 0.64 | 0.32 | 0.02 | 0 | 0.02 | 1 |
| **M4** | 0.76 | 0.2 | 0.04 | 0 | 0 | 1 |
| **M5** | 0.62 | 0.36 | 0.02 | 0 | 0.01 | 1 |
| **M6** | 0.56 | 0.41 | 0.02 | 0.01 | 0 | 0 |
| **M7** | 0.85 | 0 | 0.15 | 0 | 0 | 1 |
| **M8** | 0.92 | 0 | 0.08 | 0 | 0 | 1 |
| **M9** | 0.83 | 0.17 | 0 | 0 | 0 | 1 |
| **P1** | 0.76 | 0.16 | 0 | 0 | 0.08 | 1 |
| **P10** | 0.93 | 0.06 | 0 | 0 | 0.01 | 1 |
| **P11** | 0.64 | 0.33 | 0 | 0 | 0.04 | 1 |
| **P12** | 0.72 | 0.28 | 0 | 0 | 0 | 1 |
| **P13** | 0.38 | 0.06 | 0.28 | 0 | 0.28 | 0 |

| | | | | | | |
|---|---|---|---|---|---|---|
| **P14** | 0.93 | 0.03 | 0 | 0 | 0.03 | 1 |
| **P15** | 0.93 | 0.07 | 0 | 0 | 0 | 1 |
| **P16** | 0.85 | 0.15 | 0 | 0 | 0 | 1 |
| **P17** | 0.93 | 0.07 | 0 | 0 | 0 | 1 |
| **P18** | 1 | 0 | 0 | 0 | 0 | 1 |
| **P19** | 1 | 0 | 0 | 0 | 0 | 1 |
| **P2** | 0.96 | 0.04 | 0 | 0 | 0 | 1 |
| **P20** | 0.78 | 0.16 | 0.06 | 0 | 0 | 1 |
| **P21** | 0.71 | 0.28 | 0 | 0 | 0.01 | 1 |
| **P22** | 0.91 | 0.06 | 0 | 0 | 0.03 | 1 |
| **P23** | 0.91 | 0.06 | 0 | 0 | 0.03 | 1 |
| **P24** | 0.9 | 0.1 | 0 | 0 | 0 | 1 |
| **P25** | 1 | 0 | 0 | 0 | 0 | 1 |
| **P26** | 0.83 | 0 | 0 | 0 | 0.17 | 1 |
| **P27** | 0.03 | 0.97 | 0 | 0 | 0 | 2 |
| **P28** | 0.92 | 0.06 | 0.02 | 0 | 0 | 1 |
| **P29** | 0.66 | 0.31 | 0 | 0 | 0.03 | 1 |
| **P3** | 0.83 | 0.1 | 0 | 0 | 0.07 | 1 |
| **P30** | 0.5 | 0 | 0.2 | 0 | 0.3 | 0 |
| **P31** | 0.23 | 0.06 | 0 | 0 | 0.71 | 5 |
| **P32** | 0.12 | 0.12 | 0 | 0 | 0.76 | 5 |
| **P33** | 0.47 | 0 | 0.37 | 0 | 0.16 | 0 |
| **P34** | 0.99 | 0.01 | 0 | 0 | 0 | 1 |
| **P35** | 0.98 | 0 | 0 | 0 | 0.02 | 1 |
| **P36** | 0.39 | 0.59 | 0.01 | 0 | 0.01 | 0 |

| | | | | | | |
|---|---|---|---|---|---|---|
| **P37** | 0.8 | 0.17 | 0 | 0 | 0.02 | 1 |
| **P38** | 0.94 | 0.04 | 0 | 0 | 0.03 | 1 |
| **P39** | 0.92 | 0.05 | 0 | 0 | 0.03 | 1 |
| **P4** | 0.76 | 0.12 | 0 | 0 | 0.12 | 1 |
| **P40** | 0.37 | 0.55 | 0.06 | 0 | 0.02 | 0 |
| **P41** | 0.77 | 0.2 | 0 | 0 | 0.03 | 1 |
| **P42** | 0.78 | 0.13 | 0.05 | 0 | 0.03 | 1 |
| **P43** | 0.38 | 0.52 | 0.03 | 0.01 | 0.07 | 0 |
| **P44** | 0.87 | 0.13 | 0 | 0 | 0.01 | 1 |
| **P45** | 0.76 | 0.19 | 0 | 0 | 0.05 | 1 |
| **P46** | 0.78 | 0.16 | 0.02 | 0 | 0.03 | 1 |
| **P47** | 0.08 | 0.67 | 0.04 | 0 | 0.21 | 2 |
| **P48** | 0.72 | 0.13 | 0.01 | 0 | 0.14 | 1 |
| **P49** | 0.3 | 0.64 | 0.05 | 0 | 0.01 | 2 |
| **P5** | 0.88 | 0.12 | 0 | 0 | 0 | 1 |
| **P50** | 0.59 | 0.4 | 0 | 0.01 | 0 | 0 |
| **P51** | 0.69 | 0.14 | 0.11 | 0.01 | 0.04 | 1 |
| **P52** | 0.56 | 0.22 | 0.2 | 0 | 0.02 | 0 |
| **P53** | 0.9 | 0.03 | 0.06 | 0 | 0.01 | 1 |
| **P54** | 0.24 | 0.64 | 0.08 | 0 | 0.03 | 2 |
| **P55** | 0.7 | 0.3 | 0 | 0 | 0 | 1 |
| **P56** | 0.31 | 0.64 | 0 | 0 | 0.05 | 2 |
| **P57** | 0.16 | 0.38 | 0.44 | 0 | 0.02 | 0 |
| **P58** | 0.5 | 0.27 | 0.16 | 0 | 0.08 | 0 |
| **P59** | 0.79 | 0.18 | 0.03 | 0 | 0 | 1 |

| | | | | | | |
|---|---|---|---|---|---|---|
| **P6** | 1 | 0 | 0 | 0 | 0 | 1 |
| **P60** | 0.64 | 0.36 | 0 | 0 | 0 | 1 |
| **P61** | 0.58 | 0.42 | 0 | 0 | 0 | 0 |
| **P62** | 0.64 | 0.36 | 0 | 0 | 0 | 1 |
| **P63** | 0.69 | 0 | 0.25 | 0 | 0.06 | 1 |
| **P64** | 0.2 | 0.65 | 0 | 0 | 0.15 | 2 |
| **P65** | 0.61 | 0.28 | 0 | 0 | 0.11 | 1 |
| **P66** | 0.45 | 0.51 | 0.04 | 0 | 0 | 0 |
| **P7** | 0.9 | 0.07 | 0 | 0 | 0.03 | 1 |
| **P8** | 0.83 | 0.11 | 0 | 0 | 0.06 | 1 |
| **P9** | 0.83 | 0.11 | 0 | 0 | 0.06 | 1 |
| **S1** | 0.4 | 0.6 | 0 | 0 | 0.01 | 2 |
| **S10** | 0.2 | 0.8 | 0 | 0 | 0 | 2 |
| **S11** | 0.08 | 0.92 | 0 | 0 | 0 | 2 |
| **S12** | 0.83 | 0.17 | 0 | 0 | 0 | 1 |
| **S13** | 0.54 | 0 | 0.46 | 0 | 0 | 0 |
| **S14** | 0.61 | 0 | 0.04 | 0 | 0.35 | 1 |
| **S15** | 0.72 | 0.28 | 0 | 0 | 0 | 1 |
| **S16** | 0.89 | 0 | 0.11 | 0 | 0 | 1 |
| **S17** | 0.23 | 0.57 | 0 | 0 | 0.2 | 0 |
| **S18** | 0.55 | 0.33 | 0.04 | 0 | 0.08 | 0 |
| **S19** | 0.46 | 0.49 | 0 | 0 | 0.06 | 0 |
| **S2** | 0.68 | 0.26 | 0.06 | 0 | 0 | 1 |
| **S20** | 1 | 0 | 0 | 0 | 0 | 1 |
| **S21** | 0.46 | 0.27 | 0.23 | 0 | 0.04 | 0 |

| | | | | | | |
|---|---|---|---|---|---|---|
| **S22** | 0.53 | 0.05 | 0 | 0 | 0.42 | 0 |
| **S23** | 0.31 | 0.45 | 0 | 0 | 0.24 | 0 |
| **S24** | 0.74 | 0.23 | 0.02 | 0 | 0.01 | 1 |
| **S3** | 0.53 | 0.44 | 0.02 | 0 | 0 | 0 |
| **S4** | 0.9 | 0.1 | 0 | 0 | 0 | 1 |
| **S5** | 0.61 | 0.36 | 0 | 0 | 0.03 | 1 |
| **S6** | 0.51 | 0.3 | 0 | 0 | 0.19 | 0 |
| **S7** | 0.5 | 0 | 0 | 0 | 0.5 | 0 |
| **S8** | 0.12 | 0.09 | 0 | 0 | 0.79 | 5 |
| **S9** | 0.47 | 0.02 | 0 | 0 | 0.51 | 0 |

**Table S12.** Details on the main global paleotemperatures estimates considered in this study. *Long-term curve based on equation 7b on Cramer et al. (2011).

| Reference | Data origin | Estimates | Data points | Time range (Mya) | Comments |
|---|---|---|---|---|---|
| **Hansen *et al.* 2013** (1) | $\delta^{18}O$ benthic foraminifera | Deep-sea and surface temperatures | 17,604 | 0 – 65.6 | |
| **Zachos *et al.* 2001** (5) | $\delta^{18}O$ benthic foraminifera | Deep-sea temperatures | 10,801 | 0 – 67.5 | |
| **Zachos *et al.* 2008** (6) | $\delta^{18}O$ benthic foraminifera | Deep-sea temperatures | 17,693 | 0 – 67.5 | Based on (5) |
| **Prokoph *et al.* 2008** (4) | $\delta^{18}O$ benthic foraminifera, brachiopods, belemnites, planktonic foraminifera, trilobites | Deep-sea temperatures | 55,000 | 0 – 522 (2450) | Based on (5) |
| **Veizer & Prokoph 2015** (3) | $\delta^{18}O$ benthic foraminifera, brachiopods, belemnites, planktonic foraminifera, bivalve, trilobites, oyster, corals | Deep-sea temperatures | 58,532 | 0 – 540 | Based on (4, 5) |
| **Cramer *et al.* 2009** (7) | $\delta^{18}O$ benthic foraminifera | Deep-sea temperatures | 35,354 | 0 – 112 | Accounts for fluctuations in sea water (sw) $\delta^{18}O_{sw}$ through time |
| **Cramer *et al.* 2011** (2) | $\delta^{18}O$ benthic foraminifera | Deep-sea temperatures | 624 | 0 – 62.4 | Accounts for fluctuations in seawater and corrected for ice volume* |

**Table S13.** Taxon sampling and GenBank accession numbers for caviomorph rodents.

| Species | | Genes | | | | |
|---|---|---|---|---|---|---|
| **MOUSE-RELATED CLADE** | | **cyt-b** | **12SrRNA** | **GHR** | **vWF** | **RAG1** |
| Pedetidae | *Pedetes spce* | AJ389527 | AY012113 | AF332025 | AJ238389 | AY011882 |
| CTENOHYSTRICA | | | | | | |
| CTENODACTYLOMORPHI | | | | | | |
| Diatomyidae | *Laonastes aenigmamus* | AM407933 | DQ139934 | AM407901 | AM407897 | |
| Ctenodactylidae | *Ctenodactylus vali* | AJ389532 | AJ389543 | AF332042 | JN415077 | JN633629 |
| | *Felovia vae* | KM369883 | | | | |
| | *Massoutiera mzabi* | AJ389533 | AJ389544 | AM407921 | AJ238388 | |
| HYSTRICOGNATHI | | | | | | |
| Hystricidae | *Atherurus africanus* | HQ450774 | AY093658 | | | |
| | *Atherurus macrourus* | FJ931121 | U12451 | JF938865 | AJ251131 | |
| | *Hystrix africaeaustralis* | X70674 | U12448 | AF332033 | | |
| | *Hystrix brachyura* | JQ991599 | AY012117 | | | |
| | *Hystrix cristata* | FJ472565 | AY093659 | JN414760 | JN415082 | AY011886 |
| | *Hystrix indica* | AY692229 | AY093669 | | | |
| | *Trichys fasciculata* | | | FM162081 | AJ224675 | |
| PHIOMORPHA | | | | | | |
| Bathyergidae | *Bathyergus janetta* | AF012241 | AY425843 | | | |
| | *Bathyergus suillus* | AY425913 | | FM162080 | AJ238384 | |
| | *Fukomys amatus* | AF012233 | AY427021 | | | |

| | | | | | | |
|---|---|---|---|---|---|---|
| | *Fukomys anselli* | | AY427022 | | | |
| | *Fukomys bocagei* | AF012229 | AF012213 | | | |
| | *Fukomys damarensis* | AF012220 | AY427026 | FN984748 | FN984751 | |
| | *Fukomys darlingi* | AF012232 | AY427033 | | | |
| | *Fukomys foxi* | | AY427036 | | | |
| | *Fukomys ilariae* | | | | | |
| | *Fukomys kafuensis* | | AY427037 | | | |
| | *Fukomys mechowi* | AF012230 | AY427041 | | | |
| | *Fukomys occlusus* | | | | | |
| | *Fukomys ochraceocinereus* | | AY427045 | | | |
| | *Fukomys vandewoestijneae* | | | | | |
| | *Fukomys whytei* | | AY425863 | AY427046 | | |
| | *Fukomys zechi* | | | | | |
| | *Cryptomys hottentotus* | AY425885 | AY427064 | FJ855202 | AJ251132 | |
| | *Cryptomys anomalous* | | AY427054 | | | |
| | *Cryptomys holosericeus* | | AY427051 | | | |
| | *Cryptomys natalensis* | | M63568 | | | |
| | *Cryptomys nimrodi* | | | | | |
| | *Georychus capensis* | AF012243 | AY427066 | FJ855203 | | |
| | *Heliophobius argenteocinereus* | KJ742646 | AY427070 | FJ855204 | AJ251133 | KJ742671 |
| | *Heterocephalus glaber* | AF155870 | AY427075 | AF332034 | AJ251134 | AY011889 |
| Petromuridae | *Petromus typicus* | DQ139935 | M63571 | JN414761 | AJ251144 | JN633636 |
| Thryonomyidae | *Thryonomys swinderianus* | KJ742647 | NC_002658 | AF332035 | AJ224674 | KJ742672 |
| CAVIOMORPHA | | | | | | |

| | | | | | | |
|---|---|---|---|---|---|---|
| ERETHIZONTOIDEA | | | | | | |
| Erethizontidae | | | | | | |
| Chaetomyinae | *Chaetomys subspinosus* | EU544660 | | | | |
| Erethizontinae | *Erethizon dorsatum* | | AY012118 | AF332037 | AJ251135 | AY011887 |
| | *Coendou bicolor* | KC463889 | | | | |
| | *Coendou ichillus* | KC463860 | | | | |
| | *Coendou insidiosus* | KC463861 | | | | |
| | *Coendou melanurus* | JX312693 | JX312693 | | | |
| | *Coendou mexicanus* | KC463862 | AJ389549 | | AJ224664 | |
| | *Coendou nycthemera* | KC463863 | | FJ855212 | | |
| | *Coendou prehensilis* | KC463864 | | | | |
| | *Coendou pruinosus* | KC463873 | AF520695 | AF520663 | | |
| | *Coendou quichua* | KC463880 | | | | |
| | *Coendou roosmalenorum* | KC463882 | | | | |
| | *Coendou spinosus* | KC463884 | | | | |
| | *Coendou vestitus* | KC463887 | | | | |
| | *Coendou villosus* | KC463888 | | | | |
| CAVIOIDEA | | | | | | |
| Caviidae | | | | | | |
| Caviinae | *Cavia aperea* | GU136753 | AF433908 | AF433930 | | |
| | *Cavia fulgida* | GU136737 | | | | |
| | *Cavia magna* | GU136734 | AY765986 | | | |
| | *Cavia porcellus* | AF490405 | AF433909 | AF433931 | AJ224663 | XM_003463833 |
| | *Cavia tschudii* | GU136731 | AY012121 | FJ855206 | | AY011890 |
| | *Galea musteloides* | KJ742648 | AF433910 | AF433933 | KJ742608 | KJ742673 |

| | | | | | | |
|---|---|---|---|---|---|---|
| | *Galea spixii* | GU067491 | AF433913 | AF433935 | | |
| | *Microcavia australis* | AF491750 | AF433915 | AF433937 | | |
| | *Microcavia niata* | GU136725 | | | | |
| Dolichotinae | *Dolichotis patagonum* | GU136724 | AF433917 | AF433939 | | |
| | Dolichotis salinicola | GU136723 | AF433919 | AF433940 | | |
| Hydrochoerinae | *Hydrochoerus hydrochaeris* | GU136721 | AF433924 | FJ855208 | AJ251137 | AY011891 |
| | *Hydrochoerus isthmius* | | | | | |
| | *Kerodon acrobata* | GU477346 | | | | |
| | *Kerodon rupestris* | GU136722 | AY765988 | AF433938 | | |
| Dasyproctidae | *Dasyprocta fuliginosa* | AF437784 | | | | |
| | *Dasyprocta leporina* | AF437791 | AY093660 | FJ855207 | U31607 | |
| | *Dasyprocta punctata* | | AF433921 | AF433943 | JN415079 | |
| | *Myoprocta acouchy* | KJ742649 | AF433922 | AF433945 | KJ742609 | KJ742695 |
| | *Myoprocta pratti* | U34850 | AF433923 | AF433946 | | |
| Cuniculidae | *Cuniculus paca* | AY206555 | AF520693 | AF433928 | AJ251136 | |
| | *Cuniculus taczanowskii* | KJ742656 | AY012125 | AF433929 | JN415074 | AY011894 |
| CHINCHILLOIDEA | | | | | | |
| Chinchillidae | *Chinchilla lanigera* | AF464760 | AF520696 | AF332036 | AJ238385 | KF590658 |
| | *Lagidium peruanum* | AY254885 | | | | |
| | *Lagidium viscacia* | AY254886 | | FJ855209 | | |
| | *Lagidium wolffsohni* | AY227023 | | | | |
| | *Lagostomus crassus* | | | | | |
| | *Lagostomus maximus* | AF245485 | | FJ855210 | | |
| Dinomyidae | *Dinomys branickii* | AY254884 | AY012124 | AF520659 | AJ251145 | AY011893 |

| | | | | | | |
|---|---|---|---|---|---|---|
| OCTODONTOIDEA | | | | | | |
| Abrocomidae | *Abrocoma bennettii* | AF244387 | | FJ855213 | AJ251143 | JN633625 |
| | *Abrocoma boliviensis* | KJ742657 | | | | |
| | *Abrocoma cinerea* | AF244388 | AF520666 | AF520643 | | |
| | *Cuscomys ashaninka* | KJ742658 | KJ742598 | KJ742626 | KJ742610 | KJ742683 |
| Octodontidae | *Aconaemys fuscus* | AF405351 | AF520674 | AF520657 | | |
| | *Aconaemys porteri* | | AF520671 | AF520644 | | |
| | *Aconaemys sagei* | KJ742650 | AF520672 | AF520645 | | KJ742675 |
| | *Octodon bridgesi* | KJ742651 | AF520676 | AF520646 | KJ742611 | KJ742676 |
| | *Octodon degus* | AF007058 | AF520678 | AF520647 | | |
| | *Octodon lunatus* | AF227514 | AF520681 | AF520650 | AJ238386 | |
| | *Octodontomys gliroides* | AF370706 | AF520683 | AF520649 | KF590672 | KF590663 |
| | *Octomys mimax* | GQ121097 | AF520686 | AF520652 | | |
| | *Pipanacoctomys aureus* | GQ121117 | AY249753 | AY249752 | | |
| | *Salinoctomys loschalchalerosorum* | KJ742652 | KJ742607 | KJ742635 | KJ742612 | KJ742684 |
| | *Spalacopus cyanus* | AF007061 | AF520688 | AF520653 | | |
| | *Tympanoctomys barrerae* | AF007060 | AF520691 | AF520655 | | |
| Ctenomyidae | *Ctenomys argentinus* | AF370680 | | | | |
| | *Ctenomys australis* | AF370697 | | | | |
| | *Ctenomys azarae* | JN791407 | | | | |
| | *Ctenomys bergi* | AF144284 | | | | |
| | *Ctenomys boliviensis* | AF007038 | U12446 | FJ855214 | | FJ855214 |
| | *Ctenomys bonettoi* | AF144286 | | | | |
| | *Ctenomys colburni* | HM777474 | | | | |
| | *Ctenomys conoveri* | AF007054 | | | | |

| | | | | | |
|---|---|---|---|---|---|
| *Ctenomys coyhaiquensis* | AF119112 | KF590700 | KF590678 | KF590666 | KF590659 |
| *Ctenomys dorbignyi* | AF500044 | | | | |
| *Ctenomys flamarioni* | AF119107 | | | | |
| Ctenomys fodax | HM777475 | | | | |
| Ctenomys frater | AF007045 | | | | |
| Ctenomys fulvus | AF370688 | | | | |
| Ctenomys goodfellowi | AF007050 | | | | |
| Ctenomys haigi | AF422920 | AF422853 | | | |
| Ctenomys ibicuiensis | JQ389020 | | | | |
| Ctenomys juris | AF144275 | | | | |
| Ctenomys lami | HM777477 | | | | |
| Ctenomys latro | HM777478 | | | | |
| Ctenomys leucodon | AF007056 | HM544131 | | | |
| Ctenomys lewisi | AF007049 | | | | |
| Ctenomys magellanicus | HM777479 | | | | |
| Ctenomys maulinus | AF370703 | | | AJ251138 | |
| Ctenomys mendocinus | HM777480 | | | | |
| Ctenomys minutus | HM777483 | | | | |
| Ctenomys nattereri | HM777484 | | | | |
| Ctenomys occultus | HM777485 | | | | |
| Ctenomys opimus | AF370700 | | | | |
| Ctenomys pearsoni | HM777486 | | | | |
| Ctenomys perrensi | HM777489 | | | | |
| Ctenomys pilarensis | AF144265 | | | | |
| Ctenomys porteousi | AF370682 | | | | |

| | | | | | | |
|---|---|---|---|---|---|---|
| | Ctenomys pundit | HM777490 | | | | |
| | Ctenomys rionegrensis | AF119103 | HM544130 | | | |
| | Ctenomys roigi | HM777492 | | | | |
| | Ctenomys saltarius | HM777493 | | | | |
| | Ctenomys scagliai | HM777494 | | | | |
| | Ctenomys sericeus | HM777496 | | | | |
| | Ctenomys sociabilis | HM777495 | HM544129 | | | |
| | Ctenomys steinbachi | AF007044 | AF520667 | AF520656 | | |
| | Ctenomys talarum | AF370699 | | | | |
| | Ctenomys torquatus | AF119111 | | | | |
| | Ctenomys tuconax | AF370684 | | | | |
| | Ctenomys tucumanus | AF370691 | | | | |
| | Ctenomys yolandae | AF144285 | | | | |
| Echimyidae | | | | | | |
| Dactylomyinae | *Dactylomys boliviensis* | L23339 | AF422875 | JX515334 | AJ849307 | EU313298 |
| | *Dactylomys dactylinus* | L23335 | AF422874 | KF590681 | KF590667 | EU313300 |
| | *Dactylomys peruanus* | EU313206 | | | | |
| | *Kannabateomys amblyonyx* | AF422917 | AF422850 | | AJ849310 | |
| | *Olallamys albicauda* | KF590697 | *Fabre et al. 2017* | KF590690 | KF590673 | |
| | *Olallamys edax* | *this study* | *this study* | | | |
| Echimyinae | *Callistomys pictus* | KJ742659 | KJ742594 | KJ742627 | KJ742614 | KJ742677 |
| | *Diplomys caniceps* | *this study* | *this study* | | | |
| | *Diplomys labilis* | KJ742660 | *Fabre et al. 2017* | KJ742636 | KJ742613 | KJ742685 |
| | *Santamartamys rufodorsalis* | KJ742664 | *Fabre et al. 2017* | | | |
| | *Echimys chrysurus* | L23341 | AF422877 | JX515333 | AJ251141 | EU313303 |

| | | | | | |
|---|---|---|---|---|---|
| *Echimys saturnus* | *this study* | *this study* | | | |
| *Pattonomys occasius* | KJ742661 | Emmons & Fabre 2018 | KJ742637 | | |
| *Pattonomys semivillosus* | KJ742662 | Emmons & Fabre 2018 | | KJ742616 | KJ742687 |
| *Isothrix barbarabrownae* | EU313214 | KF590701 | KF590682 | KF590668 | EU313304 |
| *Isothrix bistriata* | L23349 | | JX515336 | AJ849308 | EU313307 |
| *Isothrix negrensis* | L23355 | AF422873 | | | |
| *Isothrix orinoci ‡* | EU313223 | KF590702 | KF590683 | KF590669 | KF590660 |
| *Isothrix pagurus ‡* | EU313227 | KF590703 | KF590684 | KF590670 | KF590661 |
| *Isothrix sinnamariensis* | AY745734 | KF590704 | KF590685 | AJ849309 | EU313312 |
| *Toromys grandis* | KF590699 | Emmons & Fabre 2018 | KF590694 | KF590676 | EU313336 |
| *Toromys rhipidurus* | KJ742663 | Emmons & Fabre 2018 | KJ742638 | KJ742617 | KJ742686 |
| *Toromys albiventris* | Emmons & Fabre 2018 | Emmons & Fabre 2018 | | | |
| *Makalata didelphoides* | L23362 | KJ742600 | KJ742639 | AJ849311 | KJ742688 |
| *Makalata macrura* | L23356 | AF422879 | KF590687 | AJ849312 | EU313328 |
| *Phyllomys blainvillii* | JF297836 | KF590706 | KF590692 | JF297734 | KF590664 |
| *Phyllomys brasiliensis* | EF608182 | | | JF297729 | |
| *Phyllomys dasythrix* | JF297832 | KJ742605 | KJ742641 | JF297708 | KJ742689 |
| *Phyllomys lamarum* | EF608181 | | | JF297730 | |
| Phyllomys lundi | EF608183 | | | JF297721 | |
| Phyllomys mantiqueirensis | EF608179 | | | JF297720 | |
| *Phyllomys nigrispinus* | JF297807 | | | JF297714 | |
| *Phyllomys pattoni* | EF608187 | KJ742606 | KJ742642 | JF297744 | KJ742690 |
| Phyllomys sulinus | JF297833 | | | JF297710 | |
| Carterodon sulcidens | KJ742666 | KJ742596 | KJ742640 | KJ742615 | KJ742678 |
| Clyomys laticeps | AF422918 | KJ742597 | KJ742628 | AJ849306 | KJ742679 |

| | | | | | |
|---|---|---|---|---|---|
| Euryzygomatomys spinosus | EU544667 | AF422854 | KJ742629 | AJ849319 | KJ742680 |
| Hoplomys gymnurus | AF422922 | AF520668 | AF520661 | JN415080 | JN633632 |
| Lonchothrix emiliae | AF422921 | AF422857 | | | |
| Mesomys hispidus | KF590705 | KF590696 | KF590688 | KF590671 | KF590662 |
| Mesomys occultus | L23388 | AF422858 | KF590689 | | EU313331 |
| Mesomys stimulax | KJ742667 | KJ742603 | KJ742630 | KJ742618 | KJ742674 |
| Myocastor coypus | EU544663 | AF520669 | AF520662 | AJ251140 | AY011892 |
| Proechimys brevicauda | *this study* | *this study* | | | |
| Proechimys cuvieri | AJ251400 | KF590707 | KF590693 | KF590675 | KF590665 |
| Proechimys echinothrix | *this study* | *this study* | | | |
| Proechimys gardneri | *this study* | *this study* | | | |
| Proechimys goeldii | *this study* | *this study* | | | |
| Proechimys guairae | *this study* | *this study* | | | |
| Proechimys guyannensis | AJ251396 | *this study* | | | |
| Proechimys hoplomyoides | *this study* | *this study* | | | |
| Proechimys kulinae | *this study* | *this study* | | | |
| Proechimys longicaudatus | HM544128 | HM544128 | KJ742643 | KJ742619 | KJ742681 |
| Proechimys mincae | *this study* | *this study* | | | |
| Proechimys pattoni | *this study* | *this study* | | | |
| Proechimys poliopus | *this study* | *this study* | | | |
| Proechimys quadruplicatus | U35413 | AF422863 | | AJ849313 | |
| Proechimys roberti | *this study* | *this study* | | AJ251139 | |
| Proechimys semispinosus | *this study* | *this study* | | | |
| Proechimys simonsi | U35414 | AF422864 | KJ742631 | AJ849320 | EU313332 |
| Proechimys steerei | *this study* | *this study* | | | |

| | | | | | | |
|---|---|---|---|---|---|---|
| | Proechimys trinitatus | *this study* | *this study* | | | |
| | Thrichomys apereoides | EU313252 | AF422855 | JX515325 | AJ849315 | EU313334 |
| | Thrichomys inermis | AY083343 | | | | |
| | Thrichomys pachyurus | AY083329 | | | | |
| | Trinomys albispinus | U34856 | | | | |
| | Trinomys dimidiatus | U35169 | AF422867 | | KJ742620 | KJ742682 |
| | Trinomys eliasi | U35166 | AF422869 | | | |
| | Trinomys gratiosus | AF194281 | | | | |
| | Trinomys iheringi | EU313254 | AF422868 | KF590695 | KF590677 | EU313337 |
| | Trinomys paratus | U35165 | AF422866 | | AJ849316 | |
| | Trinomys setosus | AF422924 | AF422871 | | AJ849317 | |
| | Trinomys yonenagae | AF194295 | AF422865 | | AJ849318 | |
| Capromyidae | | | | | | |
| Capromyinae | *Capromys pilorides* | AF422915 | AF433926 | AF433950 | AJ251142 | JN633628 |
| | *Geocapromys browni* | KJ742653 | KJ742599 | KJ742644 | KJ742621 | KJ742692 |
| | *Geocapromys ingrahami* | KJ742668 | | | | |
| | *Geocapromys thoracatus* | *this study* | *this study* | | | |
| | *Mesocapromys angelcabrerai* | KJ742654 | KJ742595 | KJ742632 | KJ742622 | KJ742694 |
| | *Mesocapromys auritus* | KJ742655 | KJ742601 | KJ742633 | KJ742623 | KJ742693 |
| | *Mesocapromys melanurus* | KJ742669 | KJ742602 | | | KJ742691 |
| | *Mesocapromys nanus* | *this study* | *this study* | | | |
| | *Mysateles prehensilis* | KJ742670 | KJ742604 | KJ742634 | KJ742624 | KJ742696 |
| Plagiodontinae | *Plagiodontia aedium* | KJ742665 | | KJ742645 | KJ742625 | KJ742697 |

### References for SI text

1. J. Hansen, M. Sato, G. Russell, P. Kharecha, Climate sensitivity, sea level and atmospheric carbon dioxide. *Philos. Trans. R. Soc. A Math. Phys. Eng. Sci.* **371**, 20120294 (2013).
2. B. S. Cramer, K. G. Miller, P. J. Barrett, J. D. Wright, Late Cretaceous-Neogene trends in deep ocean temperature and continental ice volume: Reconciling records of benthic foraminiferal geochemistry (δ18O and Mg/Ca) with sea level history. *J. Geophys. Res. Ocean.* **116**, C12023 (2011).
3. J. Veizer, A. Prokoph, Temperatures and oxygen isotopic composition of Phanerozoic oceans. *Earth-Science Rev.* **146**, 92–104 (2015).
4. A. Prokoph, G. A. Shields, J. Veizer, Compilation and time-series analysis of a marine carbonate δ18O, δ13C, 87Sr/86Sr and δ34S database through Earth history. *Earth-Science Rev.* **87**, 113–133 (2008).
5. J. Zachos, M. Pagani, L. Sloan, E. Thomas, K. Billups, Trends, Rhythms, and Aberrations in Global Climate 65 Ma to Present. *Science* **292**, 686–693 (2001).
6. J. C. Zachos, G. R. Dickens, R. E. Zeebe, An early Cenozoic perspective on greenhouse warming and carbon-cycle dynamics. *Nature* **451**, 279–283 (2008).
7. B. S. Cramer, J. R. Toggweiler, J. D. Wright, M. E. Katz, K. G. Miller, Ocean overturning since the late cretaceous: Inferences from a new benthic foraminiferal isotope compilation. *Paleoceanography* **24**, PA4216 (2009).
8. D. M. Olson, *et al.*, Terrestrial Ecoregions of the World: A New Map of Life on Earth. *Bioscience* **51**, 933 (2001).
9. C. A. W. C. W. Kilpatrick, Infraorder Hystricognathi Brandt, 1855. *Mammal species world a Taxon. Geogr. Ref. (D. E Wilsonand D. M Reeder eds.). 3rd ed. Johns Hopkins Univ. Press. Balt. Maryl.*, 1538–1600 (2005).
10. C. J. Burgin, J. P. Colella, P. L. Kahn, N. S. Upham, How many species of mammals are there? *J. Mammal.* **99**, 1–14 (2018).
11. N. S. Upham, B. D. Patterson, Evolution of caviomorph rodents: a complete phylogeny and timetree for living genera. *Biol. caviomorph rodents Divers. Evol. Buenos Aires SAREM Ser. A* **1**, 63–120 (2015).
12. P.-H. Fabre, *et al.*, Rodents of the Caribbean: origin and diversification of hutias unravelled by next-generation museomics. *Biol. Lett.* **10**, 20140266

(2014).
13. P. Fabre, T. Galewski, M. Tilak, E. J. P. Douzery, Diversification of South American spiny rats (Echimyidae): a multigene phylogenetic approach. *Zool. Scr.* **42**, 117–134 (2013).
14. D. Huchon, F. M. Catzeflis, E. J. Douzery, Molecular evolution of the nuclear von Willebrand factor gene in mammals and the phylogeny of rodents. *Mol. Biol. Evol.* **16**, 577–589 (1999).
15. D. Huchon, E. J. P. Douzery, From the Old World to the New World: a molecular chronicle of the phylogeny and biogeography of hystricognath rodents. *Mol. Phylogenet. Evol.* **20**, 238–251 (2001).
16. P.-H. Fabre, L. Hautier, D. Dimitrov, E. J. P Douzery, A glimpse on the pattern of rodent diversification: A phylogenetic approach. *BMC Evol. Biol.* **12**, 88 (2012).
17. M.-K. Tilak, *et al.*, A cost-effective straightforward protocol for shotgun Illumina libraries designed to assemble complete mitogenomes from non-model species. *Conserv. Genet. Resour.* **7**, 37–40 (2015).
18. M. Kearse, *et al.*, Geneious Basic: an integrated and extendable desktop software platform for the organization and analysis of sequence data. *Bioinformatics* **28**, 1647–1649 (2012).
19. N. Galtier, M. Gouy, C. Gautier, SEA VIEW and PHYLO_ WIN: Two graphic tools for sequence alignment and molecular phytogeny. *Comput. Appl. Biosci.* **12**, 543–548 (1996).
20. V. Ranwez, E. J. P. Douzery, C. Cambon, N. Chantret, F. Delsuc, MACSE v2: toolkit for the alignment of coding sequences accounting for frameshifts and stop codons. *Mol. Biol. Evol.* **35**, 2582–2584 (2018).
21. A. Stamatakis, RAxML-VI-HPC: Maximum likelihood-based phylogenetic analyses with thousands of taxa and mixed models. *Bioinformatics* **22**, 2688–2690 (2006).
22. A. Stamatakis, P. Hoover, J. Rougemont, A rapid bootstrap algorithm for the RAxML web servers. *Syst. Biol.* **57**, 758–771 (2008).
23. R. Lanfear, P. B. Frandsen, A. M. Wright, T. Senfeld, B. Calcott, PartitionFinder 2: new methods for selecting partitioned models of evolution for molecular and morphological phylogenetic analyses. *Mol. Biol. Evol.* **34**, 772–773 (2016).

24. A. J. Drummond, A. Rambaut, BEAST: Bayesian evolutionary analysis by sampling trees. *BMC Evol. Biol.* **7**, 214 (2007).
25. A. R. Lemmon, J. M. Brown, K. Stanger-Hall, E. M. Lemmon, The effect of missing data on phylogenetic estimates obtained by maximum likelihood and Bayesian interference. *Syst. Biol.* **58**, 130–145 (2009).
26. A. Rambaut, A. J. Drummond, Tracer, version 1.5, MCMC trace analysis package. Available: http://tree.bio.ed.ac.uk/software/ (2009).
27. M. J. Benton, The Red Queen and the Court Jester: species diversity and the role of biotic and abiotic factors through time. *Science* **323**, 728–732 (2009).
28. J. F. Parham, *et al.*, Best practices for justifying fossil calibrations. *Syst. Biol.* **61**, 346–359 (2011).
29. A. E. Zanne, *et al.*, Three keys to the radiation of angiosperms into freezing environments. *Nature* **506**, 89–92 (2014).
30. , The Plant List (2010). Version 1. Published on the Internet; http://www.theplantlist.org/ (accessed May 2019).
31. T. E. Särkinen, *et al.*, Recent oceanic long-distance dispersal and divergence in the amphi-Atlantic rain forest genus *Renealmia* L.f. (Zingiberaceae). *Mol. Phylogenet. Evol.* **44**, 968–980 (2007).
32. C. D. Specht, D. W. Stevenson, A new phylogeny-based generic classification of Costaceae (Zingiberales). *Taxon* **55**, 153–163 (2006).
33. M. D. Pirie, P. J. M. Maas, R. A. Wilschut, H. Melchers-Sharrott, L. W. Chatrou, Parallel diversifications of *Cremastosperma* and *Mosannona* (Annonaceae), tropical rainforest trees tracking Neogene upheaval of South America. *R. Soc. Open Sci.* **5**, 171561 (2018).
34. P. E. Berry, W. J. Hahn, K. J. Sytsma, J. C. Hall, A. Mast, Phylogenetic relationships and biogeography of *Fuchsia* (Onagraceae) based on noncoding nuclear and chloroplast DNA data. *Am. J. Bot.* **91**, 601–614 (2004).
35. G. Davidse, *Flora mesoamericana. 4: Parte 1. Cucurbitaceae a Polemoniaceae* (Univ. Nacional Autónoma de México, Inst. de Biología, 2009).
36. , No Title. *Tropicos.org* (2008).
37. M. F. Simon, *et al.*, Recent assembly of the Cerrado, a Neotropical plant diversity hotspot, by in situ evolution of adaptations to fire. *Proc. Natl. Acad. Sci.* **106**, 20359–20364 (2009).

38. G. Stride, S. Nylinder, U. Swenson, Revisiting the biogeography of *Sideroxylon* (Sapotaceae) and an evaluation of the taxonomic status of *Argania* and *Spiniluma*. *Aust. Syst. Bot.* **27**, 104–118 (2014).
39. Y.-Y. Huang, S. A. Mori, L. M. Kelly, Toward a phylogenetic-based generic classification of Neotropical Lecythidaceae-I. Status of *Bertholletia*, *Corythophora*, *Eschweilera* and *Lecythis*. *Phytotaxa* **203**, 85–121 (2015).
40. T. E. Särkinen, "Historical assembly of seasonally dry tropical forest diversity in the tropical Andes," University of Oxford. (2010).
41. A. Antonelli, J. A. A. Nylander, C. Persson, I. Sanmartín, Tracing the impact of the Andean uplift on Neotropical plant evolution. *Proc. Natl. Acad. Sci. U. S. A.* **106**, 9749–9754 (2009).
42. M. Lavin, M. F. Wojciechowski, P. Gasson, C. Hughes, E. Wheeler, Phylogeny of robinioid legumes (Fabaceae) revisited: *Coursetia* and *Gliricidia* recircumscribed, and a biogeographical appraisal of the Caribbean endemics. *Syst. Bot.* **28**, 387–409 (2003).
43. R. H. J. Erkens, J. W. Maas, T. L. P. Couvreur, From Africa via Europe to South America: migrational route of a species-rich genus of Neotropical lowland rain forest trees (*Guatteria*, Annonaceae). *J. Biogeogr.* **36**, 2338–2352 (2009).
44. A. Antonelli, I. Sanmartín, Mass extinction, gradual cooling, or rapid radiation? Reconstructing the spatiotemporal evolution of the ancient angiosperm genus *Hedyosmum* (Chloranthaceae) using empirical and simulated approaches. *Syst. Biol.* **60**, 596–615 (2011).
45. L. P. Lagomarsino, F. L. Condamine, A. Antonelli, A. Mulch, C. C. Davis, The abiotic and biotic drivers of rapid diversification in Andean bellflowers (Campanulaceae). *New Phytol.* **210**, 1430–1442 (2016).
46. F. L. Condamine, N. S. Nagalingum, C. R. Marshall, H. Morlon, Origin and diversification of living cycads: A cautionary tale on the impact of the branching process prior in Bayesian molecular dating. *BMC Evol. Biol.* **15**, 65 (2015).
47. C. S. Drummond, R. J. Eastwood, S. T. S. Miotto, C. E. Hughes, Multiple continental radiations and correlates of diversification in *Lupinus* (Leguminosae): testing for key innovation with incomplete taxon sampling. *Syst. Biol.* **61**, 443–460 (2012).

48. M. C. M. P. de Medeiros, L. G. Lohmann, Phylogeny and biogeography of *Tynanthus* Miers (Bignonieae, Bignoniaceae). *Mol. Phylogenet. Evol.* **85**, 32–40 (2015).
49. S. Faurby, W. L. Eiserhardt, W. J. Baker, J.-C. Svenning, An all-evidence species-level supertree for the palms (Arecaceae). *Mol. Phylogenet. Evol.* **100**, 57–69 (2016).
50. M. F. Simon, *et al.*, The evolutionary history of *Mimosa* (Leguminosae): toward a phylogeny of the sensitive plants. *Am. J. Bot.* **98**, 1201–1221 (2011).
51. O. A. Pérez-Escobar, *et al.*, Recent origin and rapid speciation of Neotropical orchids in the world's richest plant biodiversity hotspot. *New Phytol.* **215**, 891–905 (2017).
52. E. L. Spriggs, P.-A. Christin, E. J. Edwards, C4 photosynthesis promoted species diversification during the Miocene grassland expansion. *PLoS One* **9**, e97722 (2014).
53. K. G. Dexter, *et al.*, Dispersal assembly of rain forest tree communities across the Amazon basin. *Proc. Natl. Acad. Sci.* **114**, 2645–2650 (2017).
54. M. L. Serrano-Serrano, J. Rolland, J. L. Clark, N. Salamin, M. Perret, Hummingbird pollination and the diversification of angiosperms: an old and successful association in Gesneriaceae. *Proc. R. Soc. B Biol. Sci.* **284**, 20162816 (2017).
55. P. V. A. Fine, F. Zapata, D. C. Daly, Investigating processes of Neotropical rain forest tree diversification by examining the evolution and historical biogeography of the Protieae (Burseraceae). *Evolution* **68**, 1988–2004 (2014).
56. T. Särkinen, L. Bohs, R. G. Olmstead, S. Knapp, A phylogenetic framework for evolutionary study of the nightshades (Solanaceae): a dated 1000-tip tree. *BMC Evol. Biol.* **13**, 214 (2013).
57. D. L. De-Silva, *et al.*, North Andean origin and diversification of the largest ithomiine butterfly genus. *Sci. Rep.* **7**, 45966 (2017).
58. T. L. Weese, L. Bohs, A three-gene phylogeny of the genus *Solanum* (Solanaceae). *Syst. Bot.* **32**, 445–463 (2007).
59. R. J. Schley, *et al.*, Is Amazonia a 'museum'for Neotropical trees? The evolution of the *Brownea* clade (Detarioideae, Leguminosae). *Mol. Phylogenet. Evol.* **126**, 279–292 (2018).
60. A. C. Martins, M. D. Scherz, S. S. Renner, Several origins of floral oil in the

Angelonieae, a southern hemisphere disjunct clade of Plantaginaceae. *Am. J. Bot.* **101**, 2113–2120 (2014).

61. M. F. Santos, *et al.*, Biogeographical patterns of *Myrcia* sl (Myrtaceae) and their correlation with geological and climatic history in the Neotropics. *Mol. Phylogenet. Evol.* **108**, 34–48 (2017).
62. R. Govaerts, M. Sobral, P. Ashton, F. Barrie, World checklist of Myrtaceae. Royal Botanic Gardens (2016).
63. R. S. Couto, *et al.*, Time calibrated tree of *Dioscorea* (Dioscoreaceae) indicates four origins of yams in the Neotropics since the Eocene. *Bot. J. Linn. Soc.* **188**, 144–160 (2018).
64. R. Arévalo, B. W. van Ee, R. Riina, P. E. Berry, A. C. Wiedenhoeft, Force of habit: shrubs, trees and contingent evolution of wood anatomical diversity using *Croton* (Euphorbiaceae) as a model system. *Ann. Bot.* **119**, 563–579 (2017).
65. H.-J. Esser, P. E. Berry, R. Riina, EuphORBia: a global inventory of the spurges. *Blumea-Biodiversity, Evol. Biogeogr. Plants* **54**, 11–12 (2009).
66. Y. Wang, *et al.*, Phylogeny and infrageneric classification of *Symplocos* (Symplocaceae) inferred from DNA sequence data. *Am. J. Bot.* **91**, 1901–1914 (2004).
67. G. C. Gibb, *et al.*, Shotgun mitogenomics provides a reference phylogenetic framework and timescale for living xenarthrans. *Mol. Biol. Evol.* **33**, 621–642 (2016).
68. D. E. Wilson, D. M. Reeder, *Mammal species of the world: a taxonomic and geographic reference* (JHU Press, 2005).
69. D. Rojas, O. M. Warsi, L. M. DaVALOS, Bats (Chiroptera: Noctilionoidea) challenge a recent origin of extant neotropical diversity. *Syst. Biol.* **65**, 432–448 (2016).
70. O. R. P. Bininda-Emonds, *et al.*, The delayed rise of present-day mammals. *Nature* **446**, 507–512 (2007).
71. M. S. Springer, *et al.*, Macroevolutionary dynamics and historical biogeography of primate diversification inferred from a species supermatrix. *PLoS One* **7**, e49521 (2012).
72. A. B. Rylands, R. A. Mittermeier, "The Diversity of the New World primates (Platyrrhini): an annotated taxonomy BT - South American primates:

comparative perspectives in the study of behavior, ecology, and conservation” in P. A. Garber, A. Estrada, J. C. Bicca-Marques, E. W. Heymann, K. B. Strier, Eds. (Springer New York, 2009), pp. 23–54.

73. S. A. Jansa, F. K. Barker, R. S. Voss, The early diversification history of didelphid marsupials: A window into South America’s “splendid isolation.” *Evolution* **68**, 684–695 (2014).

74. R. Maestri, *et al.*, The ecology of a continental evolutionary radiation: Is the radiation of sigmodontine rodents adaptive? *Evolution* **71**, 610–632 (2017).

75. B. K. Lim, Divergence times and origin of neotropical sheath-tailed bats (tribe Diclidurini) in South America. *Mol. Phylogenet. Evol.* **45**, 777–791 (2007).

76. O. Toljagić, K. L. Voje, M. Matschiner, L. H. Liow, T. F. Hansen, Millions of years behind: slow adaptation of ruminants to grasslands. *Syst. Biol.* **67**, 145–157 (2017).

77. K. Nyakatura, O. R. P. Bininda-Emonds, Updating the evolutionary history of Carnivora (Mammalia): a new species-level supertree complete with divergence time estimates. *BMC Biol.* **10**, 12 (2012).

78. N. S. Upham, B. D. Patterson, Diversification and biogeography of the Neotropical caviomorph lineage Octodontoidea (Rodentia: Hystricognathi). *Mol. Phylogenet. Evol.* **63**, 417–429 (2012).

79. W. Jetz, G. H. Thomas, J. B. Joy, K. Hartmann, A. O. Mooers, The global diversity of birds in space and time. *Nature* **491**, 444–448 (2012).

80. J. V Remsen Jr, HBW and BirdLife International Illustrated Checklist of the Birds of the World Volume 1: Non-passerines (2015).

81. J. A. McGuire, *et al.*, Molecular phylogenetics and the diversification of hummingbirds. *Curr. Biol.* **24**, 910–916 (2014).

82. E. P. Derryberry, *et al.*, Lineage diversification and morphological evolution in a large-scale continental radiation: The Neotropical ovenbirds and woodcreepers (aves: furnariidae). *Evolution* **65**, 2973–2986 (2011).

83. D. L. Slager, C. J. Battey, R. W. Bryson Jr, G. Voelker, J. Klicka, A multilocus phylogeny of a major New World avian radiation: the Vireonidae. *Mol. Phylogenet. Evol.* **80**, 95–104 (2014).

84. R. A. Pyron, F. T. Burbrink, Early origin of viviparity and multiple reversions to oviparity in squamate reptiles. *Ecol. Lett.* **17**, 13–21 (2014).

85. P. Uetz, P. Freed, J. Hošek, The Reptile Database, http://www.reptile-

database.org, accessed 2018.
86. C. R. Hutter, S. M. Lambert, J. J. Wiens, Rapid diversification and time explain Amphibian richness at different scales in the tropical Andes, Earth's most biodiverse Hotspot. *Am. Nat.* **190**, 828–843 (2017).
87. , AmphibiaWeb. 2019. <https://amphibiaweb.org> University of California, Berkeley, CA, USA. Accessed 2018.
88. D. R. Frost, Amphibian Species of the World: an online reference. http://research.amnh.org/herpetology/amphibia/index.html. Accessed 2018. *Am. Museum Nat. Hist. New York, USA* (2019).
89. S. N. Stuart, *Threatened amphibians of the world* (Lynx Edicions, 2008).
90. R. A. Pyron, Biogeographic analysis reveals ancient continental vicariance and recent oceanic dispersal in amphibians. *Syst. Biol.* **63**, 779–797 (2014).
91. S. Louca, M. W. Pennell, Extant timetrees are consistent with a myriad of diversification histories. *Nature* **580**, 502–505 (2020).
92. S. Louca, *et al.*, Bacterial diversification through geological time. *Nat. Ecol. Evol.* **2**, 1458–1467 (2018).

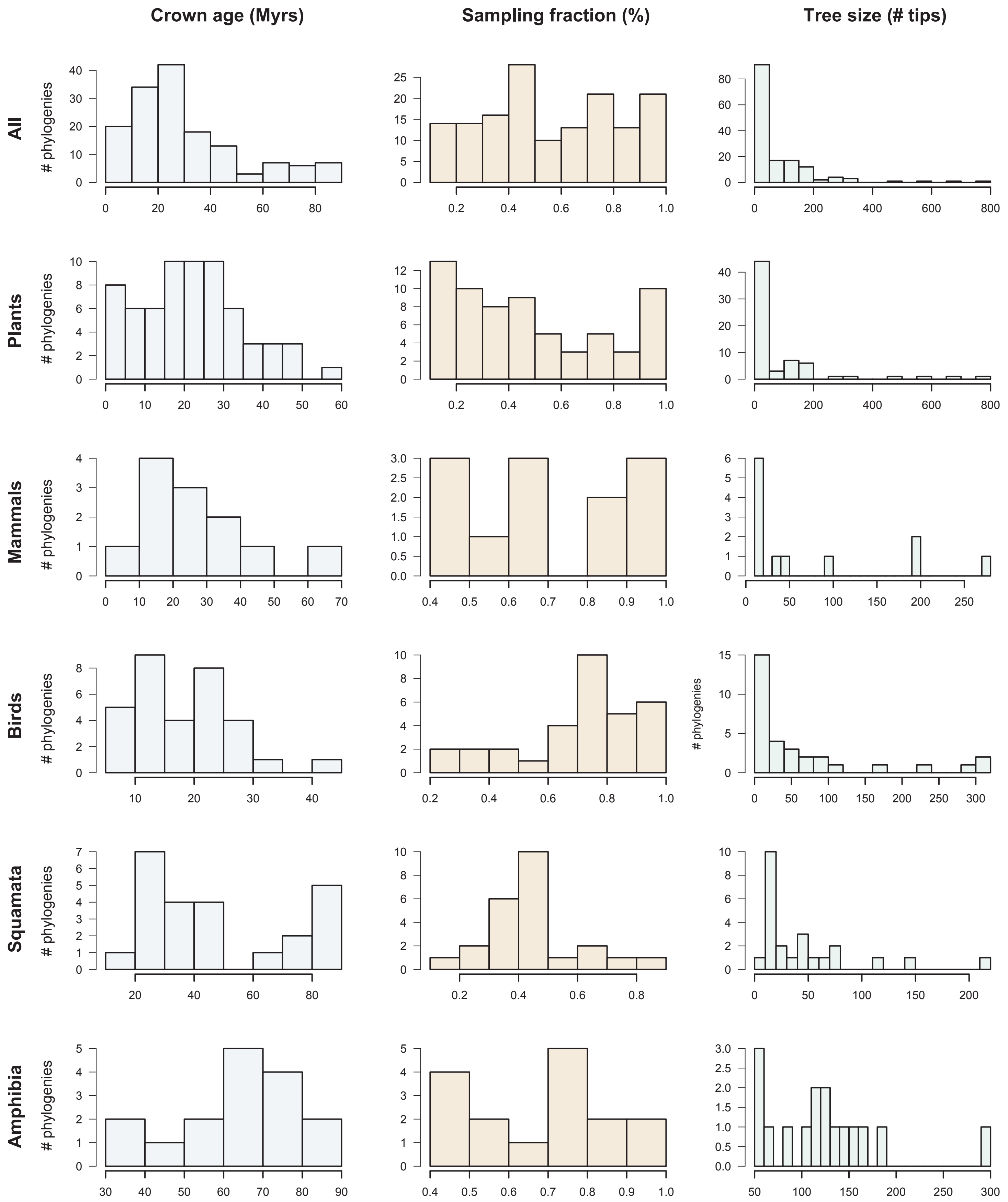

**Figure S1.** Dataset overview. The histograms represent the clade age (crown age in million years "Myrs"), sampling fraction (% of described species represented in the trees) and number of tips for the 150 phylogenies of Plants, Mammals, Birds, Squamata and Amphibian analysed here.

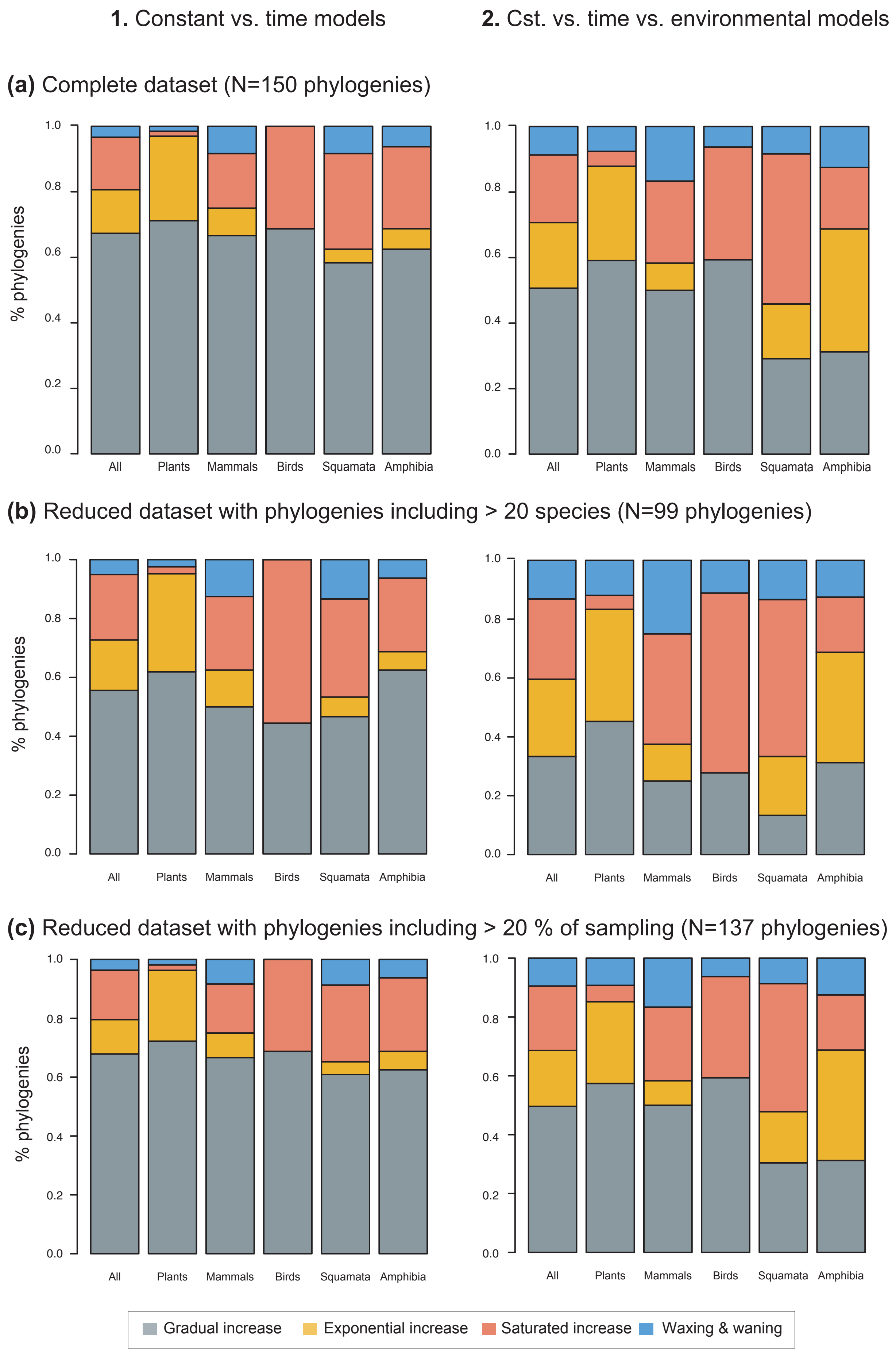

**Figure S2. Species richness dynamics on 150 phylogenies of Neotropical plants and tetrapods.** The histograms show the proportion of phylogenies best fitting gradual increase (Sc. 1), exponential increase (Sc. 2), saturated increase (Sc. 3) and waxing & waning (Sc. 4) species richness dynamics, as derived from diversification rates when comparing **(1)** time-dependent against constant models, and **(2)** environmental (time- and uplift dependent) models against time-dependent and constant models. The results are reported for **(a)** the complete dataset , **(b)** a reduced dataset including only trees with more than 20 species**,** and **(c)** a reduced dataset including only trees with a sampling fraction over 20%. Abbreviations: Cst = constant.

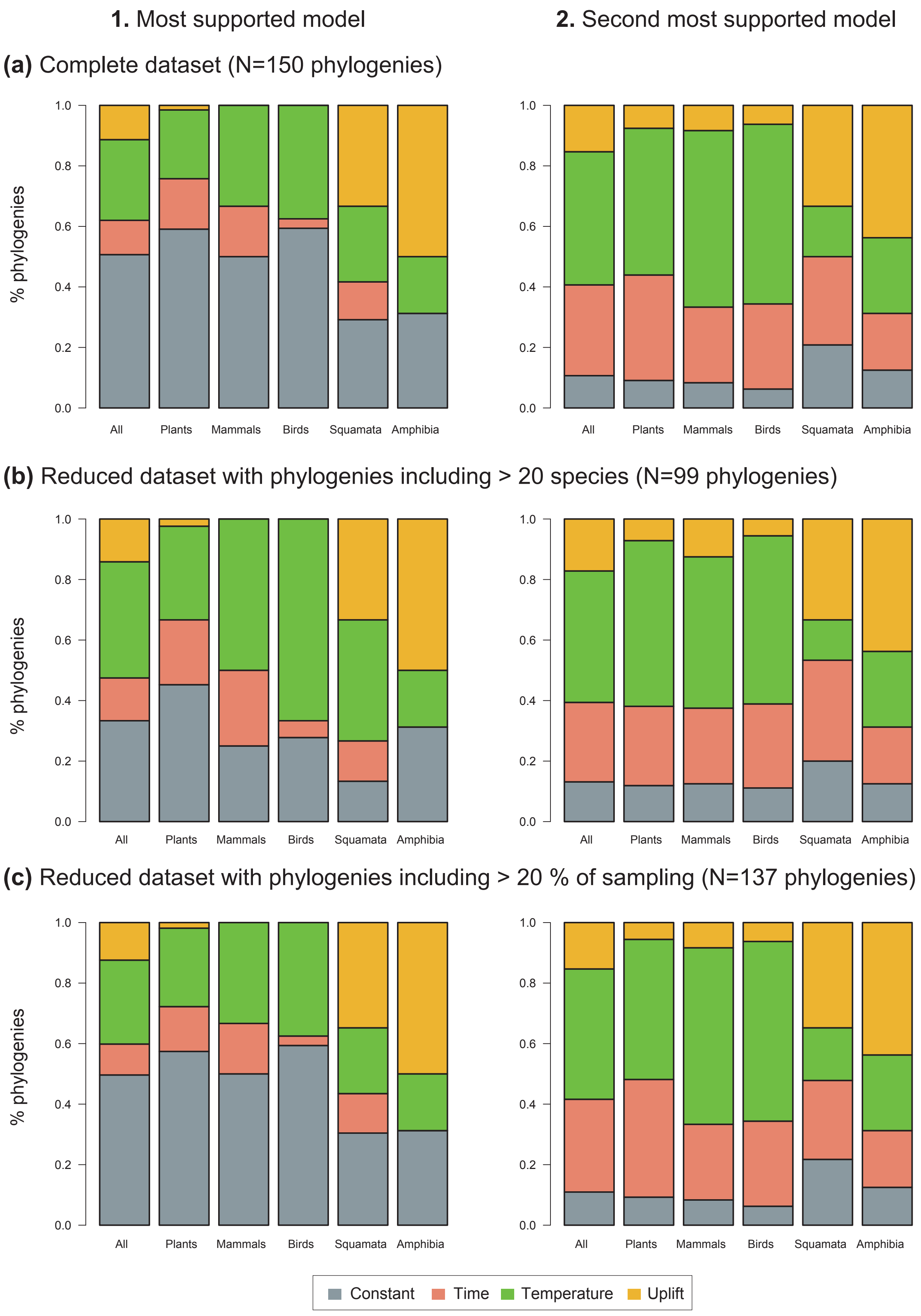

**Figure S3. Drivers of Neotropical diversification.** The histograms report the proportion of phylogenies whose diversification rates are best explained by a model with constant, time-dependent, temperature-dependent, or uplift-dependent diversification based on **(1)** the most supported model (lowest AIC value), and **(2)** the second most supported model. The results are reported for the complete dataset **(a)**, for a reduced dataset including only trees with more than 20 species **(b),** and for a reduced dataset including only trees with a sampling fraction over 20% **(c)**.

**(a)** **Paleotemperature estimates**

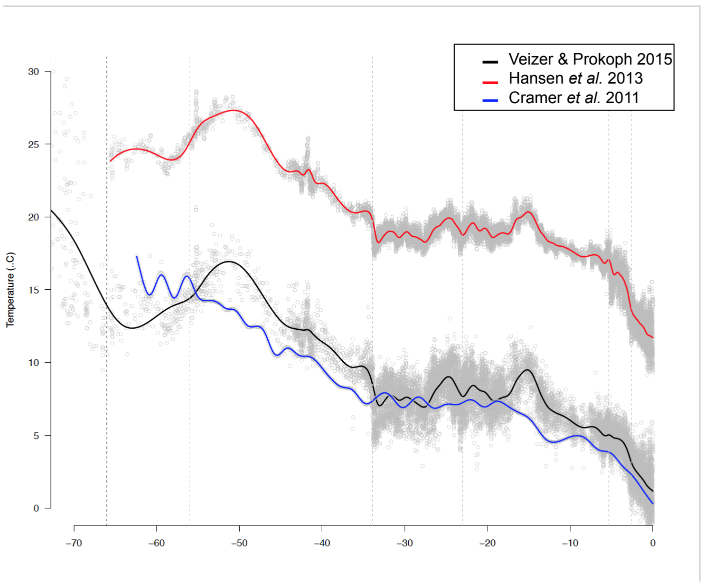

**(b)** **Diversification drivers**

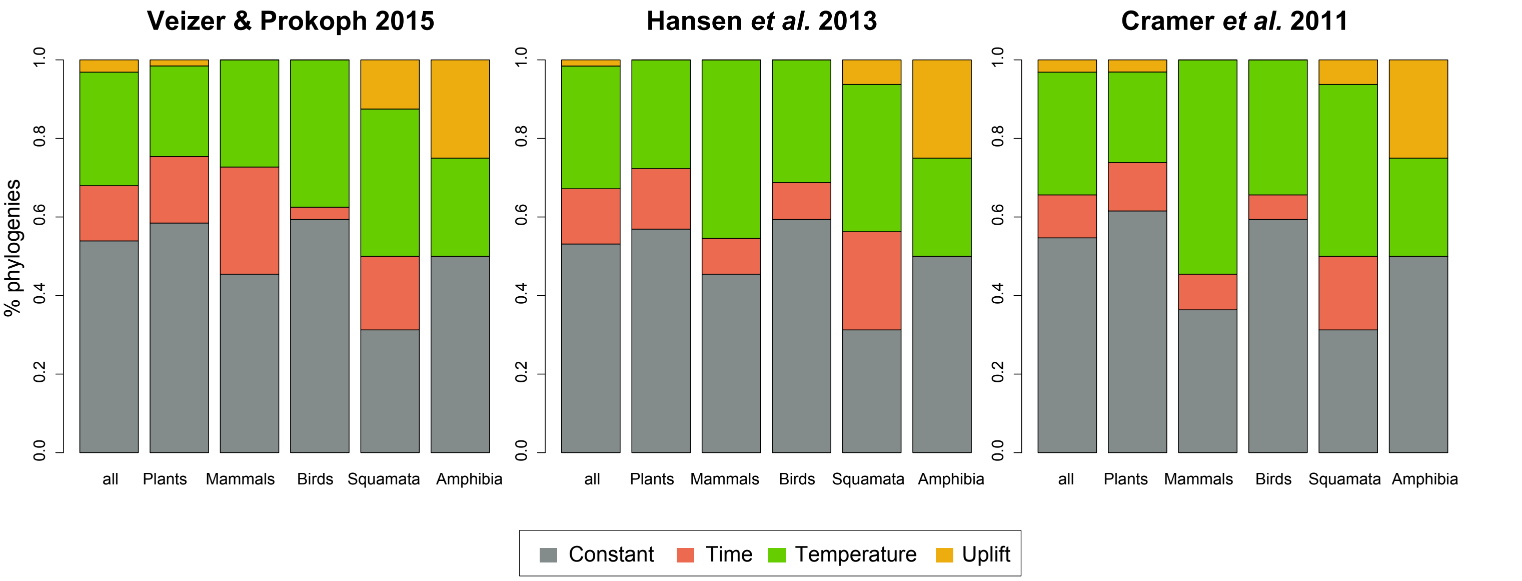

**Figure S4**. Result comparisons when the temperature-dependency of diversification rates is estimated based on the paleo-temperature curve of Veizer & Prokoph (2015), or Cramer et al. (2011) or Hansen et al (2013), see Methods for details. **(a)** Mean global paleotemperature estimates mostly differ in the magnitude of the changes but share the same overall trend. The histograms report **(b)** the proportion of phylogenies best supported by a model with diversification rates that are constant (in red), time- (in blue), temperature- (in green), or Andean- (in orange) compared for the analyses based on different temperatures estimates.

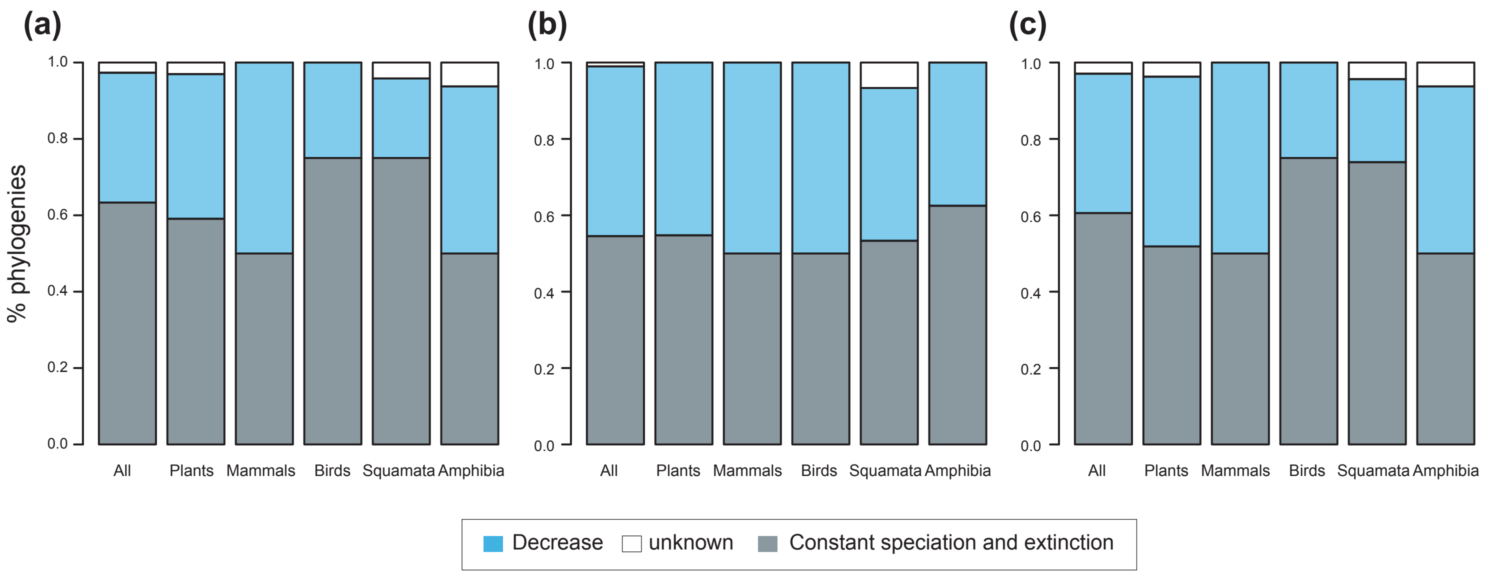

**Figure S5. Speciation trends on 150 phylogenies of Neotropical plants and tetrapods.** The histograms show the proportion of phylogenies best fitting constant vs. time-variable diversification models, as derived from pulled diversification rates for **(a)** the complete dataset, **(b)** a reduced dataset including only trees with more than 20 species (N=99), and **(c)** a reduced dataset including only trees with a sampling fraction over 20% (N=137). Speciation trends are deriverd from present-day pulled extinction rates *μp(0):* negative present-day pulled extinction rates values (*μp(0)* < 0) indicate decreasing speciation trends through time (Louca & Pennell, 2020). Positive *μp(0)* > 0 values are possible under both increasing or decreasing speciation rates, in which case speciation trends are designed as "unknown".

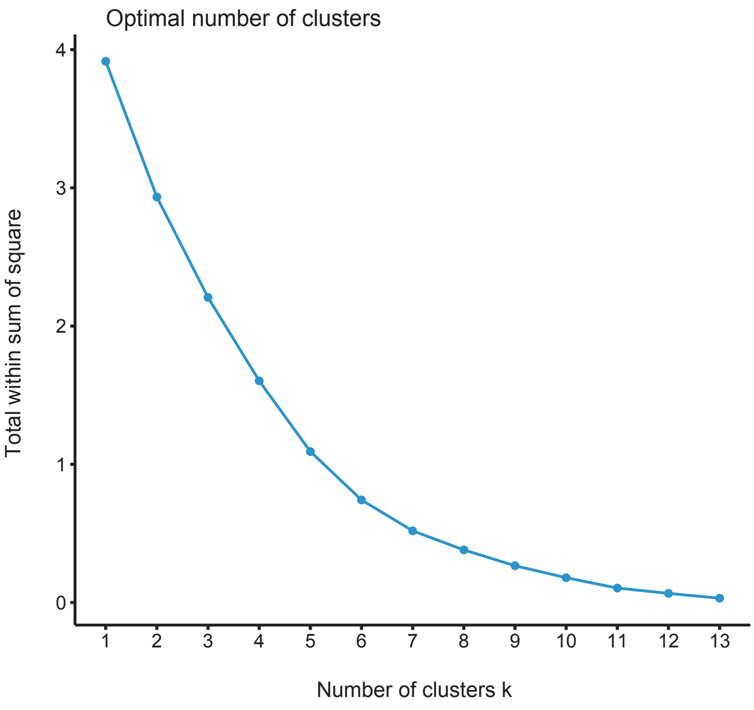

**Figure S6.** Elbow curve for K means clustering results.

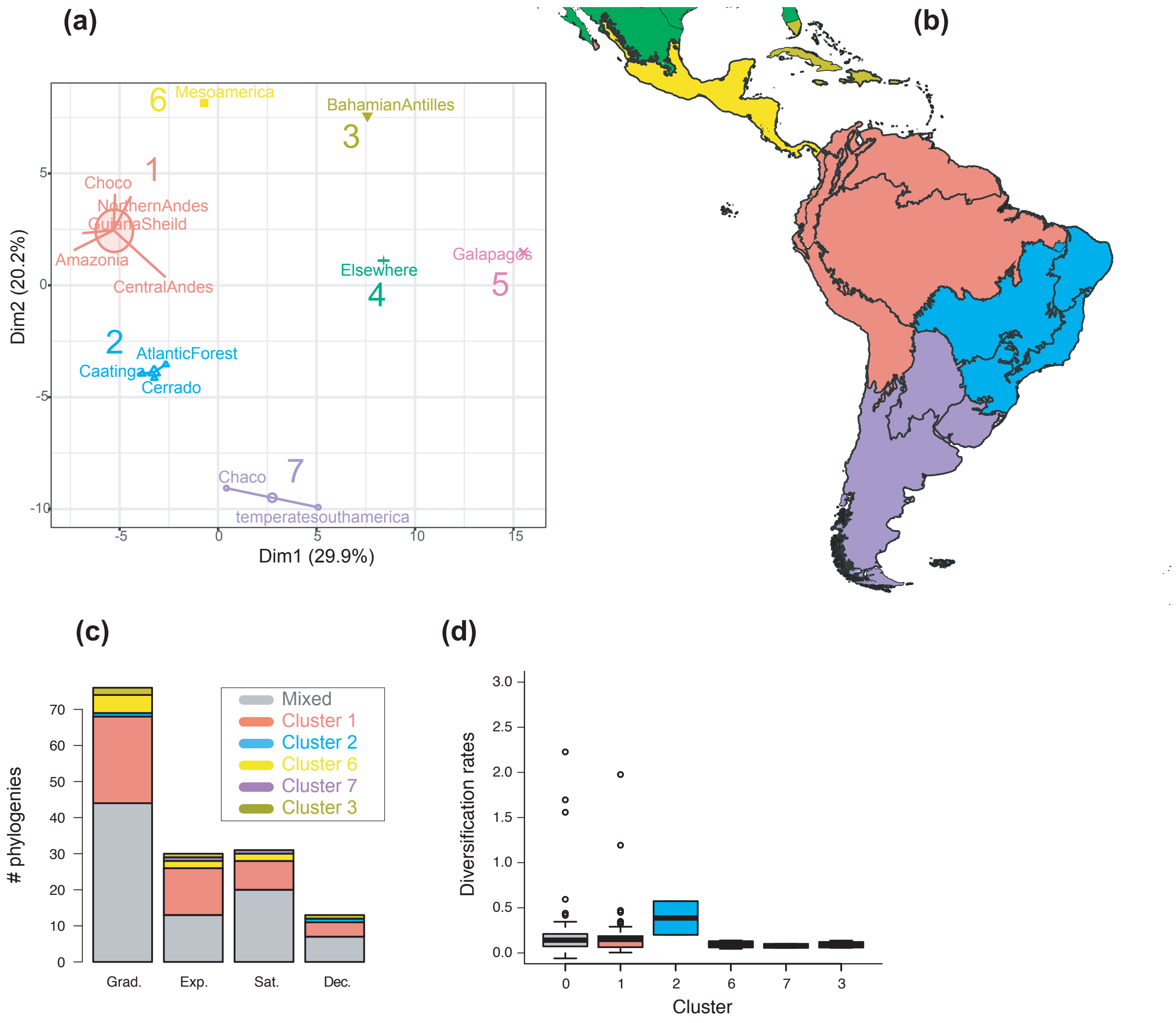

**Figure S7. The geographic structure of long-term Neotropical diversification. (a)** PCA representation of K-means clustering results for 13 areas (WWF ecoregions) and 150 clades. **(b)** Resulting clusters (1–7) on the geographic space. Colors correspond with the clusters of regions identified in (a), thick lines delineate the original ecoregions used in the analyses. **(c)** Number of phylogenies best fitting species rinchess scenarios Sc. 1–4 (Grad = Gradual increase [Sc.1], Exp = Exponential increase [Sc.2], Sat = Saturated increase [Sc.3] and Dec = Declining diversity [Sc.4]) across geographic clusters as derived from cannonical diversification rates. **(d)** Variation in diversification rates (derived from the constant-rate model) across geographic clusters.

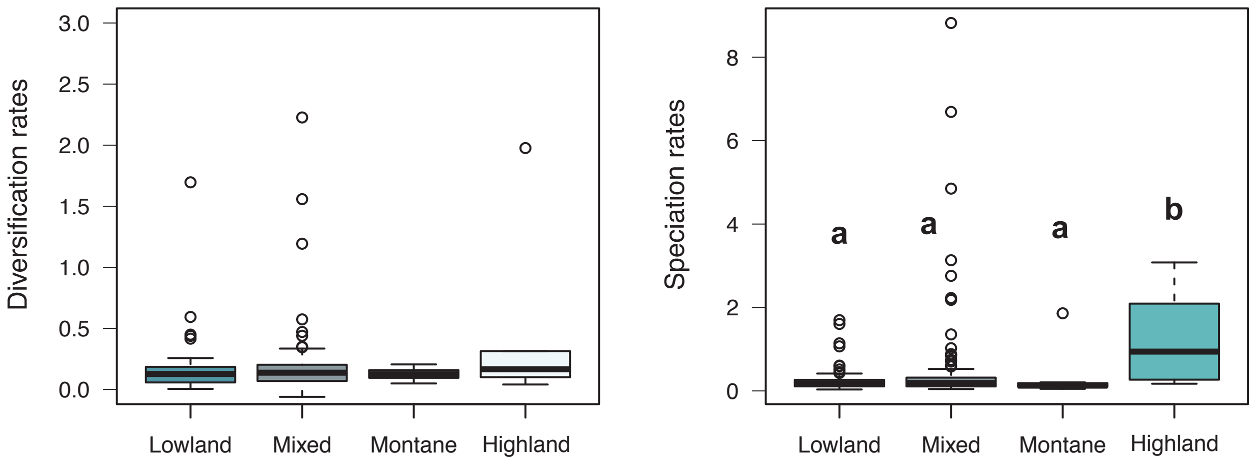

**Figure S8**. **Variation in diversification rates on 150 Neotropical phylogenies of plants and tetrapods across elevation ranges.** Diversification and speciation rates are derived from the constant-rate model (Table S6). Letters are used to denote statistically differences between groups, with groups showing significant differences in mean values denoted with different letters. This figure differs from main text Figure 6 in that the montane category has been analysed separately.

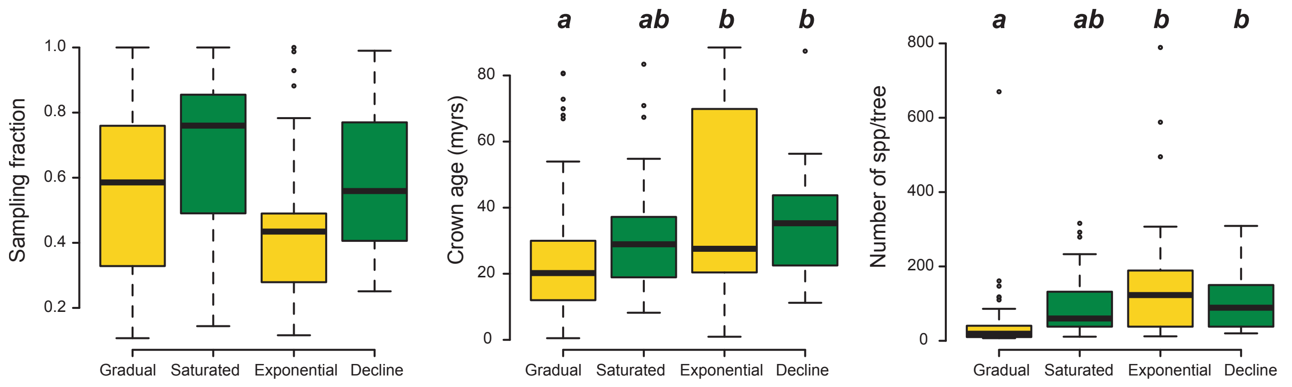

**Figure S9.** Box plots showing differences in sampling fraction, clade age (*i.e.* crown age), and number of species per tree (*i.e.* tree size) for the phylogenies supporting gradual increase (Sc. 1) vs. exponential increase (Sc. 2) vs. saturated (Sc. 3) vs. declining diversity dynamics (Sc. 4). Sampling fraction does not differ significantly between model categories, suggesting that there is no particular bias of sampling in our study. Meanwhile, there are differences in the tree size and crown age between model categories. Letters are used to denote statistically differences between groups, with groups showing significant differences in mean values denoted with different letters.

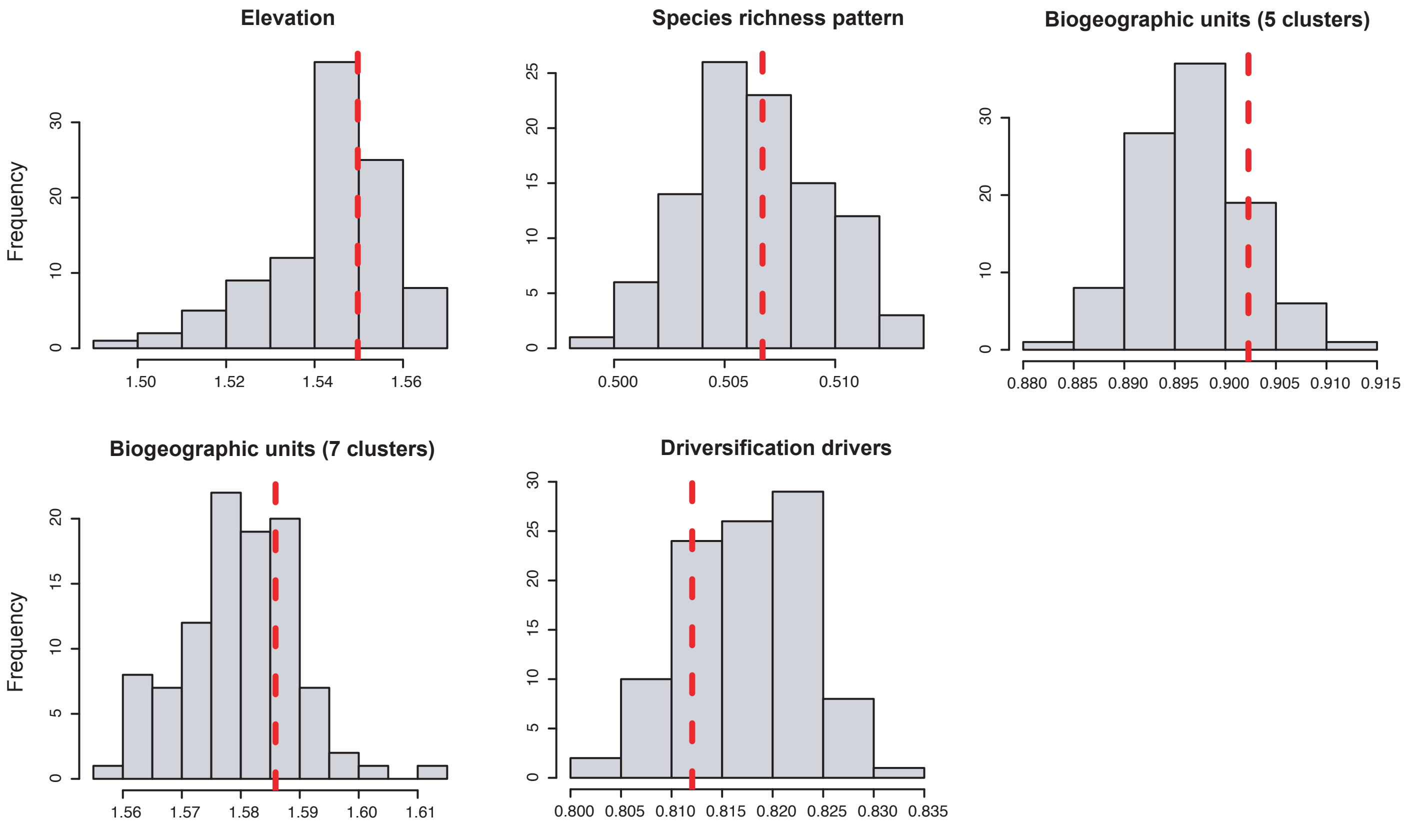

**Figure S10. Phylogenetic signal of different multicategorical traits.** Inferred δ-values (in red) compared to the distribution of values when the trait is randomised along the phylogeny.
